# Type S Non Ribosomal Peptide Synthetases for the rapid generation of tailor-made peptide libraries

**DOI:** 10.1101/2021.10.25.465728

**Authors:** Nadya Abbood, Tien Duy Vo, Jonas Watzel, Kenan A. J. Bozhueyuek, Helge B. Bode

## Abstract

Bacterial natural products in general, and non-ribosomally synthesized peptides in particular, are structurally diverse and provide us with a broad range of pharmaceutically relevant bioactivities. Yet, traditional natural product research suffers from rediscovering the same scaffolds and has been stigmatized as inefficient, time-, labour- and cost-intensive. Combinatorial chemistry, on the other hand, can produce new molecules in greater numbers, cheaper and in less time than traditional natural product discovery, but also fails to meet current medical needs due to the limited biologically relevant chemical space that can be addressed. Consequently, methods for the high throughput generation of new-to-nature natural products would offer a new approach to identifying novel bioactive chemical entities for the hit to lead phase of drug discovery programms. As a follow-up to our previously published proof-of-principle study on generating bipartite type S non-ribosomal peptide synthetases (NRPSs), we now envisaged the *de novo* generation of non-ribosomal peptides (NRPs) on an unreached scale. Using synthetic zippers, we split NRPS in up to three subunits and rapidly generated different bi- and tripartite NRPS libraries to produce 49 peptides, peptide derivatives, and *de novo* peptides at good titres up to 145 mgL^-1^. A further advantage of type S NRPSs not only is the possibility to easily expand the created libraries by re-using previously created type S NRPS, but that functions of individual domains as well as domain-domain interactions can be studied and assigned rapidly.

## Introduction

Natural products (NPs) have been used throughout the ages for the treatment of a wide range of medical conditions^1^ and still continue to be of particular importance in drug development today^2–4^. Especially bacterial NPs derived from modular megasynth(et)ases, such as non ribosomal peptides (NRPs) and polyketides (PKs), made a major contribution to modern pharmacotherapy, inter alia, for tackling infectious diseases and cancer^5^. Nevertheless, although NPs are structurally diverse and bioactive with advantagoues properties beyond Lipinski’s rule of five’^6^, like higher molecular mass and a greater molecular rigidity which can be valuable in tackling protein-protein interactions^7^, they also pose challenges for drug discovery. These challenges are mainly due to technical barriers in the identification, characterisation, isolation, screening and optimisation of natural products, which remain time, labour and cost intensive^2, 8^. As a result, and due to the lack of adequate solutions, the pharmaceutical industry withdrew from traditional natural product research. With the rapid emergence of antimicrobial resitances (AMRs)^2–4, 9–11^ and recent technological advances addressing the challenges, such as advances in cultivation^12^, DNA sequencing^13, 14, 15^, bioinformatics^16^, and synthetic biology^17^ interest in natural product research has been reignited^8^.

Despite the progress made, the complexity of NP structures make it difficult to generate synthetic NP derivatives, for example to explore structure-activity relationships and develop hits to leads. Thus, many clinical derivatives have been created by means of semi-synthesis^4^, i.e. azithromycin^18^ and cephalosporin^19^. Due to technical and chemical limitations, such modifications are often limited to a few synthetically accessible functional groups, leaving the backbone structure untouched. A commonly stressed solution to this problem is bioengineering, as it provides access to a wider range of structural diversity beyond the limitations of synthetic chemistry.^20, 21^ But as rational reprogramming efforts have been met with limited success, progress in the synthetic biology of NPs is of great importance. In this and previous work, we have therefore focused on developing tools enabling the reproducible, rapid, and simple genetic manipulation of biosynthetic gene clusters (BGCs) for the biosynthesis of NRP derivatives and even new-to-nature peptides.^22–24^

At the heart of our research are BGCs, encoding multi-functional enzymatic protein machines that enable the biosynthesis of peptides independently of the ribosome. These machines, denoted as non-ribosomal peptide synthetases (NRPSs) are in fact assembly lines, and are, inter alia, responsible for the synthesis of most antibiotic drug scaffolds in current clinical use, such as penicillin G, vancomycin, and daptomycin.^25^ In a NRPS assembly line, multiple repeating modules are responsible for selection, activation, programmed functional group modifications, and coupling of an amino acid to the growing peptide chain. An archetypal minimal module consists of three core domains: an adenylation (A) domain, which selects and activates a substrate, a thiolation (T) domain, onto which the activated amino acid is covalently attached to, and a condensation (C) domain, which catalyses peptide bond formation of the bound amino acid to the growing peptide chain. Additionally, several optional *in cis* or *in trans* acting modification domains can be present, introducing structurally complex motives into the peptide chain, for instance epimerisation, methylation, hydroxylation, and glycosylation patterns.^25^

Most recently, we developed a novel *in trans* acting synthetic type of NRPSs (type S)^22^ with reduced structural complexity compared to wildtype (WT) *in cis* acting type A NRPSs. Type S NRPSs are characterised by ‘small’ individually expressible chimeric NRPS protein subunits with attached synthetic leucine zippers, referred to as SYNZIPs (SZs)^26, 27^. Type S subunits functionally can be co-expressed and are quickly interchangeable to generate new assembly lines and peptide derivatives, respectively, in no time and with only a minimum of lab work involved. We were able to showcase how type S NRPSs can be created via splitting one protein NRPSs into two individually expressible proteins (subunits) in between exchange Unit (XU) building blocks (A-T-C tri-domain units)^22^ by leveraging the established splicing position within the C-A di-domain linker region (W][NATE)^23^ to introduce SZs. Due to the high bio-combinatorial potential of type S NRPSs and the possibility to reuse formerly cloned type S subunits in new combinations, they bear the great chance to accelerate NRPS research and NRP based early drug discovery efforts.

Here we show the potential of type S NRPSs beyond the limitations of the XU concept. We not only sought to demonstrate (I) the possibility to create a bi-partite type S NRP library by using the building blocks of only one single NRPS system, but (II) to introduce SZs within all possible NRPS linker regions. Eventually, (III) to further increase the bio-combinatoric potential of type S NRPSs, we aimed at dividing distinct BGCs into three individually expressible subunits to create tri-partite NRPS libraries.

## Results

### I. Bi-partite type S NRPS library

To create a bi-partite type S NRPS^22^ library from only one single starting NRPS, we chose the GameXPeptide synthetase (GxpS) from *Photorhabdus luminescens* TT01^28^. GxpS, which is responsible for the biosynthesis of four cyclic GameXPeptides (GXP) A-D (**1-4**), was split into all possible subunits in between individual XUs. These subunits differed in the number of XUs ranging from 1-4 XUs, both from the *N*- and *C*-terminus. In detail, we split GxpS between XUs 1 & 2, 2 & 3, 3 & 4, and 4 & 5, resulting in four initiating (subunit −1a, −1b, −1c, and −1d) and four terminating building blocks (subunit 2, −2a, −2b, −2c, and − 2d) (Fig. 1a). To functionally create type S NRPS systems, each individual initiating subunit was heterologously co-expressed in *E. coli* DH10B::mtaA^29^ together with each of the terminating subunits. With this procedure, 16 unique type S NRPSs were generated from a single WT NRPS.

**Figure 1.**
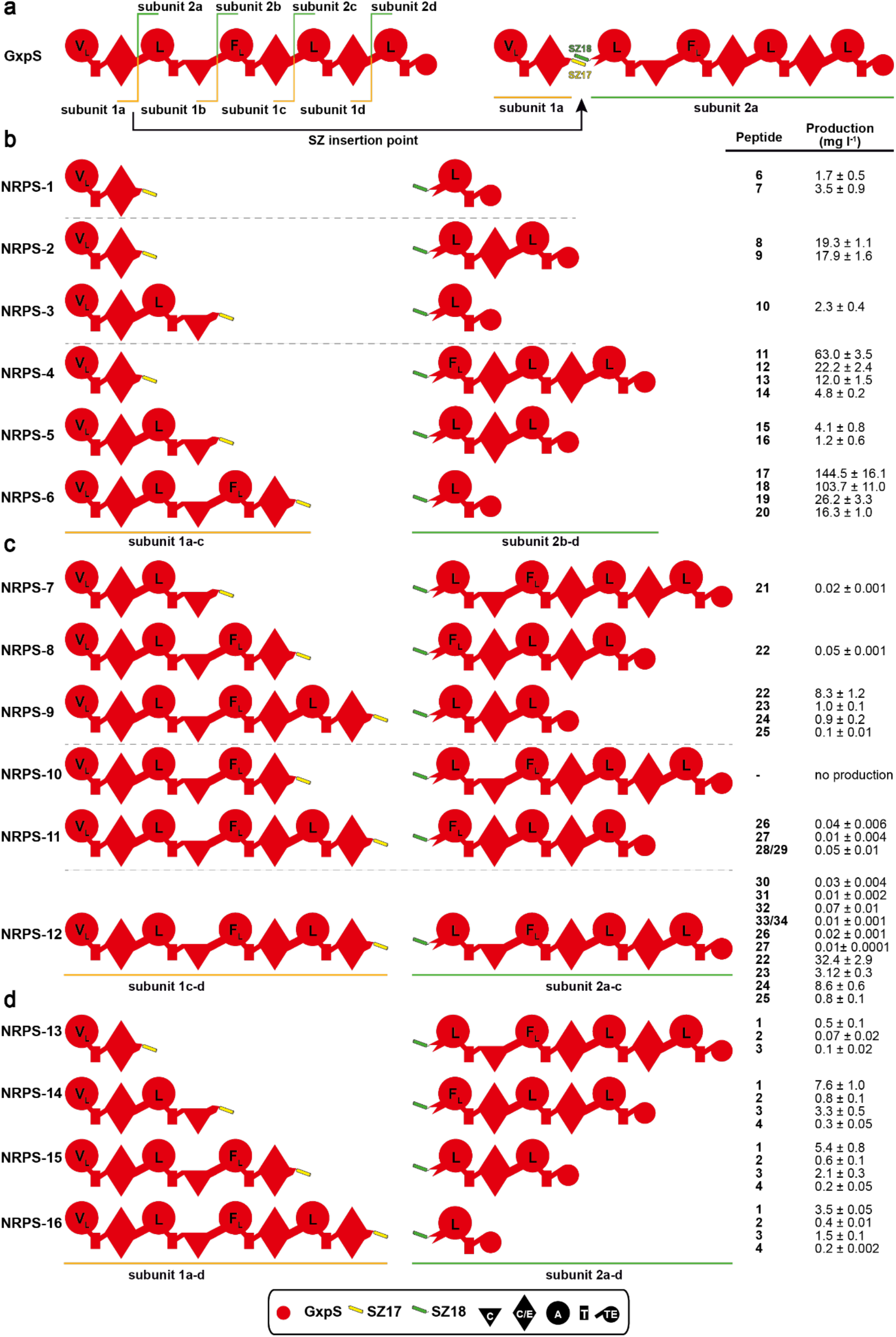
Bi-partite GxpS Library. **a** SZ 17:18 insertion between XUs 1 & 2, 2 & 3, 3 & 4 and 4 & 5 for the generation of four initiating (subunit-1a, −1b, −1c, −1d) and four terminating building blocks (subunit 2, −2a, −2b, −2c, −2d). Generated truncated type S GxpS systems (NRPS-1-6) (**b**), elongated type S GxpS systems (NRPS-7-12) (**c**), and wild type length type S GxpS systems (NRPS-7-12) (**d**) are shown. Corresponding peptide yields (mg/L) as obtained from triplicate experiments. For domain assignment the following symbols are used: A, adenylation domain, large circles; T, thiolation domain, rectangle; C, condensation domain, triangle; C/E, dual condensation/epimerization domain, diamond; TE, thioesterase domain, small circle.

Notably, HPLC-MS analysis revealed that all but one type S NRPSs (Fig. 1c, NRPS-10) showed catalytic activity, producing detectable amounts of overall 34 unique linear and cyclic peptides (**1–34**, Fig. 1) of varying length at titres up to 145 mgL^-1^ (Fig. 1). Throughout the present work, the resulting peptides (Table S1) and yields were confirmed by HPLC-MS/MS and comparison of retention times with synthetic standards (*c.f*. Supplementary Information S1-35).

An additional strength of this approach lies in in the possibility to study and characterise individual domains (*i.e*., C & TE) as well as domain-domain (*i.e*., C-A) interface interactions with respect to their substrate specificity or compatibility. An NRPS domain-domain interface is defined as the region of interdomain contact (*e.g*., A-T, T-C, C-T), involving (mostly hydrophobic) interface forming amino acid-residues at the respective domains solvent accessible surface. The interdomain interfaces are thought to form flexible and changing domain-domain contacts during the course of the catalytic cycle, which help the NRPS machinery to run and carry out catalytic reactions in an orchestrated manner^25, 30^. Traditionally, such characterisations were only done *in vitro*.^31^ However, here, the presented GxpS derived NRPS set (NRPS-1 to −16) not only enables interesting conclusions concerning the compatibility of differing C-A interface types, but also to quickly deduce the GxpS_TE-domains’ capacity to cyclise peptides differing in length from the wild type (WT) products.

At a glance, the chimeric set of 16 GxpS derived type S NRPSs consists of six truncated (NRPS-1 to −6, Fig. 1b), five elongated (NRPS-7 to −12 Fig. 1c), and five WT-length assembly-lines (NRPS-13 to 16, Fig. 1d). From the subset of truncated NRPSs 1 to 6 we detected linear di- (**6 & 7**), tri- (**8-10**), and tetra- (**11-20**) peptides, showing the expected range of derivatives due to the substrate promiscuity of GxpS_A1 (Val, Leu) and A3 (Phe, Leu). While cyclic dipeptides like diketopiperazines are found in nature^32^ or as truncated NRPs^33^ but mainly are generated by cyclodipeptide synthases using tRNA-activated amino acids^34^, the generation of tri-peptides is at least also conceivable - although we are not aware of any cyclic tri-peptides generated by NRPSs in nature but chemically synthesized examples exist^35^. Cyclic tetra-peptides are common and numerous NRPs-based examples exist (e.g. fungisporin^36^, tentoxin^37^, azumamides A-E^38^, xenotetrapeptide^39^). However, according to our data, the GxpS-TE does not seem to be able to cyclise the synthesised tri- and tetra-peptides, suggesting that a length of five consecutive building blocks is its lower limit for cyclisation (NRPS-13 to −16).

Looking at the upper boundary, NRPS-7 to −9 were able to produce and cyclise hexapeptides (**21-25**) and NRPS-11 even synthesised cyclic hepta-peptides (**27-29**). However, the TE-domain seems to have reached its capacity to efficiently cyclise peptides at a length of 7 consecutive amino acid building blocks. Although we also detected cyclic peptides (**22-27**) from production cultures of the 8-modular NRPS-12, these cyclic peptides are presumably autocatalytically cyclised shunt products of enzyme-bound hexa- and heptapeptides. NRPS-12 only was capable to hydrolyse the synthesised octapeptides (**30-34**), resulting in linear octapeptide derivatives (Fig. 1d).

Overall, through this simple and quick experimental procedure, we found that the TE domain of GxpS is quite versatile, accepting a range of peptides from two to at least eight building blocks, but is only able to effectively catalyse cyclisation within a narrow range of five to seven building blocks. In turn, information about TE domains’ substrate specificities and preferences, respectively, gained via generating a series of type S NRPSs will help to guide future engineering projects in identifying suitable termination domains. This is of particular interest when it comes to large scale NRPS engineering campaigns, as TE-domains that are less prone to a peptide’s amino acid sequence and length are more versatile and can help to overcome TE-domain limitations of artificial NRPS systems. The approach shown is not only superior to *in vitro* characterisation in terms of workload, but also much cheaper (no SNAC peptides need to be synthesised), more robust (no spontaneous or autocatalytic side effects), scalable, can be performed in high throughput, and does not suffer from the *in vitro* bias known for excised NRPS proteins as recently described.^40^

Type S NRPSs can also be used to study C domain specificities and C-A interface compatibilities. Since in the case of GxpS (NRPS-1 to −16), the respective activated and incorporated amino acids are too similar to each other to draw valuable conclusions on C domain specificities-but this was shown previously^22, 40^ – we focused on characterising the compatibility of different C-A interfaces with each other. In general, based on the particular reactions C domains catalyse (condensation or epimerisation and condensation), their donor-T domain bound substrate range (acetyl-, formyl-, D- or L-amino acids), acceptor-T domain bound substrate range (L-amino acids, amines) and from phylogenetic reconstructions, they are categorised into five different groups: ^L^C_L_, ^D^C_L_, dualC (C/E), C_start_, and C_term_.^25^ However, in our example the C-A interface depends on whether there is a C or C/E domain directly upstream of the A domain resulting in C-A or C/E-A type interfaces – as evidenced by phylogentic analysis (Fig S36^41^).

In brief, depending on whether or not the interface type naturally occurring at a specific site has been altered, we observed major differences in the production titres between NRPSs of similar length and amino acid composition, suggesting that C-A interface types indeed play an important role when it comes to (re-)designing NRPSs. For instance, the four-modular type S NRPSs (NRPS-4 to −6, Fig. 1b) produced tetra-peptides (**11-20**) with yields varying widely, ranging from 4.1 mg/L to 144.5 mg/L. As could be expected, the best producing system, NRPS-6 (144.5 mgL^-1^), has the same interface type (C/E-A) and an interface most similar to that of the WT C5-A5 interface (Identity 89.3 %, supplementary information Table S5). For NRPS-4, in which the interface type changed from C-A to C/E-A, we observed high (63 mg/L) but still significant lower yields than for NRPS-6. Eventually, expression of NRPS-4 resulted in the lowest titre (4.1 mg/L), indicating that changes from C to C/E have a greater impact on production than changes from C/E to C, since here the interface type changed from C/E-A to C-A. These observations further are supported by NRPS-2 and −3 and by NRPS-7 to −9. NRPS-2 and −3 synthesised **8, 9** and **10**. While NRPS-2 with the same interface type as the WT produced **8** and **9** at titres of 19 mg/L and 18 mg/L, respectively (Fig. 1b), the switch from C/E-A type to C-A type in NRPS-3 resulted in a sharp drop in production of **10** to 2.3 mg/L. Accordingly, comparing the titres of the hexa-peptide (**21-25**) producing NRPSs (NRPS-7 to −9), the WT-like C-A interface harbouring NRPS-9 showed ~40 to ~20-fold higher titres than NRPS-7 and −8, respectively (Fig. 1c).

In combination with the most recently published extended gatekeeping function, describing the influence of C domains and the particular formed C-A interface on the catalytic activity and substrate selectivity of A domains^40^, this data helps us to refine the NRPS design principles published previously even further. We assume that altering the C domain type directly upstream of an A domain of interest substantially impairs C-A didomain contacts, resulting in reduced catalytic activity of the A domain and therefore overall productivity of the respective NRPS protein. In retrospect, this might also explain why some of our previously published recombinant NRPS systems showed reduced production titres - while others showed no impairment or even increased catalytic activity.^22, 23^

### II. Other SZ insertion sites

To further explore and extend the applicability of SZs for the construction of type S NRPSs beyond the borders of the XU concept^23^, we decided to introduce SZ17:18 within the T-C (NRPS-16) and A-T (NRPS-17) linker regions of the cyclic xenotetrapeptide (**5**; cyclo(vLvV)) producing synthetase (XtpS) from *Xenorhabdus nematophila* ATCC 19061^39^ (Fig. S37) – and compared resulting peptide yields with WT XtpS and NRPS-15 (Fig. 2), which was constructed previously.^22^ Both, NRPS-16 and −17 synthesised **5** at ~86% and ~280% compared to WT XtpS and NRPS-15, respectively.

**Figure 2.**
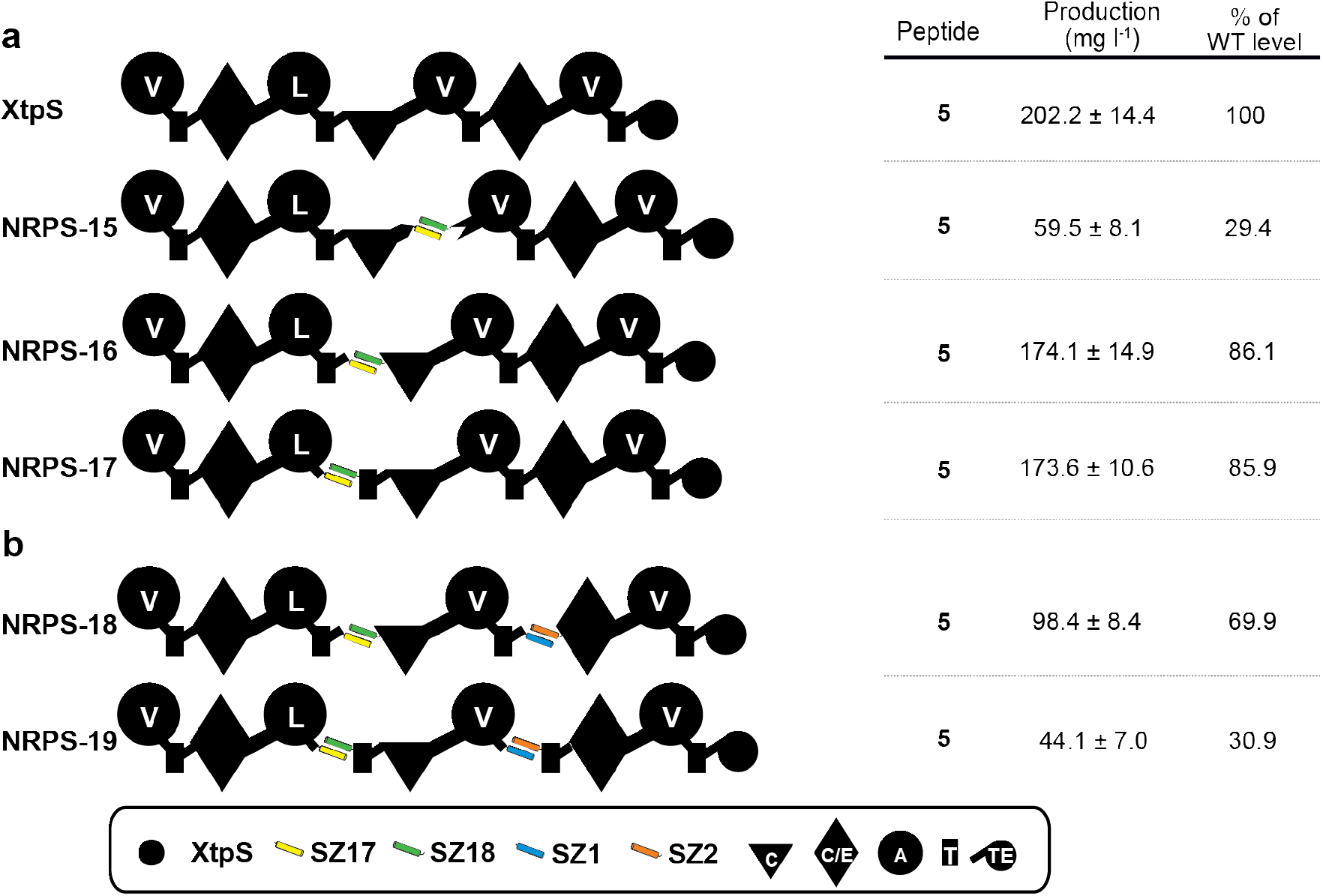
Other splicing positions and Tri-partite type S XtpS. **a** SZ17:18 insertion within the C-A (NRPS-17), T-C (NRPS-18) and A-T (NRPS-19) linker (**a**) XtpS split into three subunits by the insertion of the SZ1:2 pair. SZ17:18 and SZ1:2 pairs were inserted within T-C (NRPS-20) and A-T (NRPS-21) (**b**). Corresponding peptide yields (in mg/L) are determined from triplicate experiments.

While the catalytic activity of NRPS-16 was not surprising, as the introduced SZs are mimicking natural DDs^42^, the observed good activity of NRPS-17 was unexpected. During a catalytic cycle of a module, especially the A-T interaction is considered as highly dynamic. After the adenylation reaction, the A_sub_-domain must fulfil a torsion of 140° in respect to the A_core_-domain such that the *holo*-T-domain can meet the distance to the activated amino acid (thiolation reaction).^25^ Thus, it was assumed that the additional rigidity, inserted by the structured α-helical amino acid stretches of the SYNZIPs, would result in loss of function. The recently gathered structural data of large constructs of the linear gramicidin synthesising NRPS (LgrA)^43^, might serve as an explanation for the observed activity. There a very high structural flexibility was reported, potentially bringing closely together domains that are far apart in protein sequence and therefore facilitating synthetic cycles with inserted tailoring domains, unusual domain arrangements like A-C-T^44^, module skipping^45^, and presumably also SZs.

More dipartite type S NRPSs (NRPS-39 to −44), split in between (T][C) and within modules (A][T) as well as within C domains (C_Dsub_][CA_sub_) are depicted in Fig. S38 and S39.

### III. Tri-partite type S NRPS library

The potential of bi-partite type S NRPSs to generate bio-combinatoric libraries from a small set of NRPS subunits was shown previously^22^ and above (Fig. 1 & 2a). But, the bio-combinatoric potential could further be increased if it were possible to split NRPS systems into three or more subunits (*c.f*. Supplementary Information S40 and S41).

For a first proof of concept to functionally split NRPSs into three subunits, we again targeted XtpS and inserted, next to SZ17:18, a second SZ pair (SZ1:2) into both, NRPS-16 and −17, to build an orthogonal interaction network (Fig. 2b). The resulting tri-partite type S NRPSs −18 and −19 are split in between modules 2-3 & 3-4 (NRPS-18) and within the A-T linker regions of modules 2 & 3 (NRPS-19), respectively (Fig. 2b). Both, NRPS-18 and −19, produced **5** with 69.9% and 30.9% compared to WT XtpS but also with decreased yields compared to their bipartite counterparts (NRPS-16 & −17). In addition to the cumulative effect of inserted impairments, caused by a higher degree of engineering, we assume that SZ1:2 also contributed to the reduced production titre of **5** since the SZ1:2 pair is significantly longer than SZ17:18 and probably disturbs catalytic reactions of the tri-partite type S XtpSs by the inserted additional rigidity. Although NRPS-18 produced **5** at slightly higher titres than NRPS-19, in a next step we decided to use the A-T splicing position (NRPS-17 & −19) for the construction of a small but diverse tri-partite NRPS library with subunits derived from various *Photorhabdus* and *Xenorhabdus* strains, because T-C-A tri-domains as a catalytically active unit to reprogram NRPSs are underrepresented.

Eleven NRPS subunits with attached SZs were extracted from five different BGCs, namely from GxpS, XtpS as well as from the gargantuanin (GarS), xenolindicin (XldS)^29^, and the szentiamide (SzeS)^46^ producing synthetases. Overall 18 (NRPS-21 to −38) from 45 possible co-expressions of three plasmids each yielded detectable amounts (0.1 – 38 mg l^-1^) of 18 different peptides, 13 of which were new (Fig. 3, Figs. S24-S40). Despite the method’s general simplicity, the overall efficacy or recombination potential of T-C-A units compared to XUs appears to be slightly more restricted^22^. For example, neither coexpression of all type S subunits to reconstitute SzeS, nor any combination involving the Ser and Thr specifying subunits from XldS and GarS, yielded any detectable peptide, respectively. These results probably indicate an incompatibility of formed chimeric A-T interfaces or substrate incompatibilities at the respective C domains donor site. Yet, in light of previous results concerning C domain specificities^31, 40, 47–49^, the latter seems to be unlikely. Especially as we were not even able to reconstitute catalytic activity of the tripartite SzeS, we concluded that the respective subunits might have lost their functionality. Due to the sequence and structural flexibility of the targeted A-T linker regions, key interactions within protein-protein interfaces that must be maintained are hard to predict. Therefore, it is likely that the insertion of SZ pairs structurally affected these subunits, resulting in a loss of function or their ability to ‘communicate’ with downstream subunits – as was already expected for NRPS-19 (Fig. 2b).

**Figure 3.**
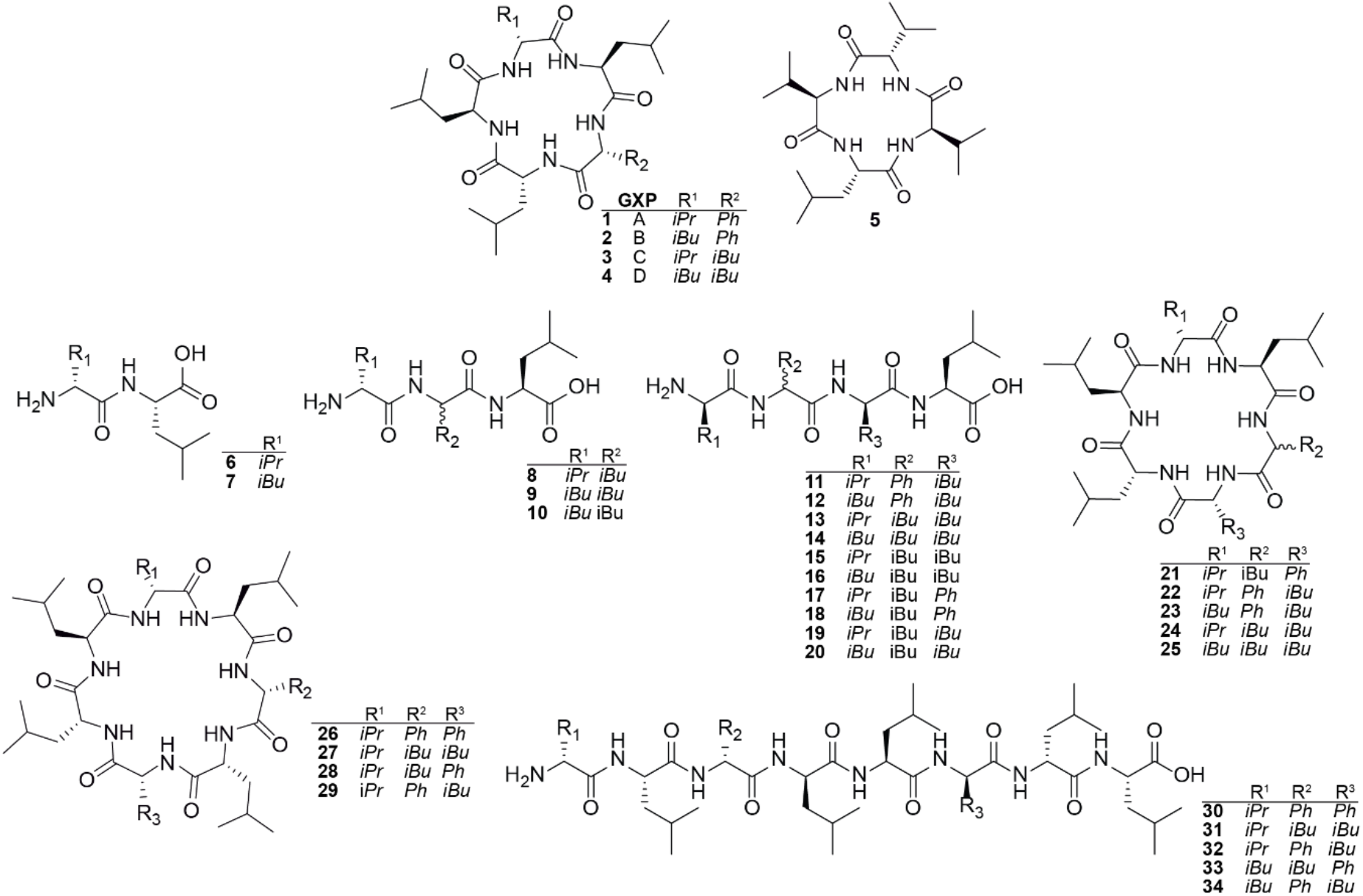
Structures of produced compounds. GameXPeptide A-D (**1-4**), xenotetrapeptide (**5**) and GXP derivatives (**6-34**) are depicted.

**Figure 4.**
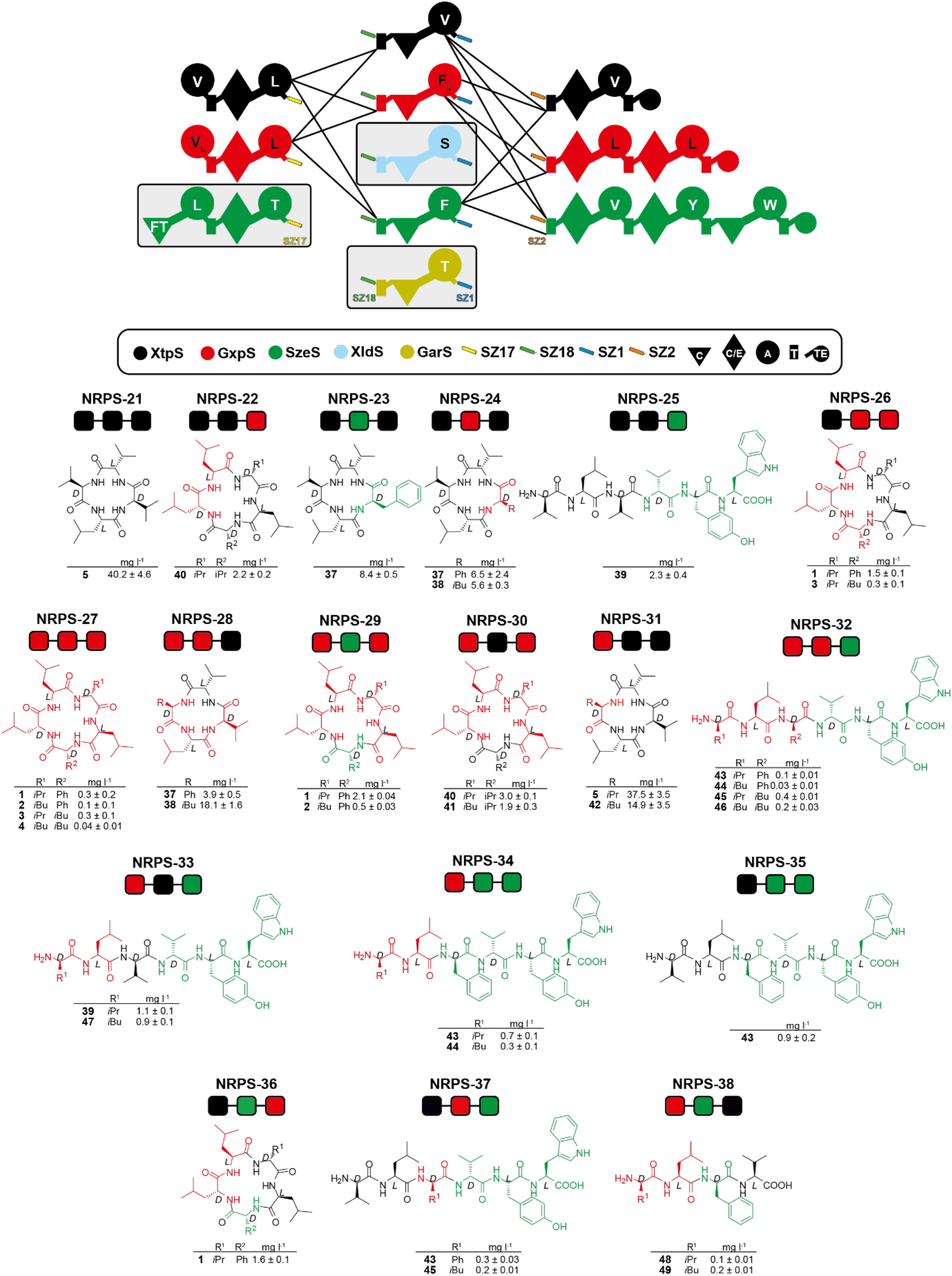
Tri-partite type S NRPS library. The upper section, generated building blocks, depicted in the symbol assignment as introduced previously, are illustrated. Solid lines represent functional combinations. In the lower section, building blocks were simplified and illustrated as boxes representing subunit 1 to 3. From 11 generated and 8 functional building blocks, a total of 18 type S NRPS were confirmed as functional by HPLC-MS. Corresponding peptide yields (mg/L) were obtained from triplicate experiments.

Furthermore, for some tri-partite NRPSs (NRPS-32, −34, −35, −37 and −38), we were able to detect peptides (**43-46, 48**, and **49**) only in very low amounts, which might be explained by the aforementioned impairment within the A-T domain interface and/or the mere length of the chosen SZ1:2 pair. Taking these points into consideration, we assume that productivity can significantly be increased when another fusion site is chosen, another SZ pair is chosen, or, if possible, SZ1:2 is truncated. Nevertheless, the quantity of produced peptides should not distract from the strength of this method: *The ability to generate an enormous variety of new-to-nature NRPSs in an unprecedented short time and with a minimum of lab work involved*. The goal is to produce a large variety of NRPs to generate structural diversity rather than to produce peptides on an industrially relevant scale. This can be focused on in a second step, when the most interesting candidates have been found.

## Conclusion

Combinatorial chemistry can generate potential new compounds in a greater number, cheaper and in less time than the traditional NP discovery process^4, 50^. Yet, a lot of large chemical screening collections have failed in practice, because they do not exhibit enough diversity within the biologically relevant chemical space^2, 51^. Nevertheless, although ‘NP-derived’ libraries (scaffold is identical to the NP scaffold) and ‘NP-inspired’ libraries^52^ (scaffold is closely related to NP scaffolds) have proven to be enriched in bioactivity over typical collections of synthetic compounds, NP discovery has strongly decreased over the past 20 years^2, 53–56^ In part because of extraction and supply issues^57^, and the frequent (re-) discovery of natural products with structural precedent in the literature – structurally unique scaffolds represent a decreasing percentage of the total number of isolated NPs. For NRPs, this becomes quite clear from the example of the Norine database^58^. It is dedicated to collect all discovered non-ribosomally synthesised peptides, currently containing ~1800 peptides, of which ~800 peptides are characterised, but only ~175 scaffolds are unique. Interestingly, from the characterised set of peptides already more than 500 different NRP building blocks are known^59, 60^, spanning a combinatorial chemical space for novel peptides beyond the total number of molecules in the whole universe and comprehension of the human mind.

Consequently, splitting known BGCs into individually expressible subunits and to recombine them simply by attaching SZs and co-expressing a variety of unrelated BGC subunits in a high throughput manner puts us in a position to easily enlarge the known structural diversity, and to outcompete both, traditional natural product research and combinatorial synthetic chemistry approaches. This especially might be true for the generation of type S NRPSs to discover novel antimicrobials because in this particular case, the high throughput generation and production of antimicrobial peptides can be coupled to bioactivity testing against human pathogens, *i.e*., via nanoFleming^61^, a miniaturized and parallelized high-throughput inhibition assay.

With this study we sought to showcase that, although nature still bears an enormous variety of natural products only waiting to be discovered^62^, methods for the identification and characterisation of new scaffolds from nature are still far from surpassing the throughput of new chemical entities that can be generated from type S NRPSs. Imagine, that you want to create a library of 10^6^ NRP variants. According to the current state of the art (pre-type S), this would require the design and construction of 1.000.000 NRPSs, which of course is not feasible. By applying bi-partite type S NRPSs one can reduce the workload to 2000 type S NRPSs building blocks - 1000 building blocks for each subunit. This in turn implies that only 100 building blocks per subunits, or 300 in total, must be created if common one-protein type A NRPSs are split into three individually expressible subunits (100 subunits A x 100 subunits B x 100 subunits C = 10^6^ combinations). Even if some type S systems are non-functional or do not synthesise the desired products, either due to the incompatibility of interface types, C and TE domain specificity issues, or capacity limitations of TE domains (Fig. 1), these limitations are no longer a significant problem, as sufficient diversity is still produced. Especially along with our previously developed design principles^22–24^, we have created a powerful toolset to predictably engineer new-to-nature NRPSs.

The striking advantage of type S NRPSs, however, is that once generated libraries can be enlarged and extended continuously and at any time, since generated subunits are not covalently linked. Moreover, a great variety of type S NRPSs can already be achieved from a small set of BGCs, as exemplified by turning GxpS into 16 artificial NRPSs (Fig. 1). The bi-partite type S GxpS library (Fig. 1), although not intended, led to the discovery of a further puzzle piece complementing our previously established NRPS design principles^20, 23, 24, 40^ – the importance of different C-A interface types, which have never been considered before for the construction of un-impaired high yielding artificial NRPSs.

## Acknowledgements

This work was supported by the LOEWE research cluster MegaSyn, the LOEWE center TBG, both funded by the state of Hesse and an ERC advanced grant (grant agreement number 835108).

