## Supplementary Information for "Type S Non Ribosomal Peptide Synthetases for the rapid generation of tailor-made peptide libraries"

Figure S35. HPLC/MS data refers to Figure 3 (NRPS-38) of compounds 46 and 47 produced in *E. coli* DH10B::*mtaA*. 42
Figure S36. Phylogenetic three of extracted C-A and C/E-A didomains from GxpS. 42
Figure S37. Sequence alignments of XtpS linker sequences. 43
Figure S38. Additional examples of bipartite type S NRPS split in between and within RtpS modules[1]. 44
Figure S39. Additional examples of bipartite type S NRPS split within modules of xefoampeptide-producing NRPS (XfpS)[8] and between XfpS and XtpS. 44
Figure S40. Extended comparison of workflows to generate chimeric NRPSs. 45
Figure S41. Comparing the throughput between di- and tripartite NRPSs.. 46
Figure S42. HPLC/MS data refers to Supplementary Figure 18 (NRPS-39) of compounds 48, 49, 50 and 51 produced in *E. coli* DH10B::*mtaA*. 47
Figure S43. HPLC/MS data refers to Supplementary Figure 18 (NRPS-40) of compound 55 produced in *E. coli* DH10B::*mtaA*. 47
Figure S44. HPLC/MS data refers to Supplementary Figure 19 (NRPS-43) of compounds 52 and 53 produced in *E. coli* DH10B::*mtaA*. 48
Figure S45. HPLC/MS data refers to Supplementary Figure 19 (NRPS-44) of compound 54 produced in *E. coli* DH10B::*mtaA*. 48
**42. Reference 48**

### 1. Material and Methods

#### 1.1 Cultivation of Strains

All *E. coli*, *Photorhabdus* and *Xenorhabdus* strains were cultivated in LB (10 g/L Trypton, 5 g/L yeast extract, 10 g/L NaCl, pH 7,5) or TB liquid medium (12 g/L Tryton, 24 g/L yeast extract, 0.4% (v/v) Glycerin, 10% (v/v), 17 mM KH2PO4, 72 mM K2HPO4, pH 6.5) at 37°C (*E. coli*) or 30°C (*Photorhabdus*, *Xenorhabdus*) for 16-18 h at 160-200 rpm. 1% (w/v) agar was added for growth on solid LB. If necessary, medium was supplemented 1:1000 with kanamycin- (50 µg/mL in sterile ddH2O), chloramphenicol- (34 µg/mL in ethanol) and/or spectinomycin stock solution (50 mg/mL in sterile ddH2O). For short-time storage LB agar plates were stored either at 4°C (*E. coli*) or 18°C (*Photorhabdus*, *Xenorhabdus*). For permanent storage, liquid cultures were supplemented with 20% (v/v) glycerol and frozen at -80°C.

#### 1.2 Cloning of biosynthetic gene clusters

##### 1.2.1 Bi-partite plasmid library

Basic vectors pCK_0402 and pCOLA_ara/tacI containing SZs 17 and 18 (pENTR-SYNZIP17/18 were a gift from Amy Keating, Addgene plasmids #80671/80672; RRID:Addgene_80671/80672) were already construed before. In brief, cloning (of both vectors) was as follow: primers KB_pACYC_FW/ KB_pACYC_RV and KB_pCOLA_FW/ KB_pCOLA_FW were used to insert SZ encoding sequences of SZ17 and 18 into plasmids pCK_0402 and pCOLA_ara/tacI, respectively. Obtained pCK_0402_SZ17 and pCOLA_ara/tacI_SZ18 were linearized using primers KB_pACYC_II_FW/ KB_pACYC_II_RV and KB_pCOLA_II_FW/KB_pCOLA_II_RV. Plasmids pJW75 and pJW76 were already constructed before and remaining plasmids pNA104-pNA109 were generated by amplifying GxpS subunits 1a, 1c, 1d and 2a, 2c, 2d and cloning building blocks into plasmid BBs and pCK_0402_SZ17 and pCOLA_ara/tacI_SZ18, respectively. Therefore, genomic DNA (gDNA) from *Photorhabuds luminescens* TT01 was isolated using the Qiagen Gentra Puregene Yeast/Bact kit and subunits were amplified by applying primers (na194/na208 (pNA104), na209/na56 (pNA105), na194/na210 (pNA106), na211/na56 (pNA107), na194/na212 (pNA108), na213/na56 (pNA109)) obtained from eurofins genomics containing homologous ends to the plasmid backbones. PCR products were purified using Spin DNA Extraction-Kit (Stratec) and DNA fragments were assembled into linearized BBs by either HotFusion or HiFi cloning. Obtained plasmids were verified by plasmid digest using type II restriction endonucleases (specifically cutting the insert sequence). Correctly, verified plasmids were additionally sanger sequenced by eurofins genomics. Plasmids pNA104, pJW75, pNA106 or pNA108 (containing subunits 1 fragments) were co-expressed with pNA105, pJW76, pNA107 or pNA109 (containing subunit 2 fragments) into *E. coli*::mtaA in different combinations for the construction of GxpS derivatives.

##### 1.2.2 Other splicing positions

pJW61 and pJW62 were already constructed. Splicing positions within the T-C and A-T linker were determined by aligning XtpS $T_2$-$C_3$/$T_3$-C/$E_4$ and $A_2$-$T_2$/$A_3$-$T_3$ linker sequences with linker regions excised from Dhbf crystal structure (Protein Database ID: 5U89) to draw conclusions on secondary structures. Splicing positions were depicted within the sequence motives RV/LP (T-C linker) and VY/AAP (A-T linker) and are illustrated in Fig. S19. For the construction of plasmids pNA2, pNA3 and pNA4, pNA5, XtpS TCA and ATC building blocks were obtained from *Xenorhabdus nematophila* gDNA and building block were cloned into pCOLA_ara/tacI and pCK_0402. Verification of correctly assembled plasmids was done as described before. pNA2, pNA3 and pNA4, pNA5 were co-expressed into *E. coli*::mtaA, respectively.
For the construction of tripartite XtpS systems a second SZ pair was introduced. Therefore, SZs 1 (pQLinkHD-SYNZIP1 was a gift from Amy Keating, Addgene plasmid #80647;

RRID:Addgene_80647) and 2 (pQLinkHD-SYNZIP2 was a gift from Amy Keating, Addgene plasmid #80658; RRID:Addgene_80658) were inserted into the $T_3$-C/$E_4$ and $A_3$-$T_3$ linker and plasmids pNA15/pNA16 (A-T split) and pNA17/pNA18 (T-C split) were constructed. Primers na28/na30 (A-T and T-C split) and na34/na32 (A-T and T-C split) were used to insert the first half of SZ1 and 2 into pCOLA_ara/tacI_SZ18 and pCDF_ara/tacI, respectively. In a second polymerase chain reaction (PCR), the second half of both SZs was introduced by applying primers na29/na30 (A-T and T-C split) and na34/na33 (A-T and T-C split) to obtain pCOLA_ara/tacI_SZ18_SZ1 and pCDF_ara/tacI_SZ2. Plasmids pNA4, pNA15, pNA16 (A-T split) and pNA2, pNA17, pNA18 (T-C split) were transformed into *E. coli*::*mtaA* respectively to reconstitute XtpS.

##### 1.2.2 Tri-partite NRPS library

ATC building blocks for the construction of plasmids pNA24, pNA26, pNA27, pNA28, pNA29, pNA30, pNA31 and pNA35 were obtained from *Photorhabdus luminescens*, *Xenorhabdus szentirmaii* and *Xenorhabdus bovenii* gDNA. Subunits were cloned into pCK_0402_SZ17 (pNA26, pNA29), pCOLA_ara/tacI_SZ18_SZ1 (pNA24, pNA27, pNA30 and pNA35) and pCDF_ara/tacI_SZ2 (pNA28, pNA31) using HotFusion cloning. Verification of correctly assembled plasmids was done as described before. Plasmids pNA4, pNA26 or pNA29 (containing subunits 1 fragments) were co-expressed with pNA15, pNA24, pNA27, pNA30 or pNA35 (containing subunit 2 fragments) and pNA16, pNA28 or pNA31 (containing subunit 3 fragments) into *E. coli*::mtaA in different combinations for the construction of artificial NRPS systems.

#### 1.3 Heterologous expression of NRPS templates and HPLC-MS analysis

After plasmid transformation into *E. coli DH10B*::mtaA, cells were grown overnight in LB medium containing all necessary antibiotics (50 µg/ml kanamycin, 34 µg/ml chloramphenicol, 50 µg/ml spectinomycin). 100 µl of an overnight culture were used to inoculate 10 ml LB medium containing antibiotics, 0.002 mg/mL L-arabinose and 2 % (v/v) XAD-16. After incubation for 72 h at 22 °C, XAD-16 beads were harvested and one culture volume methanol was added. Methanol extraction was conducted for 60 min at 22 °C. The organic phase was filtrated, cleared extract were centrifuged for 20 min at 17000g. and methanol extracts were used for LC-MS analysis. Liquid chromatography was performed on an UltiMate 3000 LC system (Dionex) with an installed C18 column (ACQUITY UPLCTM BEH, 130 Å, 2.1 mm x 100 mm, Waters). Separation was conducted at a flow rate of 0.4 mL/min using acetonitrile (ANC) and water containing 0.1% fromic acid (v/v) in a 5-95% gradient over 16 min. Mass spectrometric analyses were performed using an ESI iontrap mass spectrometer (AmaZon X, Bruker). ESI-MS spectra were recorded in the positive-ion-mode with the mass range from 100-1200 *m/z* and ultraviolet (UV) at 200-600 nm. Evaluation was performed using DataAnalysis 4.3 software (Bruker).

##### 1.4 Peptide quantification

The absolute production titers of were calculated as previously described.[1] Synthetic standards **6** (for quantification of **6** and **7**), **8** (for quantification of **8** and **9**), **10** (for quantification of **10**), **11** (for quantification of **11-14**), **15** (for quantification of **15** and **16**), **17** (for quantification of **17-20**), **21** (for quantification of **21**), **22** (for quantification of **22-25**), **26** (for quantification of **26-29**), **32** (for quantification of **30-34**), **1** (for quantification of **1-4** and **38-39**) and **5** (for quantification of **5, 35, 36** and **40**) were synthesized in house. **37** (for quantification of **37** and **45**) and **41** (for quantification of **41-44** and **40**) were obtained by Synpeptide.

##### 1.5 Chemical synthesis

Chemical synthesis of all peptides was performed as described previously.[2]

#### 2. Supplementary tables

**Table S1.** ESI-MS data of all produced peptides. The red labeled AAs show a different configuration than expected probably due to a non-functional epimerization domain

| Peptide (#) | Theoretical mass-to-charge ratio (*m/z*) | Molecular formula | AA sequence | Reference |
|---|---|---|---|---|
| **1** | 586.40 | $C_{32}H_{51}O_5N_5$ | cyclo(vfLlL) | [3] |
| **2** | 600.41 | $C_{33}H_{53}O_5N_5$ | cyclo(lfLlL) | [3] |
| **3** | 552.41 | $C_{29}H_{53}O_5N_5$ | cyclo(vlLlL) | [3] |
| **4** | 566.43 | $C_{30}H_{55}O_5N_5$ | cyclo(llLlL) | [3] |
| **5** | 411.29 | $C_{21}H_{38}O_4N_4$ | vLvV | [4] |
| **6** | 231.17 | $C_{11}H_{22}O_3N_2$ | vL | this study |
| **7** | 245.19 | $C_{12}H_{24}O_3N_2$ | lL | this study |
| **8** | 344.25 | $C_{17}H_{33}O_4N_3$ | vlL | this study |

| | | | | |
|---|---|---|---|---|
| **9** | 358.28 | $C_{18}H_{35}O_4N_3$ | lIL | this study |
| **10** | 358.28 | $C_{18}H_{35}O_4N_3$ | lLL | this study |
| **11** | 491.32 | $C_{26}H_{42}O_5N_4$ | vfIL | this study |
| **12** | 505.34 | $C_{27}H_{44}O_5N_4$ | lfIL | this study |
| **13** | 457.34 | $C_{23}H_{44}O_5N_4$ | vIIL | this study |
| **14** | 471.35 | $C_{24}H_{46}O_5N_4$ | lIIL | this study |
| **15** | 457.34 | $C_{23}H_{44}O_5N_4$ | vLLL | this study |
| **16** | 471.35 | $C_{24}H_{46}O_5N_4$ | lLLL | this study |
| **17** | 491.32 | $C_{26}H_{42}O_5N_4$ | vLfL | this study |
| **18** | 505.34 | $C_{27}H_{44}O_5N_4$ | lLfL | this study |
| **19** | 457.34 | $C_{23}H_{44}O_5N_4$ | vLIL | this study |
| **20** | 471.35 | $C_{24}H_{46}O_5N_4$ | lLIL | this study |
| **21** | 699.48 | $C_{38}H_{62}O_6N_6$ | cyclo(vLLfIL) | this study |
| **22** | 699.48 | $C_{38}H_{62}O_6N_6$ | cyclo(vLfIIL) | this study |
| **23** | 713.50 | $C_{39}H_{64}O_6N_6$ | cyclo(lLfIIL) | this study |
| **24** | 665.50 | $C_{35}H_{64}O_6N_6$ | cyclo(vLIIIL) | this study |
| **25** | 679.51 | $C_{36}H_{66}O_6N_6$ | cyclo(lLIIIL) | this study |
| **26** | 846.60 | $C_{47}H_{71}O_7N_7$ | cyclo(vLfIfIL) | this study |
| **27** | 778.58 | $C_{41}H_{75}O_7N_7$ | cyclo(vLIIIIL) | this study |
| **28** | 812.56 | $C_{44}H_{73}O_7N_7$ | cylco(vLfIIIL) | this study |
| **29** | 812.56 | $C_{44}H_{73}O_7N_7$ | cyclo(vLIIfIL) | this study |
| **30** | 977.64 | $C_{53}H_{84}O_9N_8$ | vLfILfIL | this study |
| **31** | 909.67 | $C_{47}H_{88}O_9N_8$ | vLIILIIL | this study |
| **32** | 943.66 | $C_{50}H_{86}O_9N_8$ | vLIILfIL | this study |
| **33** | 957.67 | $C_{51}H_{88}O_9N_8$ | lLfILIIL | this study |
| **34** | 957.67 | $C_{51}H_{88}O_9N_8$ | lLIfLIIL | this study |
| **35** | 459,30 | $C_{25}H_{38}N_4O_4$ | cylco(vLfV) | this study |

| | | | | |
|---|---|---|---|---|
| **36** | 425,31 | $C_{22}H_{40}N_4O_4$ | cyclo(vLIV) | this study |
| **37** | 778,45 | $C_{41}H_{59}N_7O_8$ | vLvvYW | this study |
| **38** | 538.40 | $C_{28}H_{51}N_5O_5$ | cyclo(vLvIL) | this study |
| **39** | 552.41 | $C_{29}H_{53}N_5O_5$ | cyclo(ILvIL) | this study |
| **40** | 425.31 | $C_{22}H_{40}N_4O_4$ | cyclo(ILvV) | this study |
| **41** | 826.45 | $C_{45}H_{59}N_7O_8$ | vLfvYW | this study |
| **42** | 840.47 | $C_{46}H_{61}N_7O_8$ | ILfvYW | this study |
| **43** | 792.47 | $C_{42}H_{61}N_7O_8$ | vLIvYW | this study |
| **44** | 806.48 | $C_{43}H_{63}N_7O_8$ | ILIvYW | this study |
| **45** | 792.47 | $C_{42}H_{61}N_7O_8$ | ILvvYW | this study |
| **46** | 476.30 | $C_{25}H_{40}N_4O_5$ | vLfV | this study |
| **47** | 490.32 | $C_{26}H_{42}N_4O_5$ | ILfV | this study |
| **48** | 314.27 | $C_{18}H_{35}NO_3$ | 12:0-L | this study |
| **49** | 328.29 | $C_{19}H_{37}NO_3$ | 13:0-L | this study |
| **50** | 342.20 | $C_{20}H_{39}NO_3$ | 14:0-L | this study |
| **51** | 455.38 | $C_{26}H_{50}N_2O_4$ | 14:0-LI | this study |
| **52** | 510.39 | $C_{28}H_{51}N_3O_5$ | 10:0(β-hydroxy)-LLL | this study |
| **53** | 496.37 | $C_{27}H_{49}N_3O_5$ | 10:0(β-hydroxy)-LLV | this study |
| **54** | 500.20 | $C_{26}H_{49}N_3O_6$ | 10:0(β-hydroxy)-LVV | this study |
| **55** | 556.35 | $C_{27}H_{49}N_5O_5S$ | IC*LIL | [1] |

**Table S2.** Strains used in this work.

| Strain | Genotype/ NRPS | Reference |
|---|---|---|
| *E. coli* DH10B | F_mcrA (*mrr-hsd*RMS-*mcr*BC), 80*lac*ZΔ, M15, Δ*lac*X74 *rec*A1 *end*A1 *ara*D 139Δ(*ara*, *leu*)7697 *gal*U *gal*K λ *rps*L (*Strr*) *nup*G / - | [5] |
| *E. coli* DH10B::*mta*A | DH10B with *mtaA* from pCK_*mtaA*Δ*entD* / - | [6] |

| | | |
|---|---|---|
| *P. luminescens* TTO1 | - / *gxpS* | DSMZ |
| *X. bovienii SS-2004* | - / *garS* | [7] |
| *X. nematophila ATCC 19061* | - / *xtpS* | ATCC |
| *X. szentirmaii DSM16338* | - / *szeS* | DSMZ |
| *X. indica DSM 17382* | - / *xldS* | DSMZ |

**Table S3.** Plasmids used in this work.

| Plasmids | Genotype | Reference |
|---|---|---|
| pCOLA_ara/tacI | ori ColA, $kan^R$, *araC-$P_{BAD}$* and *tac*I | unpublished |
| pCK_0402 | ori p15A, $cm^R$, *araC-$P_{BAD}$* and *tac*I-*araE* | [1] |
| pCOLA_ara_xtpS_tacI_JW | ori ColA, $kan^R$, *araC-$P_{BAD}$ xtpS* and *tac*I | [1] |
| pCOLA_ara_gxpS_tacI_JW | ori ColA, $kan^R$, *araC-$P_{BAD}$ gxpS* and *tac*I | [1] |
| pNA2 | ori p15A, $cm^R$, *araC-$P_{BAD}$ xtpS_* $A_1T_1C/E_2A_2T_2$-SYNZIP17 und *tac*I-*araE* | this study |
| pNA3 | ori ColA, $kan^R$, *araC-$P_{BAD}$* SYNZIP18-*xtpS_* $C_3A_3T_3C/E_4A_4T_4TE$ und *tac*I | this study |
| pNA4 | ori p15A, $cm^R$, *araC-$P_{BAD}$ xtpS_* $A_1T_1C/E_2A_2$-SYNZIP17 und *tac*I-*araE* | this study |
| pNA5 | ori ColA, $kan^R$, *araC-$P_{BAD}$* SYNZIP18-*xtpS_* $T_2C_3A_3T_3C/E_4A_4T_4TE$ und *tac*I | this study |
| pNA15 | ori ColA, $kan^R$, *araC-$P_{BAD}$* SYNZIP18-*xtpS*_$T_2C_3A_3$-SYNZIP1 and *tac*I | this study |
| pNA16 | ori CloDF13, $spec^R$, *araC-$P_{BAD}$* SYNZIP2-*xtpS*_$T_3C/E_4A_4T_4TE$ and *tac*I | this study |
| pNA17 | ori ColA, $kan^R$, *araC-$P_{BAD}$* SYNZIP18-*xtpS*_$C_3A_3T_4$-SYNZIP1 and *tac*I | this study |
| pNA18 | ori CloDF13, $spec^R$, *araC-$P_{BAD}$* SYNZIP2-*xtpS*_$C/E_4A_4T_4TE$ and *tac*I | this study |
| pNA24 | ori ColA, $kan^R$, *araC-$P_{BAD}$* SYNZIP18-*xldS*_$T_2C_3A_3$-SYNZIP1 and *tac*I | this study |
| pNA26 | ori p15A, $cm^R$, *araC-$P_{BAD}$ gxpS*_$A_1T_1C/E_2A_2$-SYNZIP17 and *tac*I-*araE* | this study |
| pNA27 | ori ColA, $kan^R$, *araC-$P_{BAD}$* SYNZIP18-*gxpS*_$T_2C_3A_3$-SYNZIP1 and *tac*I | this study |
| pNA28 | ori CloDF13, $spec^R$, *araC-$P_{BAD}$* SYNZIP2-*gxpS*_$T_3C/E_4A_4T_4$ $C/E_5A_5T_5TE$ and *tac*I | this study |
| pNA29 | ori p15A, $cm^R$, *araC-$P_{BAD}$ sze*S_ $C_1A_1T_1C/E_2A_2$-SYNZIP17 and *tac*I-*araE* | this study |
| pNA30 | ori ColA, $kan^R$, *araC-$P_{BAD}$* SYNZIP18-*szeS*_$T_2C_3A_3$-SYNZIP1 and *tac*I | this study |

| | | |
|---|---|---|
| pNA31 | ori CloDF13, $\text{spec}^R$, *araC-$P_{BAD}$* SYNZIP2-*szeS*_$T_3C/E_4A_4T_4C/E_5A_5T_5C_6A_6T_6$TE and *tac*I | this study |
| pNA35 | ori ColA, $\text{kan}^R$, *araC-$P_{BAD}$* SYNZIP18-*garS*_$T_2C_3A_3$-SYNZIP1 and *tac*I | this study |
| pNA104 | ori p15A, $\text{cm}^R$, *araC-$P_{BAD}$ gxpS*_ $A_1T_1C/E_2$-SYNZIP17 und *tac*I-*araE* | this study |
| pNA105 | ori ColA, $\text{kan}^R$, *araC-$P_{BAD}$* SYNZIP18-*gxpS*_ $A_2T_2C_3A_3T_3C/E_4A_4T_4C/E_5A_5T_5$TE und *tac*I | this study |
| pNA106 | ori p15A, $\text{cm}^R$, *araC-$P_{BAD}$ gxpS*_ $A_1T_1C/E_2A_2T_2C_3A_3T_3C/E_4$-SYNZIP17 und *tac*I-*araE* | this study |
| pNA107 | ori ColA, $\text{kan}^R$, *araC-$P_{BAD}$* SYNZIP18-*gxpS*_ $A_4T_4C/E_5A_5T_5$TE und *tac*I | this study |
| pNA108 | ori p15A, $\text{cm}^R$, *araC-$P_{BAD}$ gxpS*_ $A_1T_1C/E_2A_2T_2C_3A_3T_3C/E_4A_4T_4C/E_5$-SYNZIP17 und *tac*I-*araE* | this study |
| pNA109 | ori ColA, $\text{kan}^R$, *araC-$P_{BAD}$* SYNZIP18-*gxpS*_$A_5T_5$TE und *tac*I | this study |
| pNA59 | ori p15A, $\text{cm}^R$, *araC-$P_{BAD}$ xfpS*_$C_1A_1$-SYNZIP17 and *tac*I-*araE* | this study |
| pNA60 | ori ColA, $\text{kan}^R$, *araC-$P_{BAD}$* SYNZIP18-*xfpS*_$T_1E_1C_2A_2T_2C_3A_3T_3$TE and *tac*I | this study |
| pJW61 | ori p15A, $\text{cm}^R$, *araC-$P_{BAD}$ xtpS*_$A_1T_1C/E_2A_2T_2C_3$-SYNZIP17 and *tac*I-*araE* | [1] |
| pJW62 | ori ColA, $\text{kan}^R$, *araC-$P_{BAD}$* SYNZIP18-*xtpS*_$A_3T_3C/E_4A_4T_4$TE and *tac*I | [1] |
| pJW118 | ori p15A, $\text{cm}^R$, *araC-$P_{BAD}$ bacA*_A1T1CyA2T2C3A3-SYNZIP17 and *tac*I-*araE* | this study |
| pJW120 | ori ColA, $\text{kan}^R$, *araC-$P_{BAD}$* SYNZIP18-*bacA*_T3$C_{Dsub}$4-*sfrA-BC*_$C_{Asub}$6A6T6E6C7A7T7TE and *tac*I | this study |
| pJW122 | ori p15A, $\text{cm}^R$, *araC-$P_{BAD}$ bacA*_A1T1CyA2T2C3A3T3-SYNZIP17 and *tac*I-*araE* | this study |
| pJW124 | ori ColA, $\text{kan}^R$, *araC-$P_{BAD}$* SYNZIP18-*bacA*_$C_{Dsub}$4-*sfrA-BC*_$C_{Asub}$6A6T6E6C7A7T7TE and *tac*I | this study |
| pJW126 | ori p15A, $\text{cm}^R$, *araC-$P_{BAD}$ bacA*_A1T1CyA2T2C3A3T3 $C_{Dsub}$4-SYNZIP17 and *tac*I-*araE* | this study |
| pJW128 | ori ColA, $\text{kan}^R$, *araC-$P_{BAD}$* SYNZIP18-*sfrA-BC*_$C_{Asub}$6A6T6E6C7A7T7TE and *tac*I | this study |
| pJW141 | ori p15A, $\text{cm}^R$, *araC-$P_{BAD}$ xldS*_$C_1$-SYNZIP17 and *tac*I-*araE* | this study |

**Table S4.** Oligonucleotides used in this work.

| Plasmids | Oligo-nucleotides | Sequence (5' →3'; overlapping ends) | Template |
|---|---|---|---|
| pNA2 | KB-pACYC-II-FW | AACGAGAAGGAGGAATTAAAATCG | pJW61 |
| | KB-pACYC-II-RV | CATGGAATTCCTCCTGTTAGC | pJW61 |
| | na03_FW | TGGGCTAACAGGAGGAATTCCATGAAAGATAGCATGGCTAAAAAGGG | *X. nematophila* ATCC 19061 |
| | na05_RV | CGATTTTAATTCCTCCTTCTCGTTAACACGATCACGGGATATTG | *X. nematophila* ATCC 19061 |
| pNA3 | KB-pCOLA-II-FW | TGACAATTAATCATCGGCTCG | pJW62 |
| | KB-pCOLA-II-RV | TGAGATAGCTGCAGTCAGCTCG | pJW62 |
| | na06_FW | AACGAGCTGACTGCAGCTATCTCATTGCCTTTATCGTTTGGTCAAC | *X. nematophila* ATCC 19061 |
| | na07_RV | CGAGCCGATGATTAATTGTCACAGCGCCTCCACTTCG | *X. nematophila* ATCC 19061 |
| pNA4 | KB-pACYC-II-FW | AACGAGAAGGAGGAATTAAAATCG | pJW61 |
| | KB-pACYC-II-RV | CATGGAATTCCTCCTGTTAGC | pJW61 |
| | na03_FW | TGGGCTAACAGGAGGAATTCCATGAAAGATAGCATGGCTAAAAAGGG | *X. nematophila* ATCC 19061 |
| | na13_RV | CGATTTTAATTCCTCCTTCTCGTTATAAATCTGGCGGGCGAA | *X. nematophila* ATCC 19061 |
| pNA5 | KB-pCOLA-II-FW | TGACAATTAATCATCGGCTCG | pJW62 |
| | KB-pCOLA-II-RV | TGAGATAGCTGCAGTCAGCTCG | pJW62 |
| | na14_FW | AACGAGCTGACTGCAGCTATCTCAGTTGCGCCACAAGGAGAA | *X. nematophila* ATCC 19061 |
| | na07_RV | CGAGCCGATGATTAATTGTCACAGCGCCTCCACTTCG | *X. nematophila* ATCC 19061 |
| pNA15 | na28_FW | CACAAAAAAGACCTGATCGCGTACCTGGAGAAAGAAATCGCGA ATCTGCGTAAGAAAATCGAAGAATGACAATTAATCATCGGCTCG | pJW62 |
| | na29_FW | AACCTGGTTGCGCAGCTCGAAAACGAAGTTGCGTCTCTGGAAAATGAGA ACGAAACCCTGAAGAAAAAGAACCTGCACAAAAAAGACCTGATCGCGTAC | pCOLA_ara/tacI_SZ18_halfSZ1 |
| | na30_RV | TGAGATAGCTGCAGTCAGCTCG | pJW62/ pCOLA_ara/tacI_SZ18_halfSZ1 |
| | na14_FW | GGCTAACAGGAGGAATTCCATGGTTGCGCCACAAGGAGAA | *X. nematophila* ATCC 19061 |
| | na31_RV | AGCTGCGCAACCAGGTTATAGACCTGCCGGGCAAAC | *X. nematophila* ATCC 19061 |
| pNA16 | na32_RV | GTTCTGTTCATCACGTTCCAGCTGCAGGTTGTCTTTTTTCAGACGTGCGA TTTTCTTACGCAGATACGCGTTACGCGCCATGGAATTCCTCCTGTTAGCC | pCDF_ara/tacI |
| | na33_RV | CTGTTCGTGAGACGCAACTTCGTTTTCGAGACGCGCGATTTCGTCAC GCAGGTTCGCGATGATTTTTTCCAGGTTCTGTTCATCACGTTCCAGC | pCDF_ara/tacI_halbSZ2 |
| | na34_FW | TGACAATTAATCATCGGCTCG | pCDF_ara/tacI/ pCDF_ara/tacI_halbSZ2 |
| | na35_FW | AAACGAAGTTGCGTCTCACGAACAGGCGGCTCCGCAGGG | *X. nematophila* ATCC 19061 |
| | na7_RV | CGAGCCGATGATTAATTGTCACAGCGCCTCCACTTCG | *X. nematophila* ATCC 19061 |
| | na28_FW | CACAAAAAAGACCTGATCGCGTACCTGGAGAAAGAAATCGCGA ATCTGCGTAAGAAAATCGAAGAATGACAATTAATCATCGGCTCG | pJW62 |
| | na29_FW | AACCTGGTTGCGCAGCTCGAAAACGAAGTTGCGTCTCTGGAAAATGAGA ACGAAACCCTGAAGAAAAAGAACCTGCACAAAAAAGACCTGATCGCGTAC | pCOLA_ara/tacI_SZ18_halfSZ1 |
| | na30_RV | TGAGATAGCTGCAGTCAGCTCG | pJW62/ pCOLA_ara/tacI_SZ18_halfSZ1 |

| | | | |
|---|---|---|---|
| pNA17 | na36_FW | AACGAGCTGACTGCAGCTATCTCATTGCCTTTATCGTTTGGTCAACAG | *X. nematophila* ATCC 19061 |
| | na37_RV | CGTTTTCGAGCTGCGCAACCAGGTTCATGGCTGGCGTTAGTACCG | *X. nematophila* ATCC 19061 |
| pNA18 | na32_RV | GTTCTGTTCATCACGTTCCAGCTGCAGGTTGTCTTTTTTCAGACGTGCGA TTTTCTTACGCAGATACGCGTTACGCGCCATGGAATTCCTCCTGTTAGCC | pCDF_ara/tacI |
| | na33_RV | CTGTTCGTGAGACGCAACTTCGTTTTCGAGACGCGCGATTTCGTCAC GCAGGTTCGCGATGATTTTTTCCAGGTTCTGTTCATCACGTTCCAGC | pCDF_ara/tacI_halbSZ2 |
| | na34_FW | TGACAATTAATCATCGGCTCG | pCDF_ara/tacI/ pCDF_ara/tacI_halbSZ2 |
| | na38_FW | AAACGAAGTTGCGTCTCACGAACAGTTGCCGCTGATTGATCTCAC | *X. nematophila* ATCC 19061 |
| | na7_RV | CGAGCCGATGATTAATTGTCACAGCGCCTCCACTTCG | *X. nematophila* ATCC 19061 |
| pNA24 | na28_FW | CACAAAAAAGACCTGATCGCGTACCTGGAGAAAGAAATCGCGA ATCTGCGTAAGAAAATCGAAGAATGACAATTAATCATCGGCTCG | pJW62 |
| | na29_FW | AACCTGGTTGCGCAGCTCGAAAACGAAGTTGCGTCTCTGGAAAATGAGA ACGAAACCCTGAAGAAAAAGAACCTGCACAAAAAAGACCTGATCGCGTAC | pCOLA_ara/tacI_SZ18_ halfSZ1 |
| | na30_RV | TGAGATAGCTGCAGTCAGCTCG | pJW62/ pCOLA_ara/tacI_SZ18_ halfSZ1 |
| | na47_FW | AACGAGCTGACTGCAGCTATCTCAGAAGCGCCCATTGGCAAA | *X. indica* DSM 17382 |
| | na48_RV | CGTTTTCGAGCTGCGCAACCAGGTTATAGCCACGTGTAACAACCGCTG | *X. indica* DSM 17382 |
| pNA26 | KB-pACYC-II-FW | AACGAGAAGGAGGAATTAAAATCG | pJW61 |
| | KB-pACYC-II-RV | CATGGAATTCCTCCTGTTAGC | pJW61 |
| | na51 | GCTAACAGGAGGAATTCCATGAAAGATAGCATGGCTAAAAAGGAAAT | *P. luminescens* TT01 |
| | na52 | CGATTTTAATTCCTCCTTCTCGTTATAAATTTGGCGAGCAAAAGC | *P. luminescens* TT01 |
| pNA27 | na28_FW | CACAAAAAAGACCTGATCGCGTACCTGGAGAAAGAAATCGCGA ATCTGCGTAAGAAAATCGAAGAATGACAATTAATCATCGGCTCG | pJW62 |
| | na29_FW | AACCTGGTTGCGCAGCTCGAAAACGAAGTTGCGTCTCTGGAAAATGAGA ACGAAACCCTGAAGAAAAAGAACCTGCACAAAAAAGACCTGATCGCGTAC | pCOLA_ara/tacI_SZ18_ halfSZ1 |
| | na30_RV | TGAGATAGCTGCAGTCAGCTCG | pJW62/ pCOLA_ara/tacI_SZ18_ halfSZ1 |
| | na53 | CGAGCTGACTGCAGCTATCTCAGTCGCGCCACAGGGAG | *P. luminescens* TT01 |
| | na534 | CGTTTTCGAGCTGCGCAACCAGGTTGTAAGCTTGGCGAGCAAAGG | *P. luminescens* TT01 |
| pNA28 | na32_RV | GTTCTGTTCATCACGTTCCAGCTGCAGGTTGTCTTTTTTCAGACGTGCGA TTTTCTTACGCAGATACGCGTTACGCGCCATGGAATTCCTCCTGTTAGCC | pCDF_ara/tacI |
| | na33_RV | CTGTTCGTGAGACGCAACTTCGTTTTCGAGACGCGCGATTTCGTCAC GCAGGTTCGCGATGATTTTTTCCAGGTTCTGTTCATCACGTTCCAGC | pCDF_ara/tacI_halbSZ2 |
| | na34_FW | TGACAATTAATCATCGGCTCG | pCDF_ara/tacI/ pCDF_ara/tacI_halbSZ2 |
| | na55 | GAAGTTGCGTCTCACGAACAGCAAGCGCCACAAGGGGA | *P. luminescens* TT01 |
| | na56 | CGAGCCGATGATTAATTGTCACAGCGCCTCCGCTTCAC | *P. luminescens* TT01 |
| pNA29 | KB-pACYC-II-FW | AACGAGAAGGAGGAATTAAAATCG | pJW61 |
| | KB-pACYC-II-RV | CATGGAATTCCTCCTGTTAGC | pJW61 |
| | na57 | GCTAACAGGAGGAATTCCATGAAAGGTAGTATTGCTAAAAAGGGAGATG | *X. szentirmaii* DSM16338 |
| | na58 | CGATTTTAATTCCTCCTTCTCGTTATAATGCTGACGGGCAAATG | *X. szentirmaii* DSM16338 |
| pNA30 | na28_FW | CACAAAAAAGACCTGATCGCGTACCTGGAGAAAGAAATCGCGA ATCTGCGTAAGAAAATCGAAGAATGACAATTAATCATCGGCTCG | pJW62 |
| | na29_FW | AACCTGGTTGCGCAGCTCGAAAACGAAGTTGCGTCTCTGGAAAATGAGA ACGAAACCCTGAAGAAAAAGAACCTGCACAAAAAAGACCTGATCGCGTAC | pCOLA_ara/tacI_SZ18_ halfSZ1 |
| | na30_RV | TGAGATAGCTGCAGTCAGCTCG | pJW62/ pCOLA_ara/tacI_SZ18_ halfSZ1 |
| | na59 | CGAGCTGACTGCAGCTATCTCAGAATCCCCACAAGGGGAGA | *X. szentirmaii* DSM16338 |
| | na60 | TCGAGCTGCGCAACCAGGTTATAATGCTGACGGGCAAACG | *X. szentirmaii* DSM16338 |
| | na32_RV | GTTCTGTTCATCACGTTCCAGCTGCAGGTTGTCTTTTTTCAGACGTGCGA TTTTCTTACGCAGATACGCGTTACGCGCCATGGAATTCCTCCTGTTAGCC | pCDF_ara/tacI |

| | | | |
|---|---|---|---|
| pNA31 | na33_RV | CTGTTCGTGAGACGCAACTTCGTTTTCGAGACGCGCGATTTCGTCAC<br>GCAGGTTCGCGATGATTTTTTCCAGGTTCTGTTCATCACGTTCCAGC | pCDF_ara/tacI_halbSZ2 |
| | na34_FW | TGACAATTAATCATCGGCTCG | pCDF_ara/tacI/<br>pCDF_ara/tacI_halbSZ2 |
| | na61 | AAAACGAAGTTGCGTCTCACGAACAGGAGTTGCCACAAGGTGAAA | *X. szentirmaii*<br>DSM16338 |
| | na62 | CGAGCCGATGATTAATTGTCAAATATTATTTACTATATTGTATTCCTCTGTACCA | *X. szentirmaii*<br>DSM16338 |
| pNA35 | na28_FW | CACAAAAAAGACCTGATCGCGTACCTGGAGAAAGAAATCGCGA<br>ATCTGCGTAAGAAAATCGAAGAATGACAATTAATCATCGGCTCG | pJW62 |
| | na29_FW | AACCTGGTTGCGCAGCTCGAAAACGAAGTTGCGTCTCTGGAAAATGAGA<br>ACGAAACCCTGAAGAAAAAGAACCTGCACAAAAAAGACCTGATCGCGTAC | pCOLA_ara/tacI_SZ18_<br>halfSZ1 |
| | na30_RV | TGAGATAGCTGCAGTCAGCTCG | pJW62/<br>pCOLA_ara/tacI_SZ18_<br>halfSZ1 |
| | na68 | TAACGAGCTGACTGCAGCTATCTCAGAAACTCCGGTCGGTAAAGTAG | *X. bovienii* SS-2004 |
| | na69 | TTCGTTTTCGAGCTGCGCAACCAGGTTATAGCCGCGCACCACTAC | *X. bovienii* SS-2004 |
| pNA59 | KB-pACYC-FW | GAACAGTTAAAACAGAAGCGTGAACAATTAAAGCAAAAGATCGCCAATC<br>TGCGTAAGGAGATCGAAGCCTACAAGTGACAATTAATCATCGGCTCG | pCK_0402 |
| | KB-pACYC-RV | TTCACGCTTCTGTTTTAACTGTTCGATGCGATTACGCAATTCAGCCTTT<br>TTCGATTTTAATTCCTCCTTCTCGTTCATGGAATTCCTCCTGTTAGC | pCK_0402 |
| | na125_FW | GCTAACAGGAGGAATTCCATGGATAACATTCTGGCCTCG | *X. bovienii* SS2004 |
| | na126_RV | CAGATTTTAAACAGAGCCGCTATGTTTATAACGAGAAGGAGGAATTAAAATCG | *X. bovienii* SS2004 |
| pNA60 | KB-pCOLA-FW | CATTGACAAAGAGCTGCGTGCCAACGAAAACGAACTTCGCGCCCTTGA<br>TAACGAGCTGACTGCAGCTATCTCATGACAATTAATCATCGGCTCG | pCOLA_ara/tacI |
| | KB-pCOLA-RV | TTGGCACGCAGCTCTTTGTCAATGGCATTTAACTCGCGGTCCAAGGCT<br>TTCAGTTCACGCTCTTCAGCATAGAACATGGAATTCCTCCTGTTAGC | pCOLA_ara/tacI |
| | na127_FW | AACGAGCTGACTGCAGCTATCTCACGTGCCGCAGAAACGG | *X. bovienii* SS2004 |
| | na128_RV | ATACGAGCCGATGATTAATTGTCAATCCACCAGCTCCAACAC | *X. bovienii* SS2004 |
| pNA104 | KB-pACYC-II-FW | AACGAGAAGGAGGAATTAAAATCG | pJW61 |
| | KB-pACYC-II-RV | CATGGAATTCCTCCTGTTAGC | pJW61 |
| | na194_FW | GGCTAACAGGAGGAATTCCATGAAAGATAGCATGGCTAAAAAGG | *P. luminescens* TT01 |
| | na208_RV | CGATTTTAATTCCTCCTTCTCGTTCCACGTTTCCAGCAACAAC | *P. luminescens* TT01 |
| pNA105 | KB-pCOLA-II-FW | TGACAATTAATCATCGGCTCG | pJW62 |
| | KB-pCOLA-II-RV | TGAGATAGCTGCAGTCAGCTCG | pJW62 |
| | na209_FW | CGAGCTGACTGCAGCTATCTCAAACGCGACAGAAACCCC | *P. luminescens* TT01 |
| | na56_RV | CGAGCCGATGATTAATTGTCACAGCGCCTCCGCTTCAC | *P. luminescens* TT01 |
| pNA106 | KB-pACYC-II-FW | AACGAGAAGGAGGAATTAAAATCG | pJW61 |
| | KB-pACYC-II-RV | CATGGAATTCCTCCTGTTAGC | pJW61 |
| | na194_FW | GGCTAACAGGAGGAATTCCATGAAAGATAGCATGGCTAAAAAGG | *P. luminescens* TT01 |
| | na210_RV | CGATTTTAATTCCTCCTTCTCGTTCCAAGTTTTTAACAACAATGTGCG | *P. luminescens* TT01 |
| pNA107 | KB-pCOLA-II-FW | TGACAATTAATCATCGGCTCG | pJW62 |
| | KB-pCOLA-II-RV | TGAGATAGCTGCAGTCAGCTCG | pJW62 |
| | na211_FW | CGAGCTGACTGCAGCTATCTCAAACGCCACTGAAACAGCCTATC | *P. luminescens* TT01 |
| | na56_RV | CGAGCCGATGATTAATTGTCACAGCGCCTCCGCTTCAC | *P. luminescens* TT01 |
| | KB-pACYC-II-FW | AACGAGAAGGAGGAATTAAAATCG | pJW61 |
| | KB-pACYC-II-RV | CATGGAATTCCTCCTGTTAGC | pJW61 |

| | | | |
|---|---|---|---|
| pNA108 | na194_FW | GGCTAACAGGAGGAATTCCATGAAAGATAGCATGGCTAAAAAGG | *P. luminescens* TT01 |
| | na212_RV | CGATTTTAATTCCTCCTTCTCGTTGCCGACTTTCAGTAACAGTTTCC | *P. luminescens* TT01 |
| pNA109 | KB-pCOLA-II-FW | TGACAATTAATCATCGGCTCG | pJW62 |
| | KB-pCOLA-II-RV | TGAGATAGCTGCAGTCAGCTCG | pJW62 |
| | na213_FW | CGAGCTGACTGCAGCTATCTCAAATGGCCCGCAAACGG | *P. luminescens* TT01 |
| | na56_RV | CGAGCCGATGATTAATTGTCACAGCGCCTCCGCTTCAC | *P. luminescens* TT01 |
| pJW118 | KB-pACYC-II-FW | AACGAGAAGGAGGAATTAAAATCG | pJW61 |
| | KB-pACYC-II-RV | CATGGAATTCCTCCTGTTAGC | pJW61 |
| | jw208_FW | GCTAACAGGAGGAATTCCATGGTTGCTAAACATTCATTAGAAAATGGG | pFF1_NRPS_6 (*3*) |
| | jw0214_RV | CGATTTTAATTCCTCCTTCTCGTTGTAGCGGCGATCCATTGT | pFF1_NRPS_6 (*3*) |
| pJW120 | KB-pCOLA-II-FW | TGACAATTAATCATCGGCTCG | pJW62 |
| | KB-pCOLA-II-RV | TGAGATAGCTGCAGTCAGCTCG | pJW62 |
| | jw0216_FW | CGAGCTGACTGCAGCTATCTCAGAAGCGCCGCGGG | pFF1_NRPS_6 (*3*) |
| | jw0212_RV | CGAGCCGATGATTAATTGTCATGAAACCGTTACGGTTTGTGTATTA | pFF1_NRPS_6 (*3*) |
| pJW122 | KB-pACYC-II-FW | AACGAGAAGGAGGAATTAAAATCG | pJW61 |
| | KB-pACYC-II-RV | CATGGAATTCCTCCTGTTAGC | pJW61 |
| | jw208_FW | GCTAACAGGAGGAATTCCATGGTTGCTAAACATTCATTAGAAAATGGG | pFF1_NRPS_6 (*3*) |
| | jw0218_RV | CGATTTTAATTCCTCCTTCTCGTTTCGGTAATATGGTTTTTCTTCGG | pFF1_NRPS_6 (*3*) |
| pJW124 | KB-pCOLA-II-FW | TGACAATTAATCATCGGCTCG | pJW62 |
| | KB-pCOLA-II-RV | TGAGATAGCTGCAGTCAGCTCG | pJW62 |
| | jw0220_FW | CGAGCTGACTGCAGCTATCTCA TTGTCTTCAGCGCAAAAAAGG | pFF1_NRPS_6 (*3*) |
| | jw0212_RV | CGAGCCGATGATTAATTGTCA TGAAACCGTTACGGTTTGTGTATTA | pFF1_NRPS_6 (*3*) |
| pJW126 (NRPS-42) | KB-pACYC-II-FW | AACGAGAAGGAGGAATTAAAATCG | pJW61 |
| | KB-pACYC-II-RV | CATGGAATTCCTCCTGTTAGC | pJW61 |
| | jw208_FW | GCTAACAGGAGGAATTCCATGGTTGCTAAACATTCATTAGAAAATGGG | pFF1_NRPS_6 (*3*) |
| | jw0222_RV | CGATTTTAATTCCTCCTTCTCGTTGGCATGGCTATTTTCCCATT | pFF1_NRPS_6 (*3*) |
| pJW128 | KB-pCOLA-II-FW | TGACAATTAATCATCGGCTCG | pJW62 |
| | KB-pCOLA-II-RV | TGAGATAGCTGCAGTCAGCTCG | pJW62 |
| | jw0224_FW | CGAGCTGACTGCAGCTATCTCACAAAAAGAACGGATGAAGGAGC | pFF1_NRPS_6 (*3*) |
| | jw0212_RV | CGAGCCGATGATTAATTGTCATGAAACCGTTACGGTTTGTGTATTA | pFF1_NRPS_6 (*3*) |
| pJW141 | KB-pACYC-II-FW | AACGAGAAGGAGGAATTAAAATCG | pJW61 |
| | KB-pACYC-II-RV | CATGGAATTCCTCCTGTTAGC | pJW61 |
| | jw0166_FW | GCTAACAGGAGGAATTCCATGAAACTTTGGAACTATAAAATGAATATGAC | *X. indica* DSM 17382 |
| | jw0254_RV | CGATTTTAATTCCTCCTTCTCGTTAAAATCTACCAATAGTTTCTGGCGC | *X. indica* DSM 17382 |

**Table S5.** C-A interface identity. WT interface was sequenced aligned with the artificial interface of NRPS-1-16. Identity is depicted in [%].

| NRPS | WT interface | Artificial interface | Identity [%] |
|---|---|---|---|
| NRPS-1 | $C/E_5$-$A_5$ | $C/E_2$-$A_5$ | 89.3 |
| NRPS-2 | $C/E_4$-$A_4$ | $C/E_2$-$A_4$ | 86.1 |
| NRPS-3 | $C/E_5$-$A_5$ | $C/E_3$-$A_5$ | 59.5 |
| NRPS-4 | $C_3$-$A_3$ | $C/E_2$-$A_4$ | 57.9 |
| NRPS-5 | $C/E_4$-$A_4$ | $C_3$-$A_4$ | 59.8 |
| NRPS-6 | $C/E_5$-$A_5$ | $C/E_4$-$A_5$ | 89.3 |
| NRPS-7 | $C/E_2$-$A_2$ | $C_3$-$A_2$ | 58.7 |
| NRPS-8 | $C_3$-$A_3$ | $C/E_4$-$A_3$ | 58.7 |
| NRPS-9 | $C/E_4$-$A_4$ | $C/E_5$-$A_4$ | 89.3 |
| NRPS-10 | $C/E_2$-$A_2$ | $C/E_4$-$A_2$ | 85.2 |
| NRPS-11 | $C_3$-$A_3$ | $C/E_5$-$A_3$ | 59.5 |
| NRPS-12 | $C/E_5$-$A_2$ | $C/E_2$-$A_2$ | 88.5 |

### 3. Supplementary Figures

4.

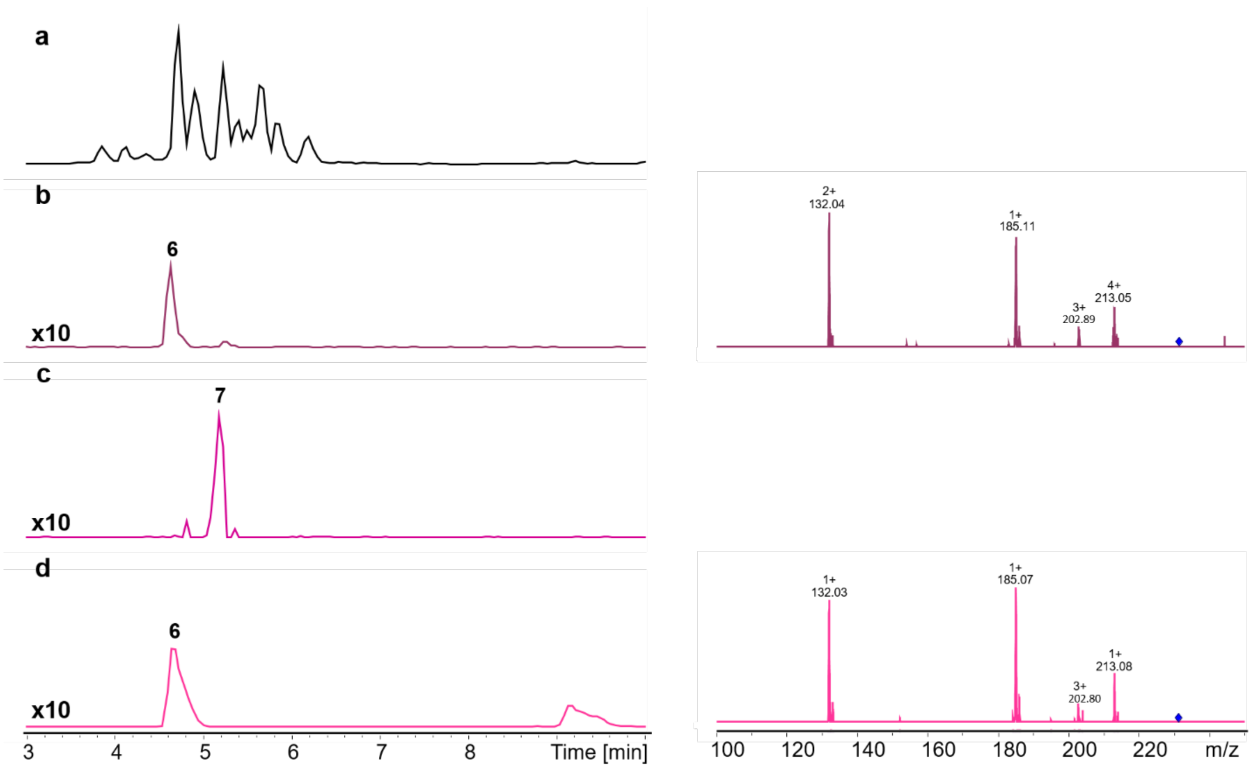

5. **Figure S1. HPLC/MS data (Figure 1b) of compounds 6 and 7 produced in *E. coli* DH10B::*mtaA*.** (a) Base Peak Chromatogram (BPC) of an exemplary culture extract (b) Extracted ion chromatogram (EIC)/MS$^2$ of **6** (*m/z* $[M+H]^+$ = 231.17, **NRPS-1**). (c) EIC of **7** (*m/z* $[M+H]^+$ = 245.19; **NRPS-1**) (d) EIC/MS$^2$ data of synthetic **6** (*m/z* $[M+H]^+$ = 231.17).

6. 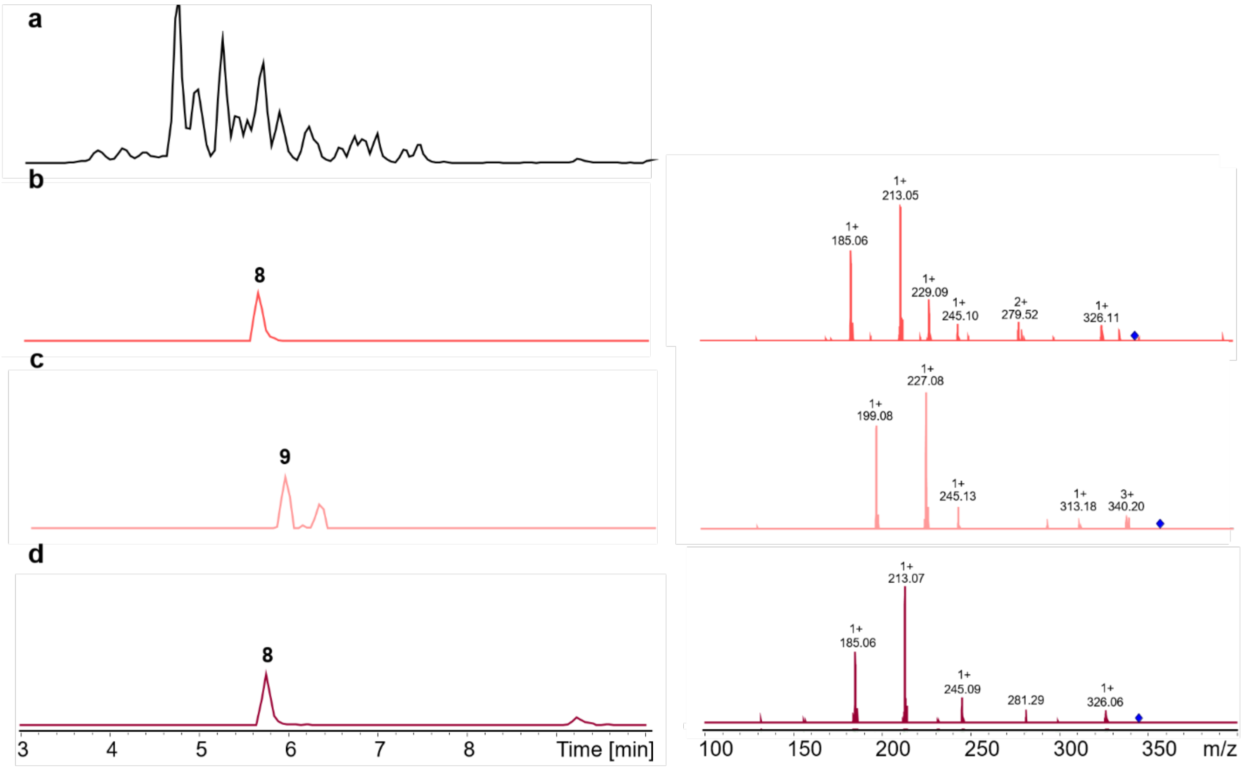

7. **Figure S2. HPLC/MS data (Figure 1b) of compounds 8 and 9 produced in *E. coli* DH10B::*mtaA*.** (a) BPC of an exemplary culture extract. (b) EIC/MS$^2$ of **8** (*m/z* [M+H]$^+$ = 344.25, **NRPS-2**). (c) EIC/MS$^2$ of **9** (*m/z* [M+H]$^+$ = 358.28; **NRPS-2**). (d) EIC/MS$^2$ data of synthetic **8** (*m/z* [M+H]$^+$ = 344.25).

8.

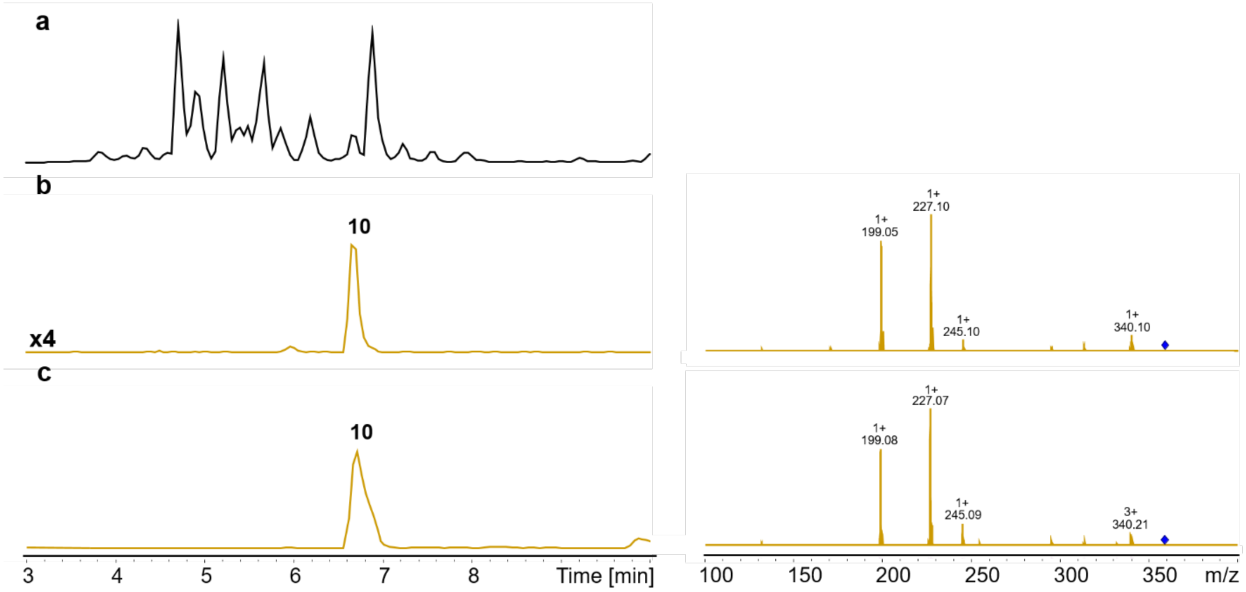

9.

10. **Figure S3. HPLC/MS data (Figure 1b) of compound 10 produced in *E. coli* DH10B::*mtaA*.** (a) BPC of an exemplary culture extract. (b) EIC/MS$^2$ of **10** (*m/z* [M+H]$^+$ = 358.28, **NRPS-3**). (c) EIC/MS$^2$ data of synthetic **10** (*m/z* [M+H]$^+$ = 358.25).

11.

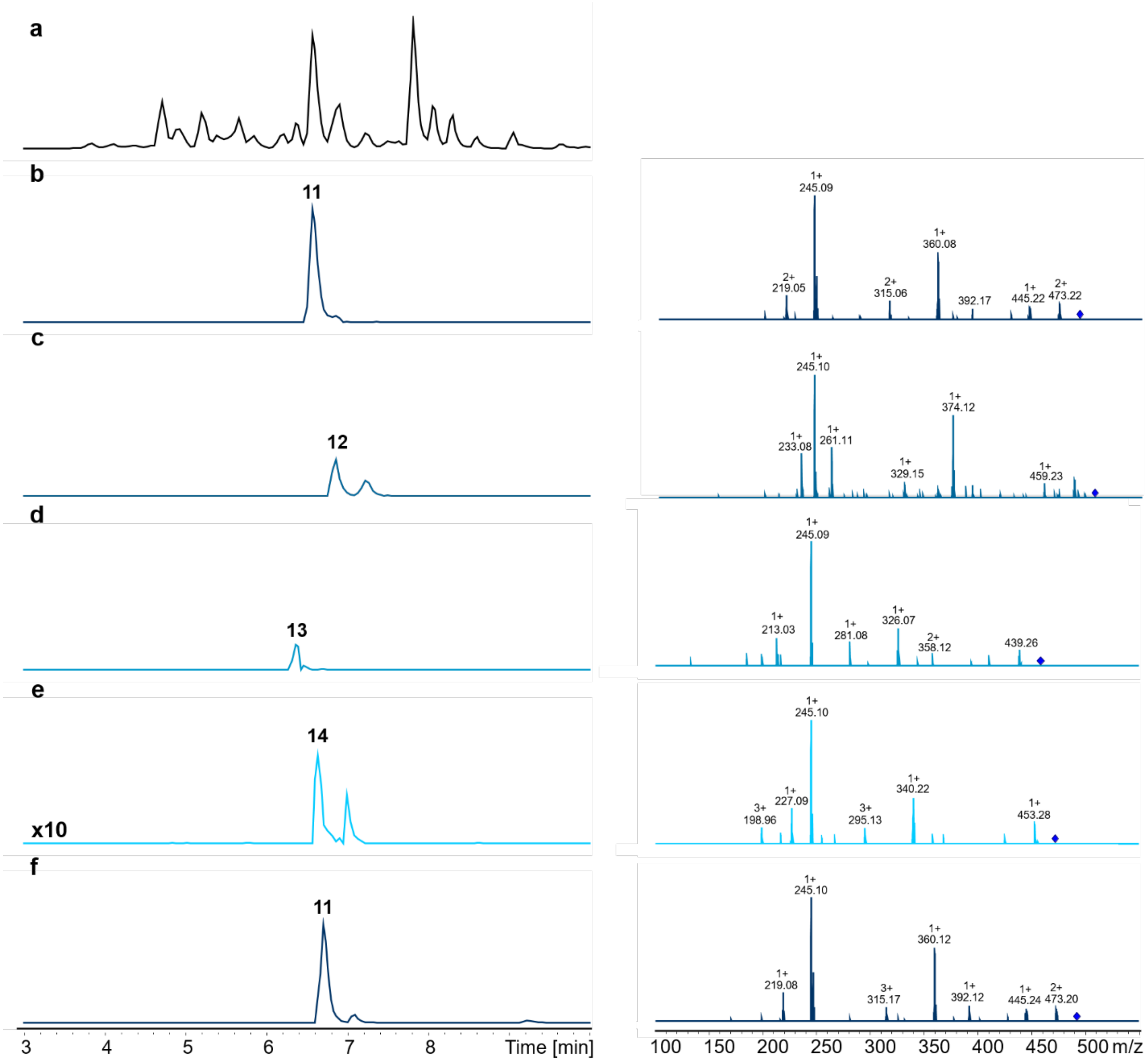

12. **Figure S4. HPLC/MS data (Figure 1b) of compounds 11, 12, 13 and 14 produced in *E. coli* DH10B::*mtaA*.** (a) BPC of an exemplary culture extract (b) EIC/MS$^2$ of **11** (*m/z* [M+H]$^+$ = 491.32, **NRPS-4**). (c) EIC/MS$^2$ data of **12** (*m/z* [M+H]$^+$ = 505.34, **NRPS-4**). (d) EIC/MS$^2$ data of **13** (*m/z* [M+H]$^+$ = 457.34, **NRPS-4**). (d) EIC/MS$^2$ data of **14** (*m/z* [M+H]$^+$ = 471.35, **NRPS-4**). (f) EIC/MS$^2$ data of synthetic **11** (*m/z* [M+H]$^+$ = 491.32).

13.

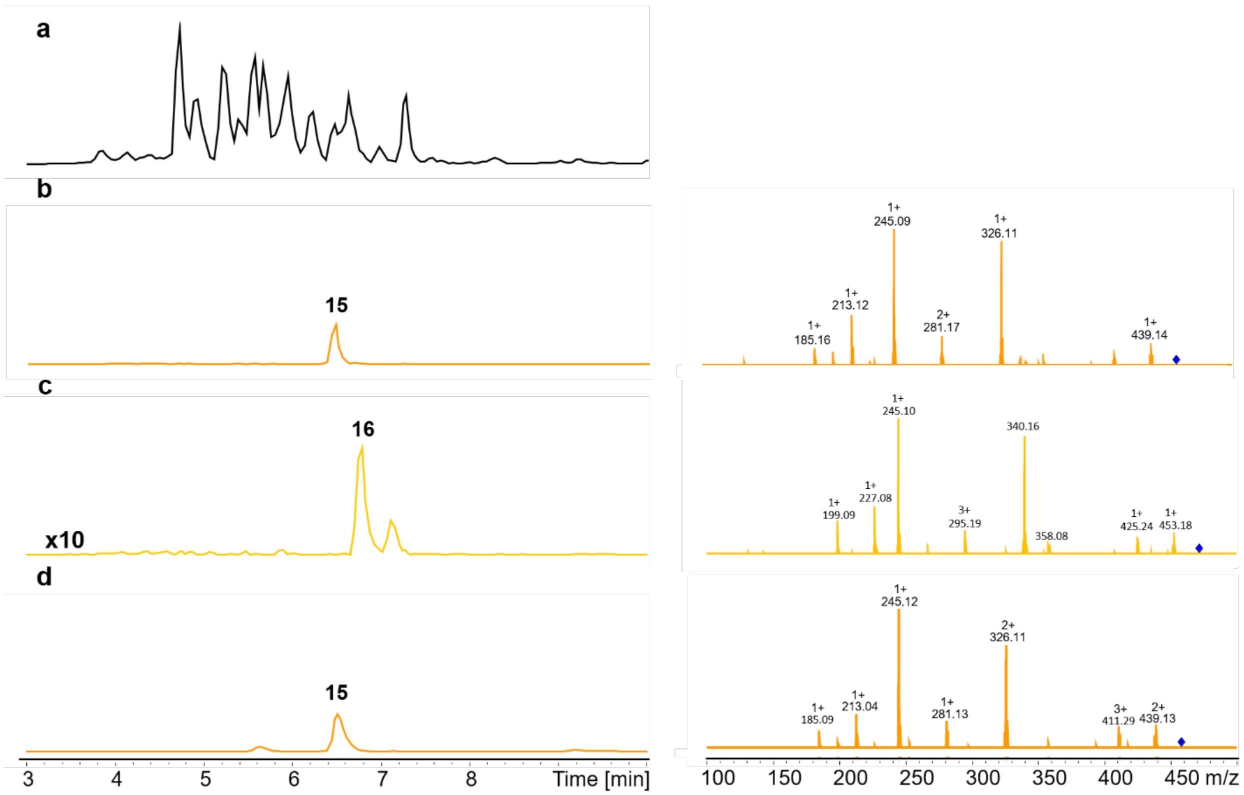

14. **Figure S5. HPLC/MS data (Figure 1b) of compounds 15 and 16 produced in *E. coli* DH10B::*mtaA*.** (a) BPC of an exemplary culture extract (b) EIC/MS$^2$ of **15** (*m/z* $[M+H]^+$ = 457.34, **NRPS-5**). (c) EIC/MS$^2$ data of **16** (*m/z* $[M+H]^+$ = 471.35, **NRPS-5**). (d) EIC/MS$^2$ data of synthetic **15** (*m/z* $[M+H]^+$ = 457.34).

15. 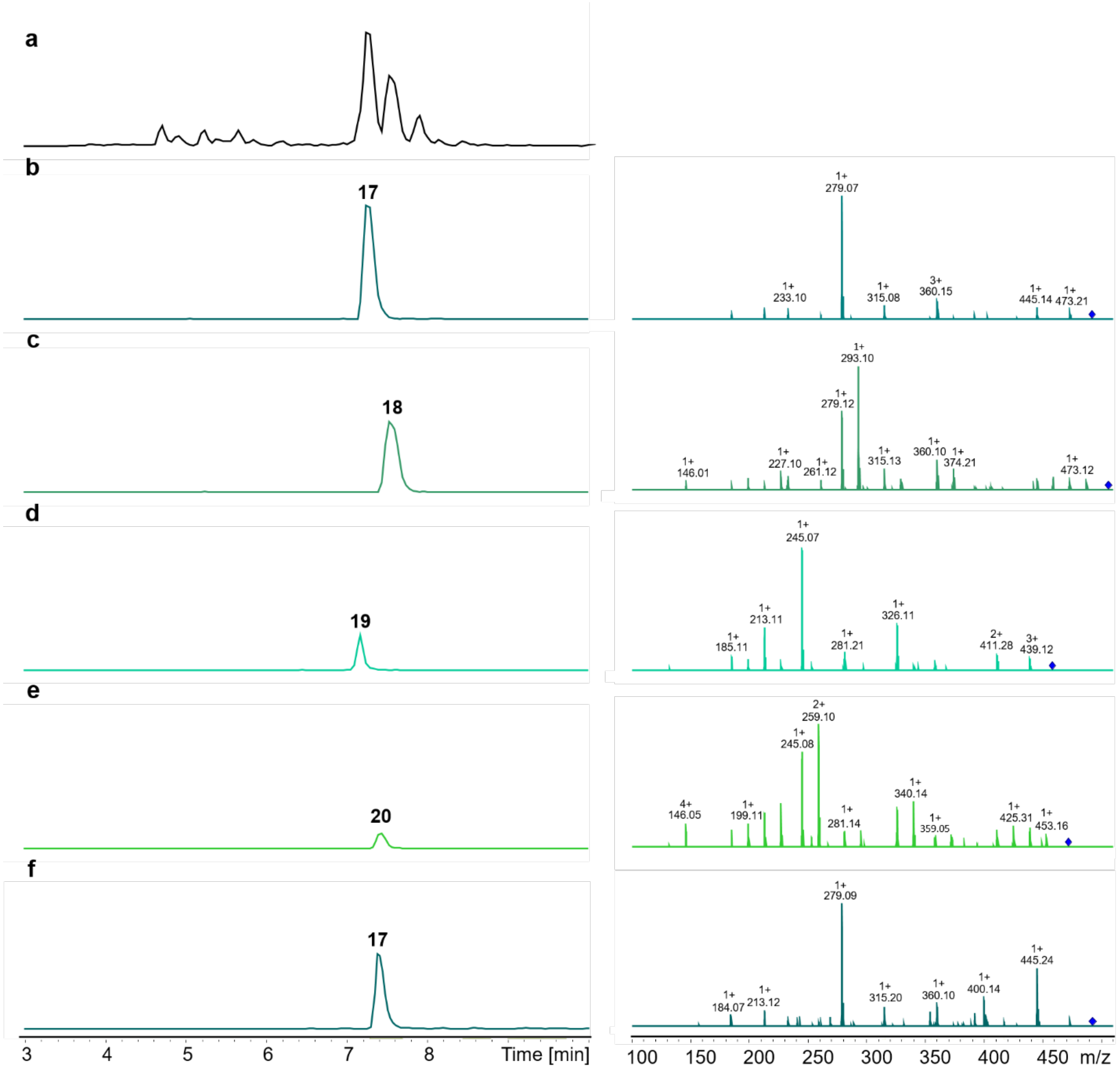

16. **Figure S6. HPLC/MS data (Figure 1b) of compounds 17, 18, 19 and 20 produced in *E. coli* DH10B::*mtaA*.** (a) BPC of an exemplary culture extract (b) EIC/MS$^2$ of **17** (*m/z* [M+H]$^+$ = 491.32, **NRPS-6**). (c) EIC/MS$^2$ data of **18** (*m/z* [M+H]$^+$ = 505.34, **NRPS-6**). (d) EIC/MS$^2$ data of **19** (*m/z* [M+H]$^+$ = 457.34, **NRPS-6**). (e) EIC/MS$^2$ data of **20** (*m/z* [M+H]$^+$ = 471.35). (f) EIC/MS$^2$ of synthetic **17** (*m/z* [M+H]$^+$ = 491.32).

17. 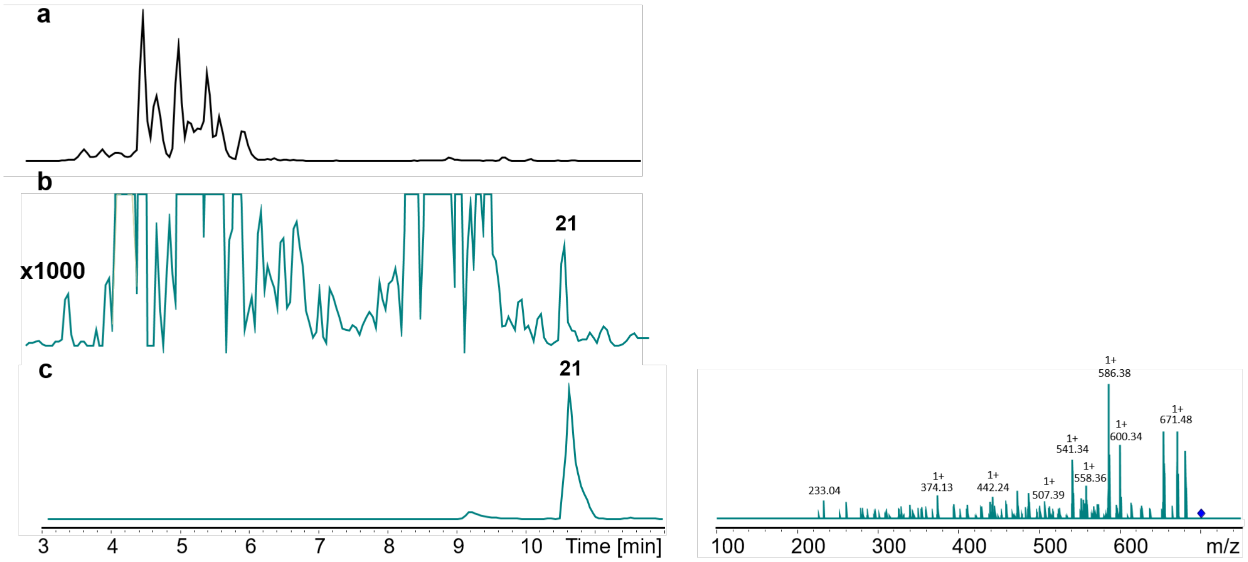

18. **Figure S7. HPLC/MS data (Figure 1c) of compound 21 produced in *E. coli* DH10B::*mtaA*.** (a) BPC of an exemplary culture extract (b) EIC of **21** (*m/z* $[M+H]^+$ = 699.48, **NRPS-7**). (c) EIC/MS$^2$ of synthetic **21** (*m/z* $[M+H]^+$ = 699.48).

19.

20. 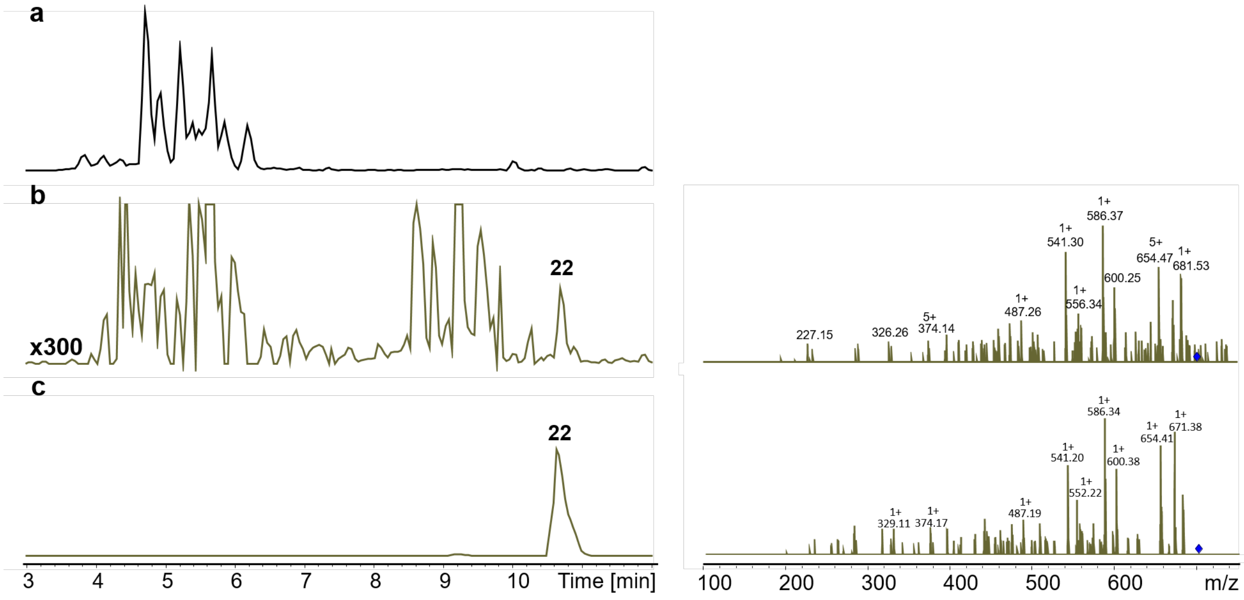

21. **Figure S8. HPLC/MS data (Figure 1c) of compound 22 produced in *E. coli* DH10B::*mtaA*.** (a) BPC of an exemplary culture extract (b) EIC/MS$^2$ of **22** (*m/z* $[M+H]^+$ = 699.48, **NRPS-8**). (c) EIC/MS$^2$ of synthetic **22** (*m/z* $[M+H]^+$ = 699.48).

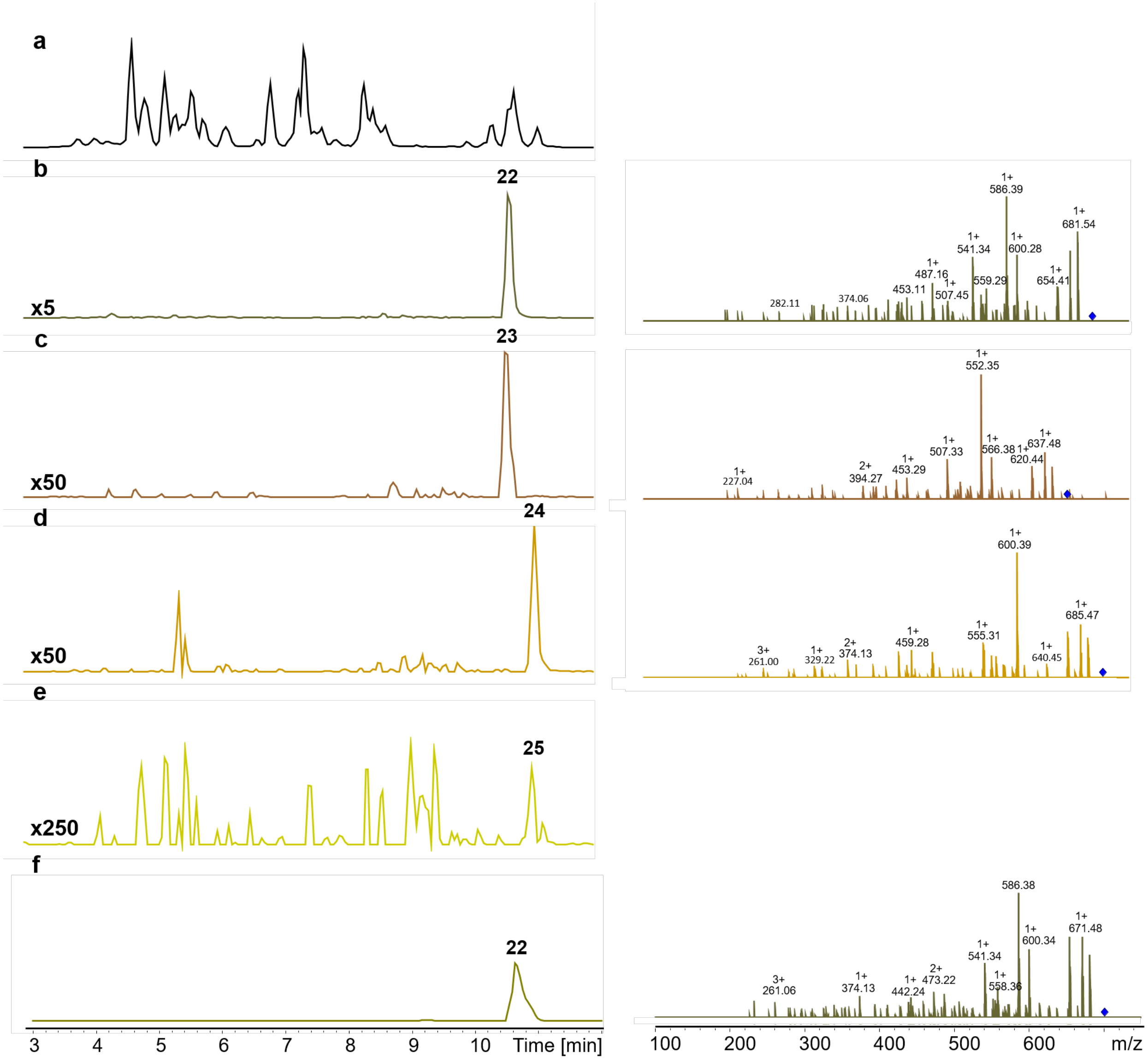

22.

23. **Figure S9. HPLC/MS data (Figure 1b) of compounds 22, 23, 24 and 25 produced in *E. coli* DH10B::*mtaA*.** (a) BPC of an exemplary culture extract (b) EIC/MS$^2$ of **22** (*m/z* [M+H]$^+$ = 699.48, **NRPS-9**). (c) EIC/MS$^2$ data of **23** (*m/z* [M+H]$^+$ = 713.50, **NRPS-9**). (d) EIC/MS$^2$ data of **24** (*m/z* [M+H]$^+$ = 665.50, **NRPS-9**). (e) EIC data of **25** (*m/z* [M+H]$^+$ = 679.51). (f) EIC/MS$^2$ of synthetic **22** (*m/z* [M+H]$^+$ = 491.32).

24.

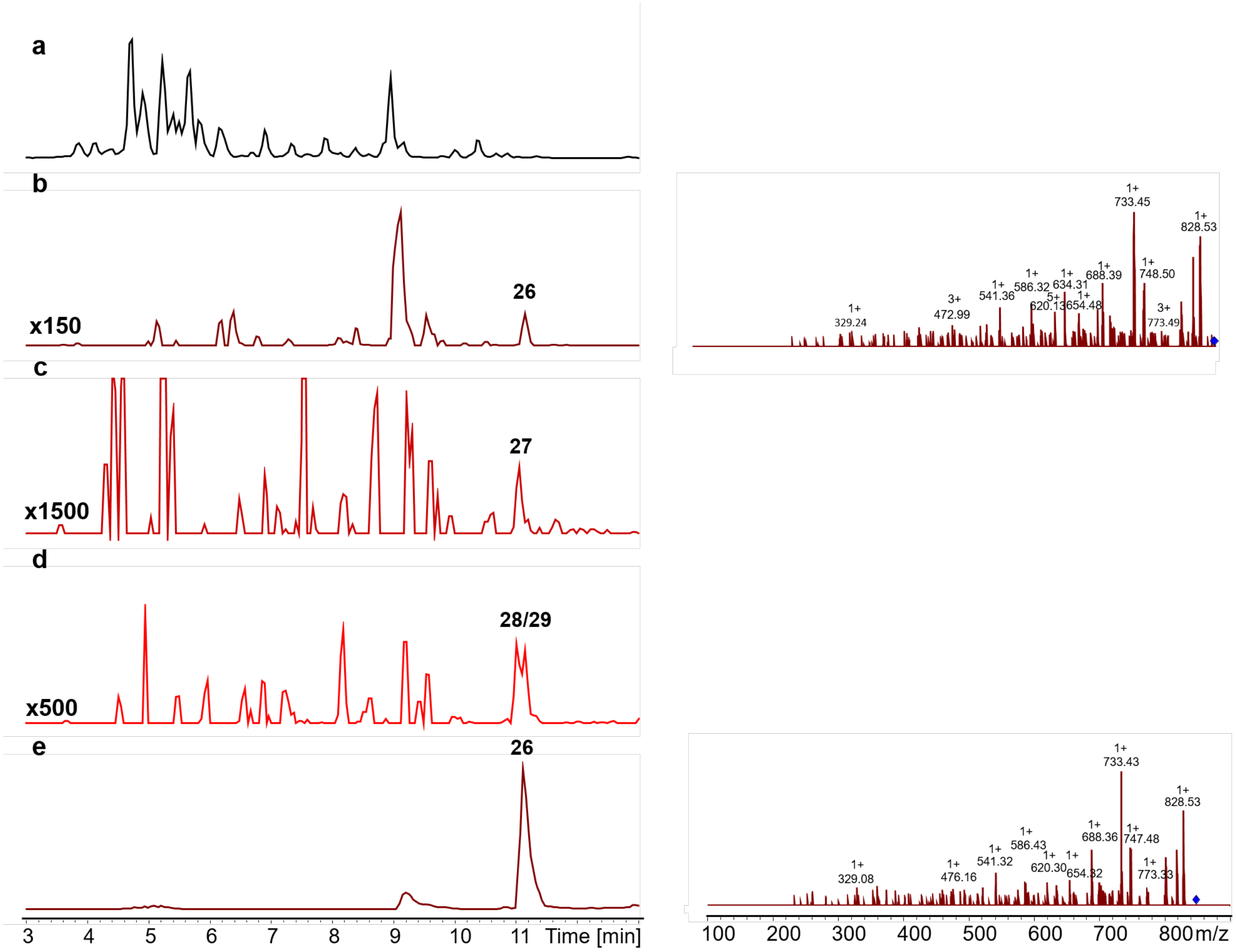

25.

26. **Figure S10. HPLC/MS data (Figure 1b) of compounds 26, 27 and 28/29 produced in *E. coli* DH10B::*mtaA*.** (a) BPC of an exemplary culture extract (b) EIC/MS$^2$ of **26** (*m/z* [M+H]$^+$ = 846.60, **NRPS-11**). (c) EIC data of **27** (*m/z* [M+H]$^+$ = 778.58, **NRPS-11**). (d) EIC data of **28/29** (*m/z* [M+H]$^+$ = 812.56, **NRPS-11**). (e) EIC/MS$^2$ of synthetic **26** (*m/z* [M+H]$^+$ = 846.60).

27. 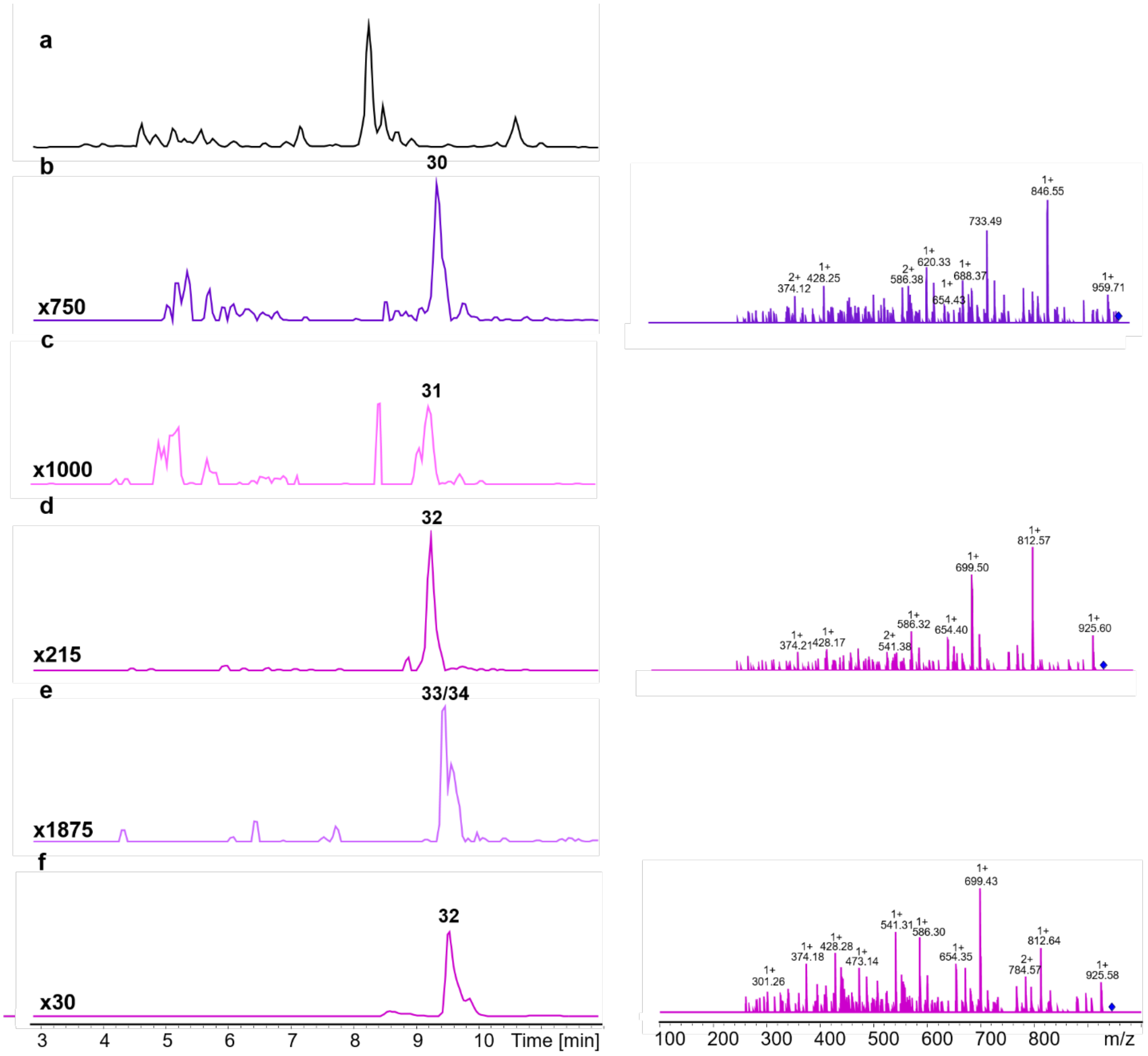

28. **Figure S11. HPLC/MS data (Figure 1b) of compounds 30, 31, 32 and 33/34 produced in *E. coli* DH10B::*mtaA*.** (a) BPC of an exemplary culture extract (b) EIC/MS$^2$ of **30** (*m/z* $[M+H]^+$ = 977.64, **NRPS-12**). (c) EIC data of **31** (*m/z* $[M+H]^+$ = 909.67, **NRPS-12**). (d) EIC data of **32** (*m/z* $[M+H]^+$ = 943.66, **NRPS-12**). (e) EIC data of **33/34** (*m/z* $[M+H]^+$ = 957.67, **NRPS-12**). (f) EIC/MS$^2$ of synthetic **32** (*m/z* $[M+H]^+$ = 943.66).

29.

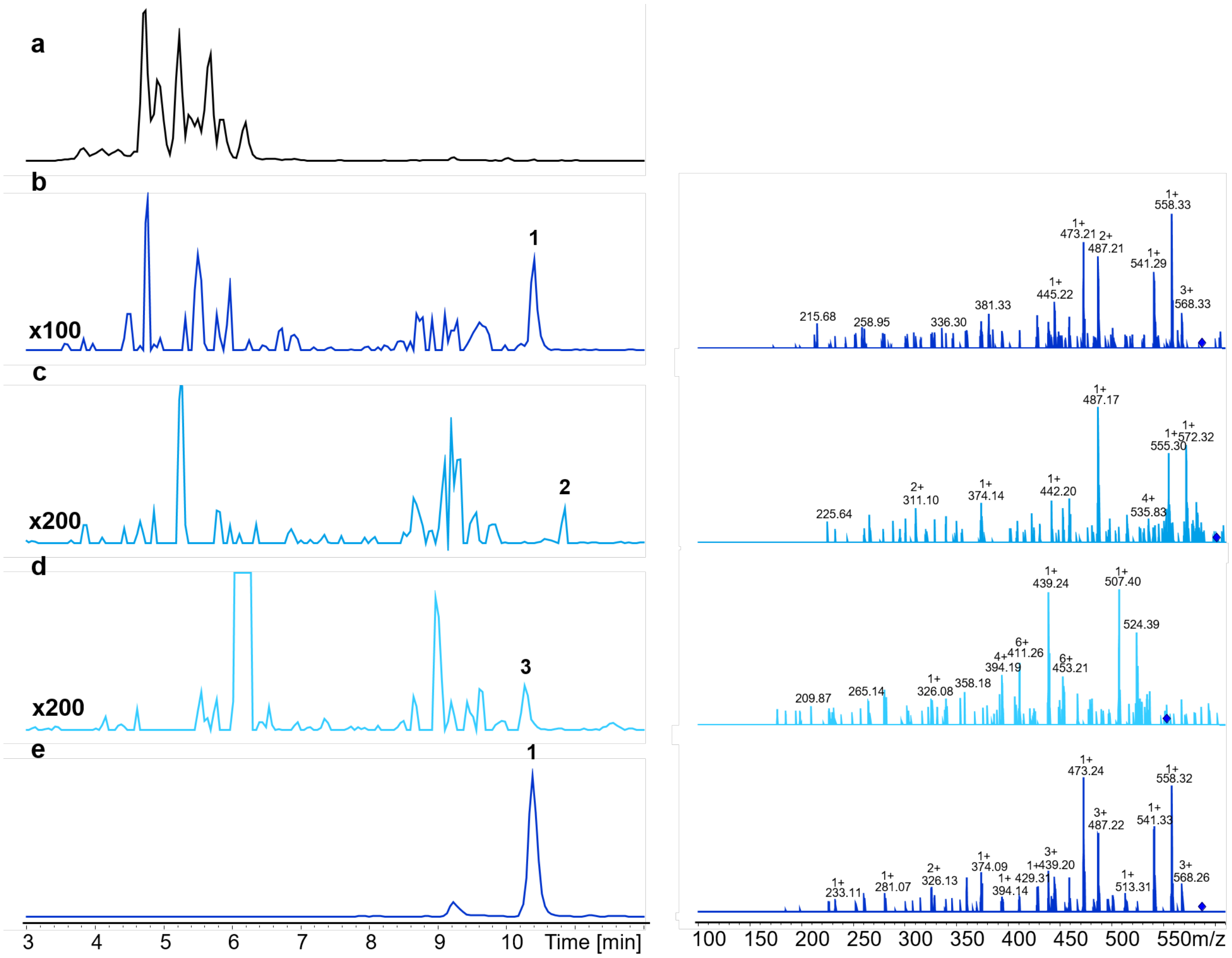

30. **Figure S12. HPLC/MS data (Figure 1c) of compounds 1, 2 and 3 produced in *E. coli* DH10B::*mtaA*.** (a) BPC of an exemplary culture extract (b) EIC/MS$^2$ of **1** (*m/z* $[M+H]^+$ = 586.40, **NRPS-13**). (c) EIC data of **2** (*m/z* $[M+H]^+$ = 600.41, **NRPS-13**). (d) EIC data of **3** (*m/z* $[M+H]^+$ = 552.41, **NRPS-13**). (e) EIC/MS$^2$ of synthetic **1** (*m/z* $[M+H]^+$ = 586.40).

31.

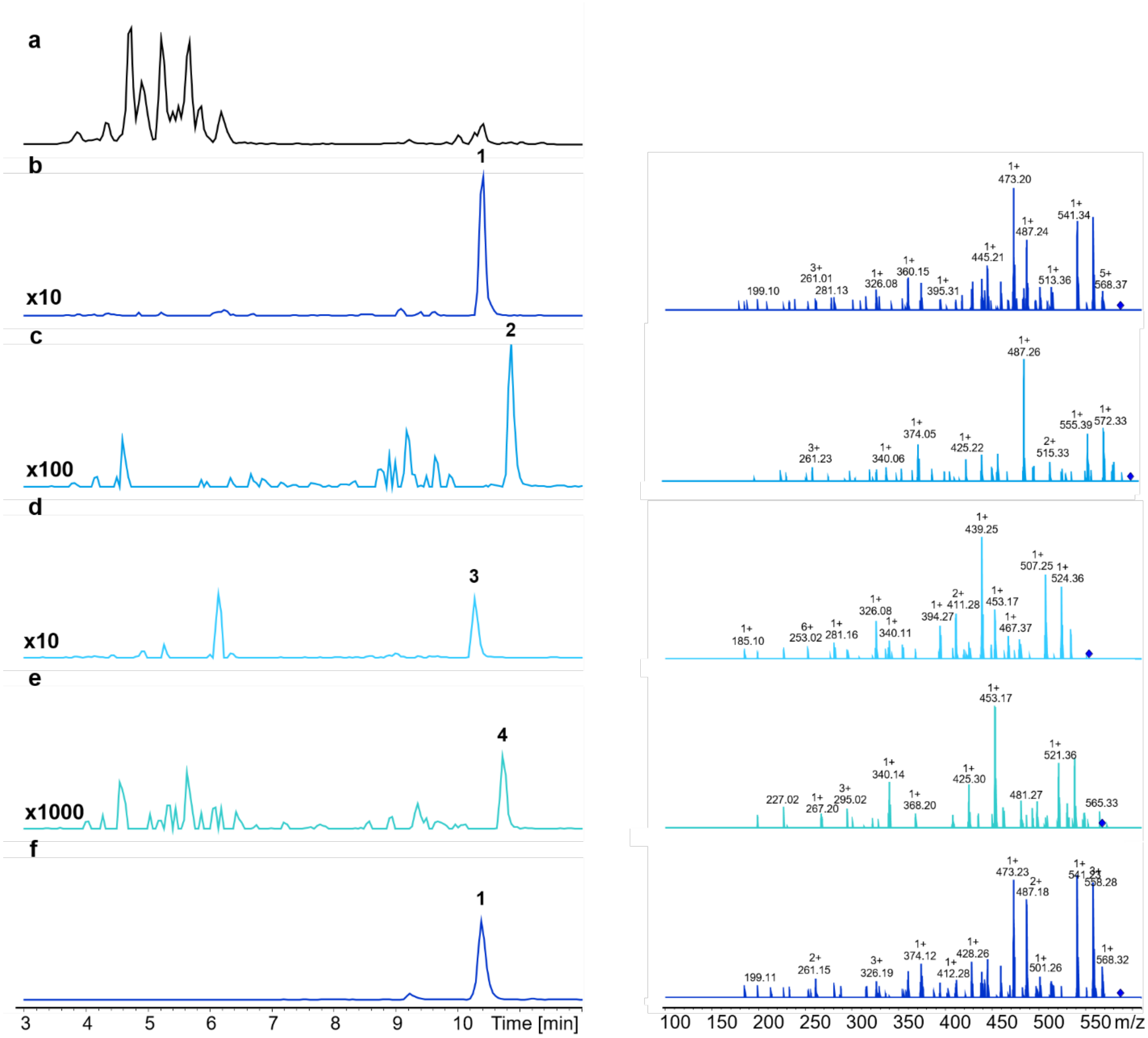

32. **Figure S13. HPLC/MS data (Figure 1c) of compounds 1, 2, 3 and 4 produced in *E. coli* DH10B::*mtaA*.** (a) BPC of an exemplary culture extract (b) EIC/MS$^2$ of **1** (*m/z* $[M+H]^+$ = 586.40, **NRPS-14**). (c) EIC data of **2** (*m/z* $[M+H]^+$ = 600.41, **NRPS-14**). (d) EIC data of **3** (*m/z* $[M+H]^+$ = 552.41, **NRPS-14**). (e) EIC data of **4** (*m/z* $[M+H]^+$ = 566.43, **NRPS-14**). (f) EIC/MS$^2$ of synthetic **1** (*m/z* $[M+H]^+$ = 586.40).

33. 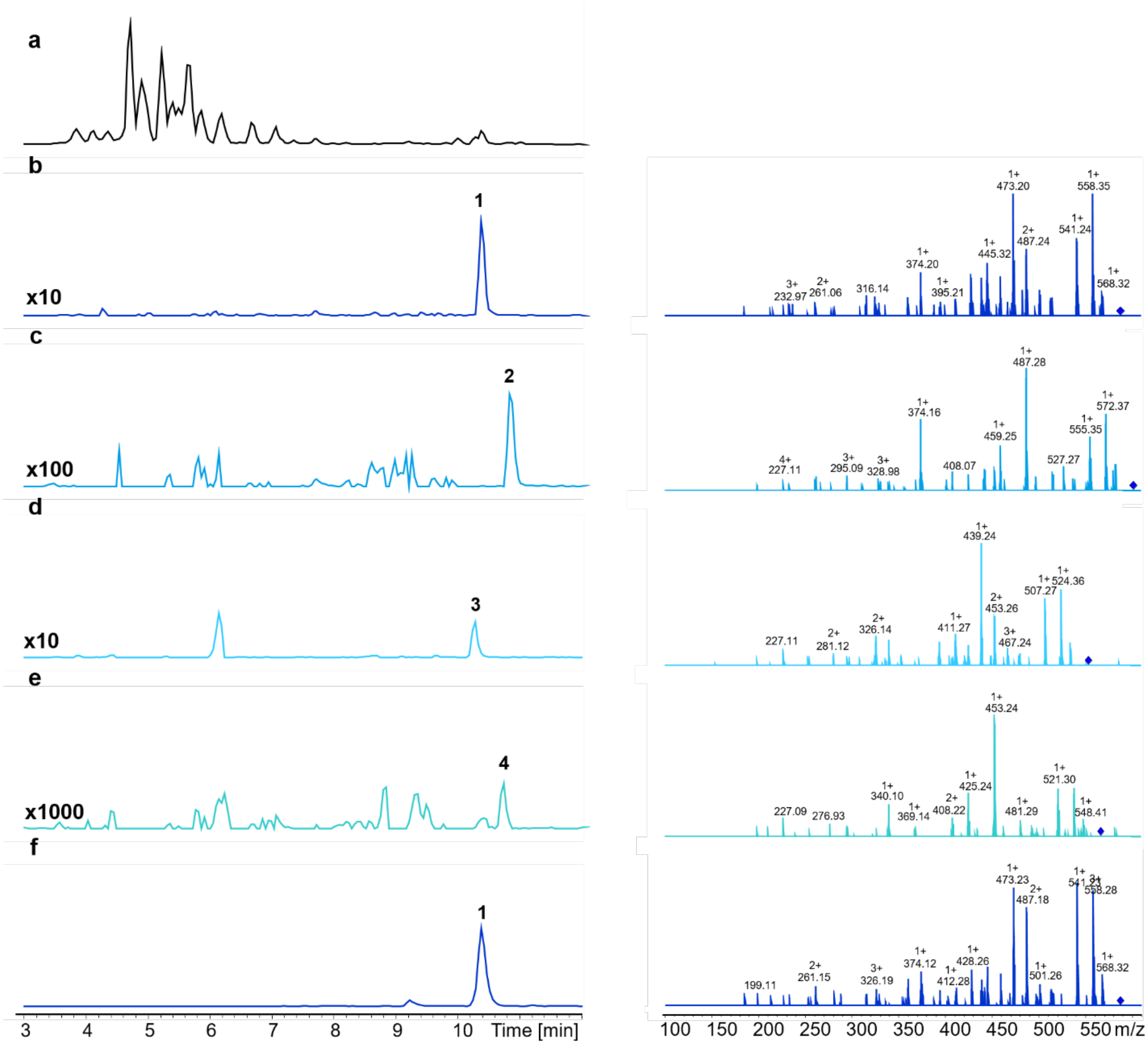

34. **Figure S14. HPLC/MS data (Figure 1c) of compounds 1, 2, 3 and 4 produced in *E. coli* DH10B::*mtaA*.** (a) BPC of an exemplary culture extract (b) EIC/MS$^2$ of **1** (*m/z* $[M+H]^+$ = 586.40, **NRPS-14**). (c) EIC data of **2** (*m/z* $[M+H]^+$ = 600.41, **NRPS-14**). (d) EIC data of **3** (*m/z* $[M+H]^+$ = 552.41, **NRPS-14**). (e) EIC data of **4** (*m/z* $[M+H]^+$ = 566.43, **NRPS-14**). (f) EIC/MS$^2$ of synthetic **1** (*m/z* $[M+H]^+$ = 586.40).

35. 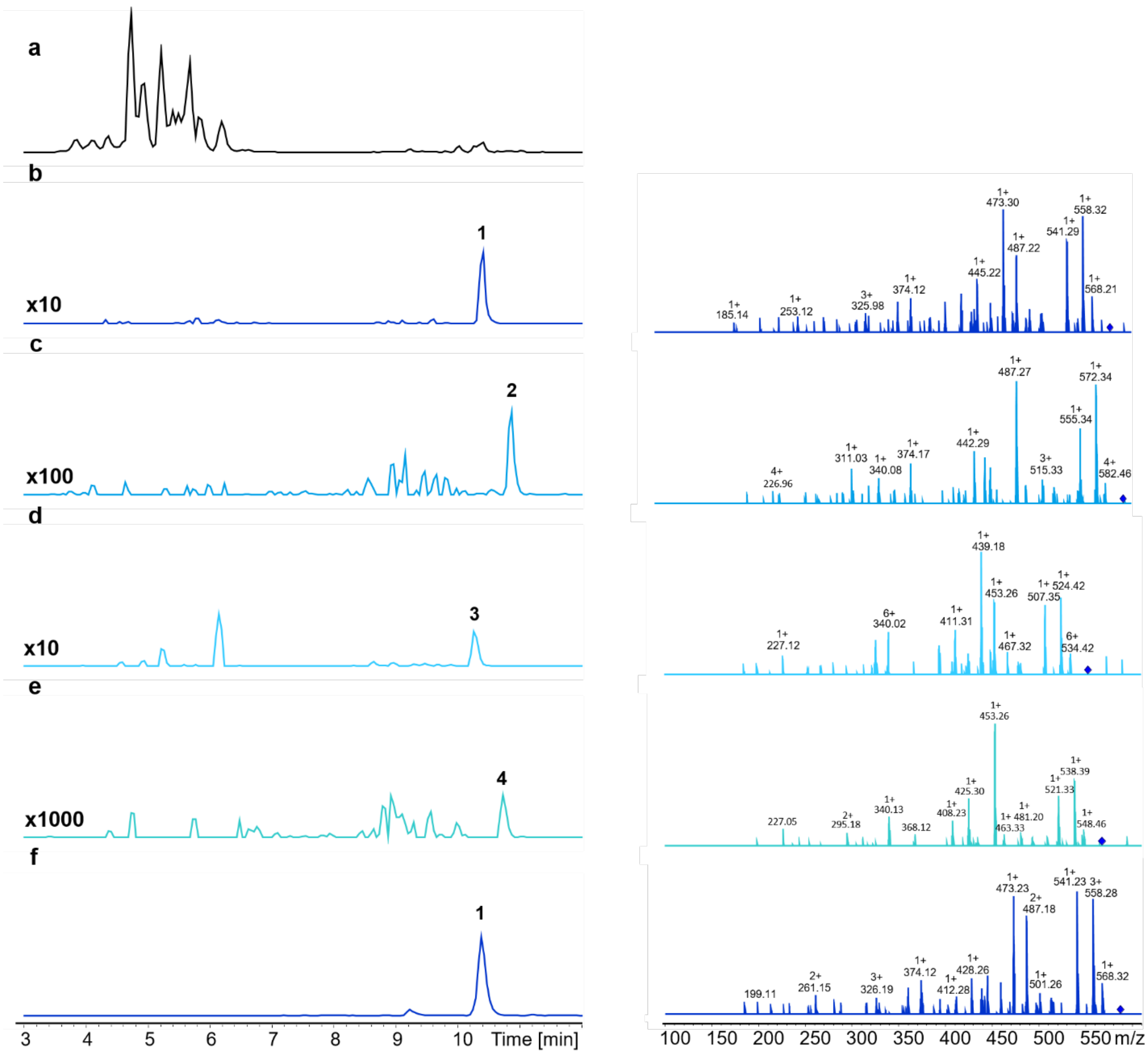

36. **Figure S15. HPLC/MS data (Figure 1c) of compounds 1, 2, 3 and 4 produced in *E. coli* DH10B::*mtaA*.** (a) BPC of an exemplary culture extract (b) EIC/MS$^2$ of **1** (*m/z* [M+H]$^+$ = 586.40, **NRPS-15**). (c) EIC data of **2** (*m/z* [M+H]$^+$ = 600.41, **NRPS-15**). (d) EIC data of **3** (*m/z* [M+H]$^+$ = 552.41, **NRPS-15**). (e) EIC data of **4** (*m/z* [M+H]$^+$ = 566.43, **NRPS-15**). (f) EIC/MS$^2$ of synthetic **1** (*m/z* [M+H]$^+$ = 586.40).

37.

38.

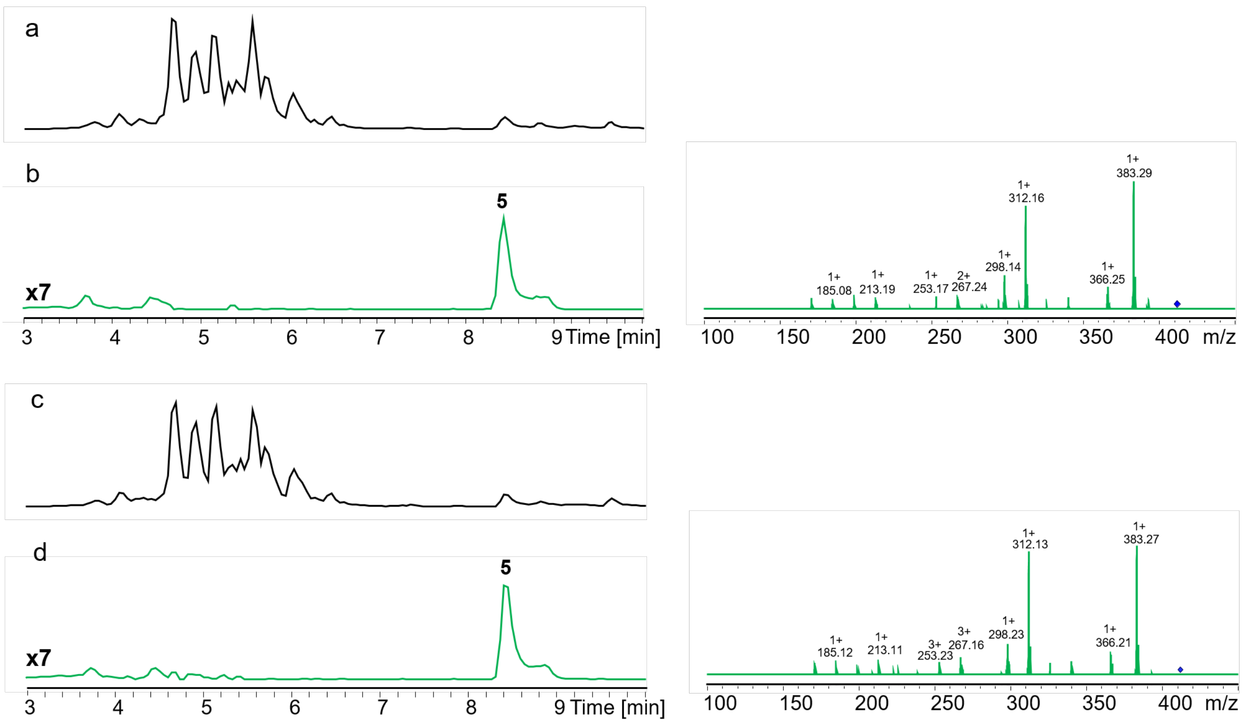

39. **Figure S16. HPLC/MS data refers to Figure 2 (NRPS-16, NRPS-17) of compound 5 produced in *E. coli* DH10B::*mtaA*.** (a) BPC of an exemplary culture extract (**NRPS-16**). (b) EIC/$MS^2$ data of **5** (*m/z* $[M+H]^+$ = 411.30, **NRPS-16**). (c) BPC of an exemplary culture extract (**NRPS-17**). (d) EIC/$MS^2$ data of **5** (*m/z* $[M+H]^+$ = 411.30, **NRPS-17**).

40. 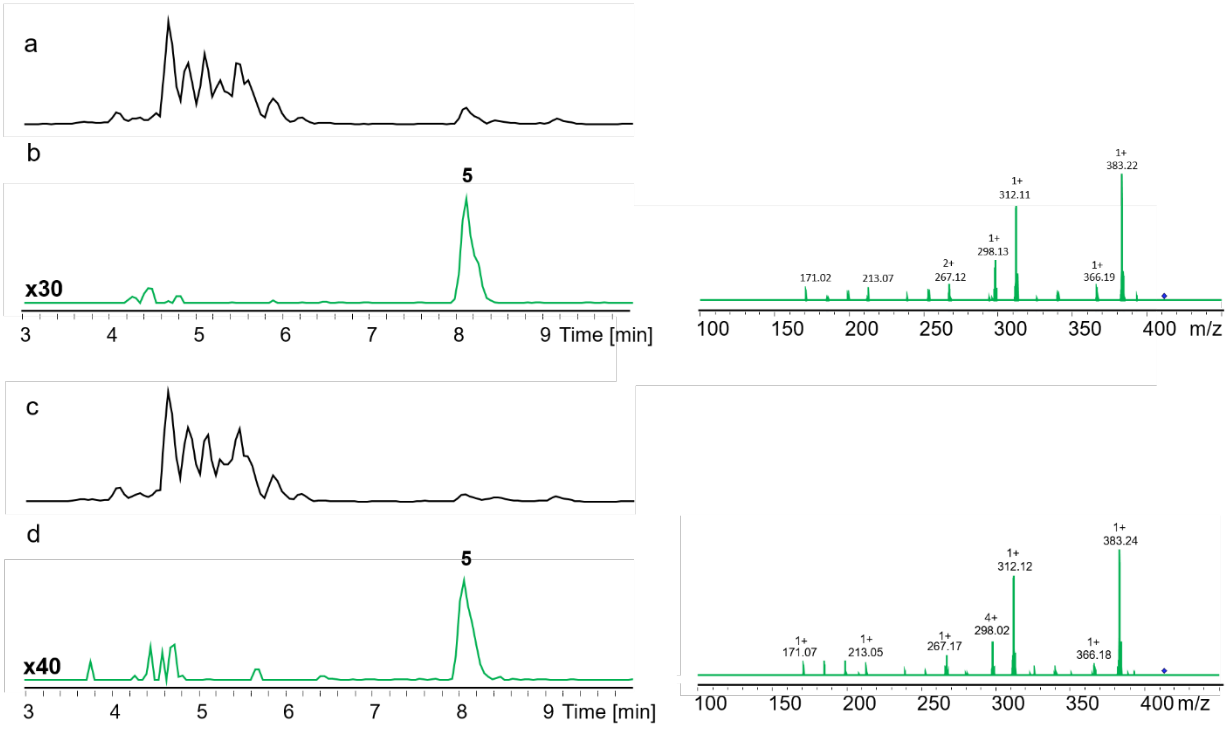

41. **Figure S17. HPLC/MS data refers to Figure 2 (NRPS-18, NRPS-19) of compound 5 produced in *E. coli* DH10B::*mtaA*.** (a) BPC of an exemplary culture extract (**NRPS-18**). (b) EIC/MS$^2$ data of **5** (*m/z* $[M+H]^+$ = 411.30, **NRPS-18**). (c) BPC of an exemplary culture extract (**NRPS-19**). (d) EIC/MS$^2$ data of **5** (*m/z* $[M+H]^+$ = 411.30, **NRPS-19**).

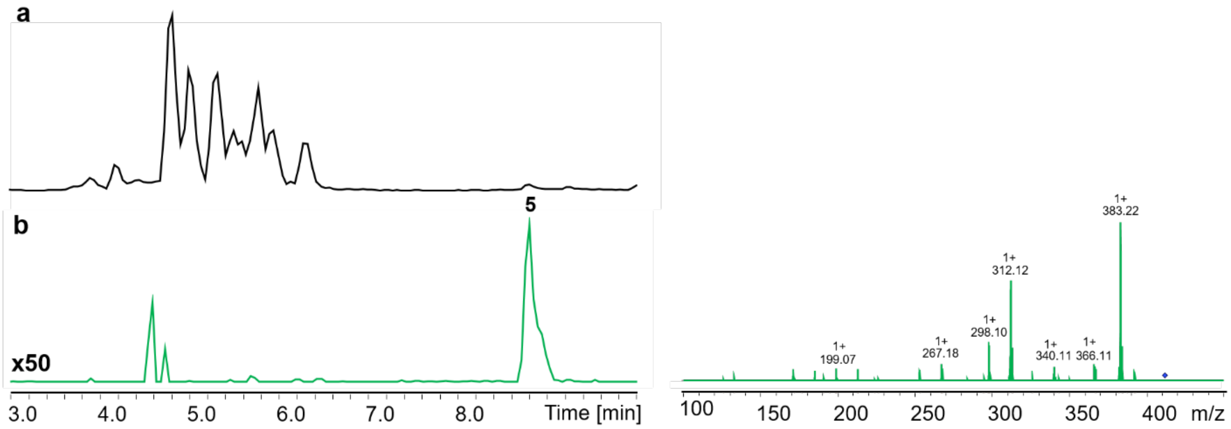

**Figure S18. HPLC/MS data refers to Figure 3 (NRPS-21) of compound 5 produced in *E. coli* DH10B::*mtaA*.** (a) BPC of an exemplary culture extract. (b) EIC/MS$^2$ data of **5** (*m/z* $[M+H]^+$ = 411.30).

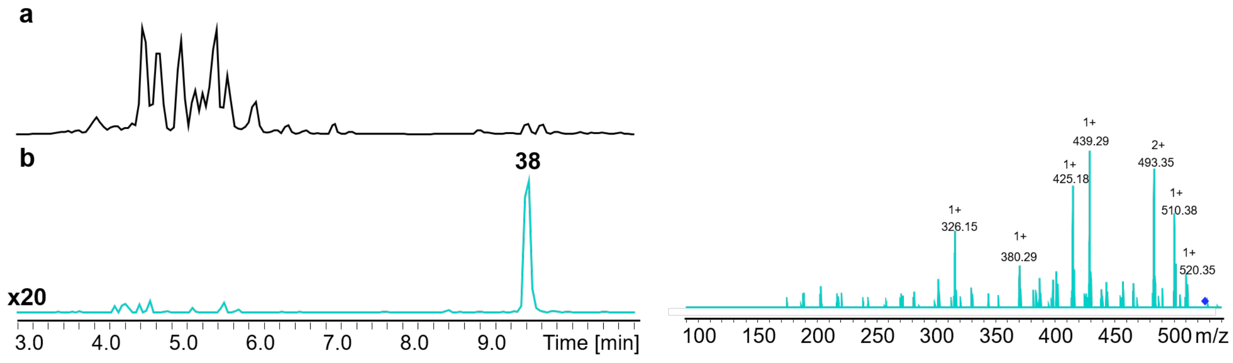

**Figure S19. HPLC/MS data refers to Figure 3 (NRPS-22) of compound 38 produced in E. coli DH10B::mtaA.** (a) BPC of an exemplary culture extract. (b) EIC/MS$^2$ data of **38** (m/z $[M+H]^+$ = 538.40).

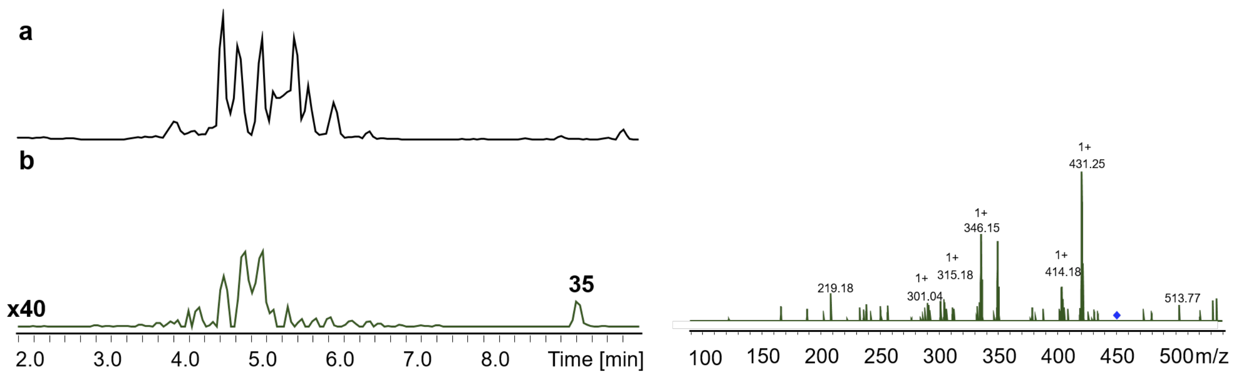

**Figure S20. HPLC/MS data refers to Figure 3 (NRPS-23) of compound 35 produced in *E. coli* DH10B::*mtaA*.** (a) BPC of an exemplary culture extract. (b) EIC/MS$^2$ data of **35** (*m/z* $[M+H]^+$ = 459.30).

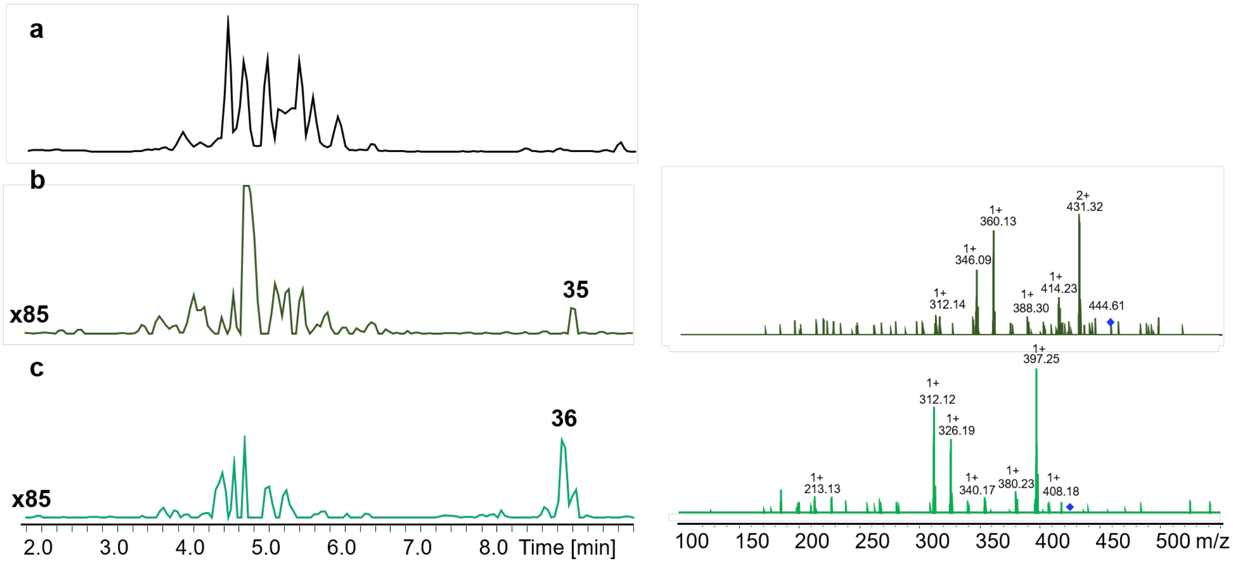

**Figure S21. HPLC/MS data refers to Figure 3 (NRPS-24) of compounds 35 and 36 produced in *E. coli* DH10B::*mtaA*.** (a) BPC of an exemplary culture extract. (b) EIC/MS$^2$ data of **35** (m/z $[M+H]^+$ = 459.30). (c) EIC/MS$^2$ data of **36** (m/z $[M+H]^+$ = 425.31).

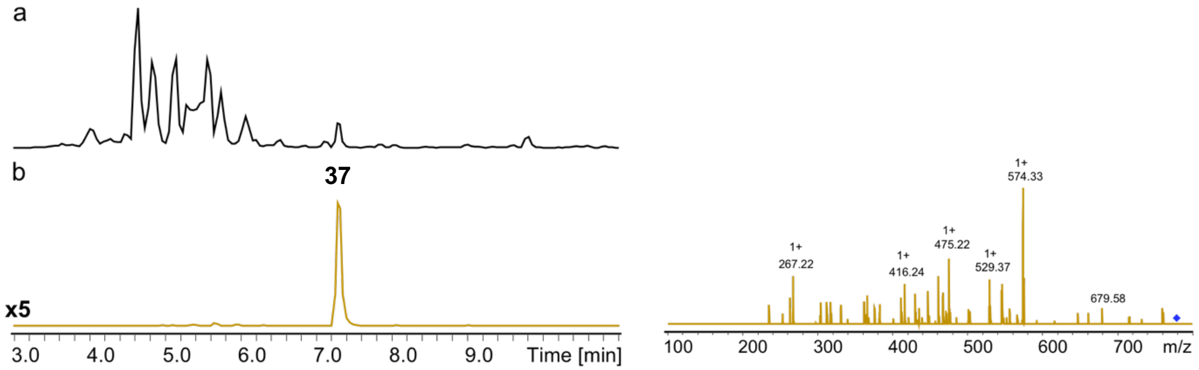

**Figure S22. HPLC/MS data refers to Figure 3 (NRPS-25) of compound 37 produced in *E. coli* DH10B::*mtaA*.** (a) BPC of an exemplary culture extract. (b) EIC/MS$^2$ data of **37** (*m/z* $[M+H]^+$ = 778.45).

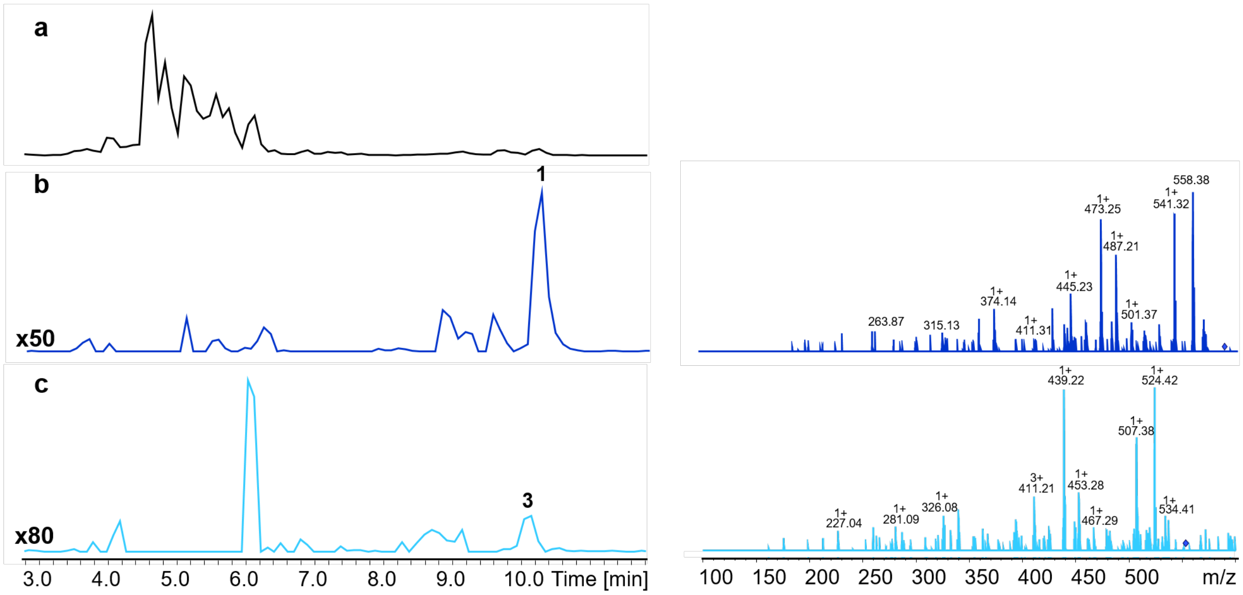

**Figure S23. HPLC/MS data refers to Figure 3 (NRPS-26) of compounds 1 and 3 produced in *E. coli* DH10B::*mtaA*.** (a) BPC of an exemplary culture extract. (b) EIC/$MS^2$ data of **1** (*m/z* $[M+H]^+$ = 586.40). (c) EIC/$MS^2$ data of **3** (*m/z* $[M+H]^+$ = 552.41).

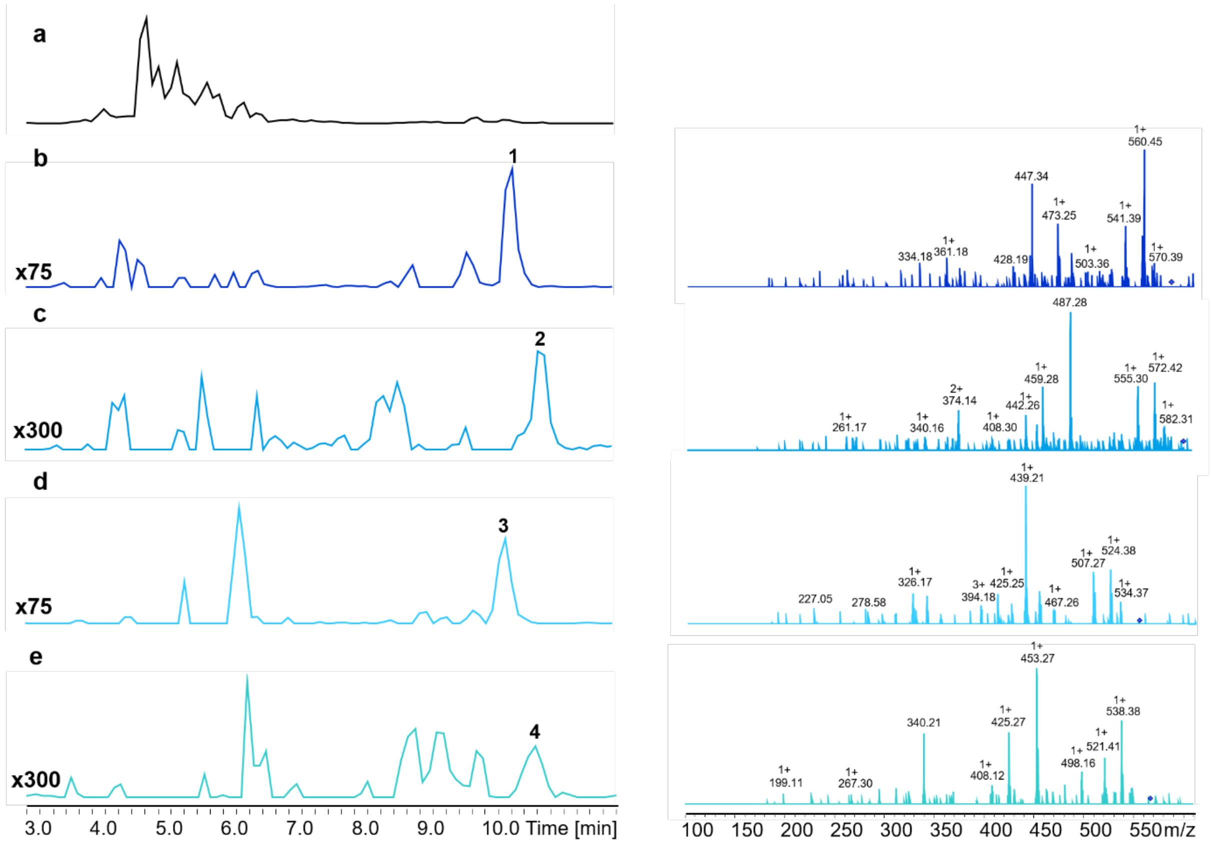

**Figure S24. HPLC/MS data refers to Figure 3 (NRPS-27) of compounds 1, 2, 3 and 4 produced in *E. coli* DH10B::*mtaA*.** (a) BPC of an exemplary culture extract. (b) EIC/MS$^2$ data of **1** (*m/z* $[M+H]^+$ = 586.40). (c) EIC/MS$^2$ data of **2** (*m/z* $[M+H]^+$ = 600.41). (d) EIC/MS$^2$ data of **3** (*m/z* $[M+H]^+$ = 552.41). (e) EIC/MS$^2$ data of **4** (*m/z* $[M+H]^+$ = 566.43).

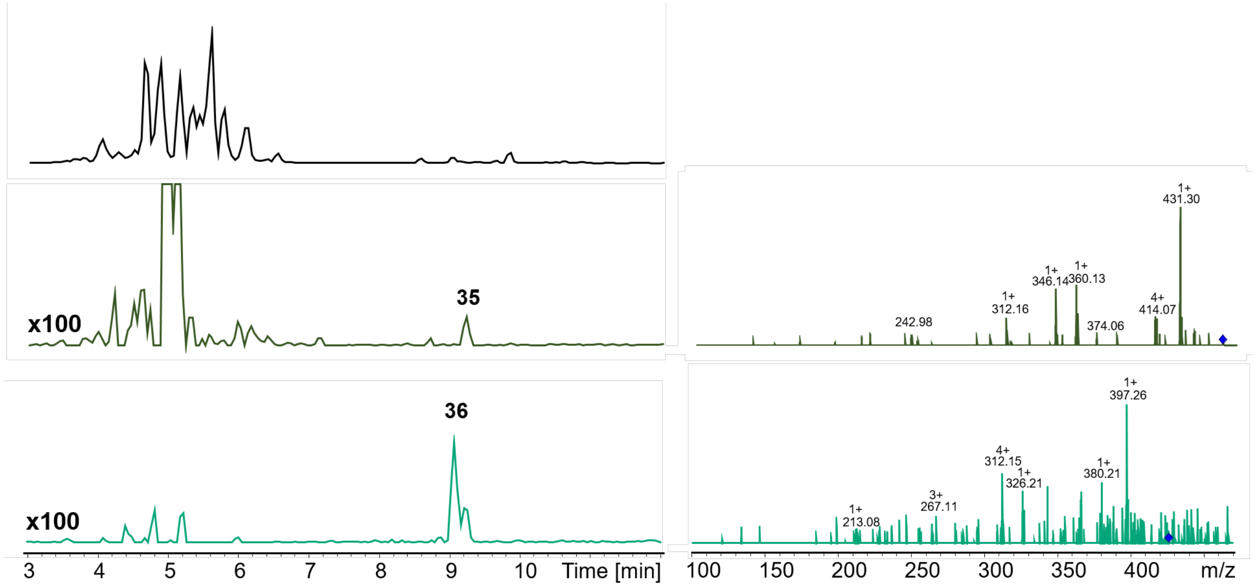

**Figure S25. HPLC/MS data refers to Figure 3 (NRPS-28) of compounds 35 and 36 produced in *E. coli* DH10B::*mtaA*.** (a) BPC of an exemplary culture extract. (b) EIC/MS$^2$ data of **35** ($m/z$ $[M+H]^+$ = 459.30). (c) EIC/MS$^2$ data of **36** ($m/z$ $[M+H]^+$ = 425.31).

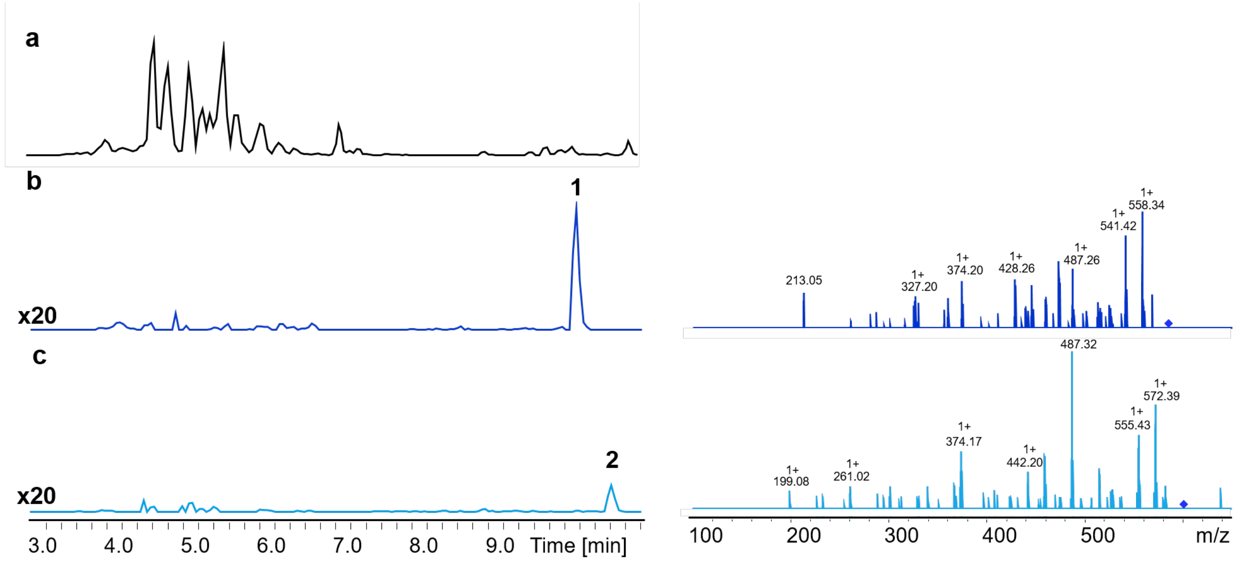

**Figure S26. HPLC/MS data refers to Figure 3 (NRPS-29) of compounds 1 and 2 produced in *E. coli* DH10B::*mtaA*.** (a) BPC of an exemplary culture extract. (b) EIC/MS$^2$ data of **1** ($m/z$ $[M+H]^+$ = 586.40). (c) EIC/MS$^2$ data of **2** ($m/z$ $[M+H]^+$ = 600.41).

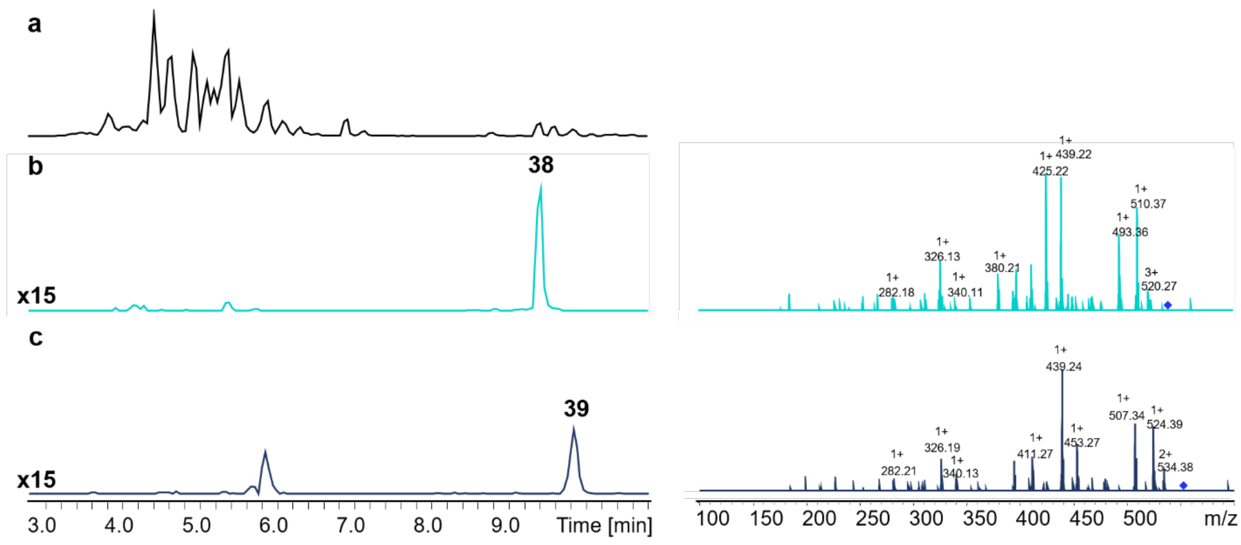

**Figure S27. HPLC/MS data refers to Figure 3 (NRPS-30) of compounds 38 and 39 produced in *E. coli* DH10B::*mtaA*.** (a) BPC of an exemplary culture extract. (b) EIC/MS$^2$ data of **38** (*m/z* $[M+H]^+$ = 588.40). (c) EIC/MS$^2$ data of **39** (*m/z* $[M+H]^+$ = 552.41).

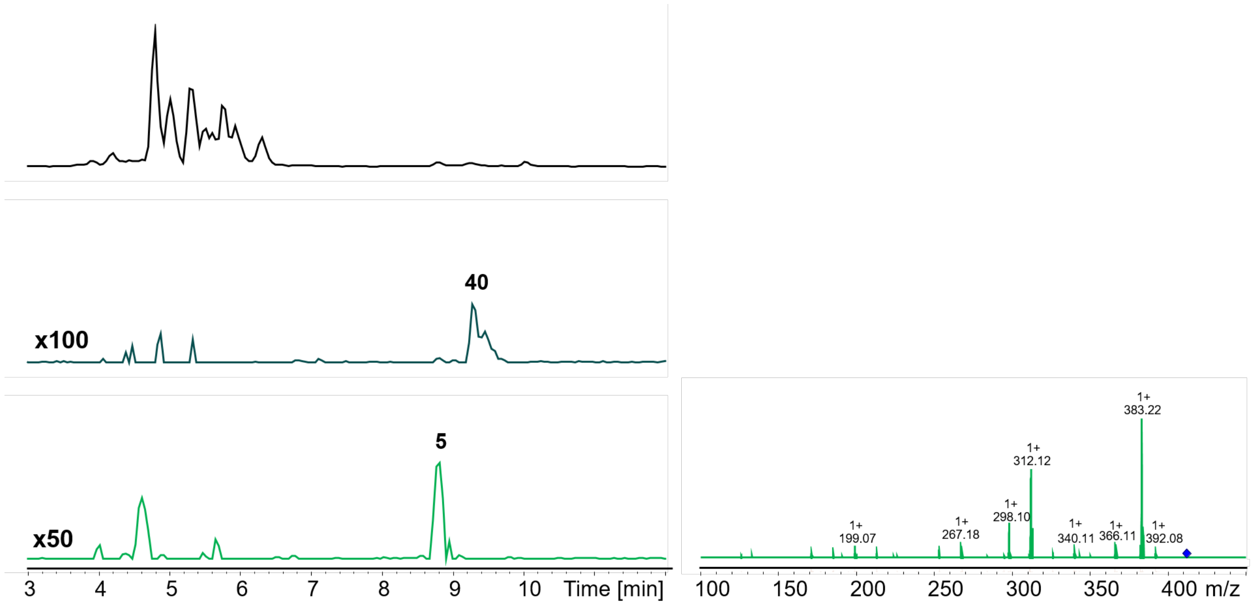

**Figure S28. HPLC/MS data refers to Figure 3 (NRPS-31) of compounds 40 and 5 produced in *E. coli* DH10B::*mtaA*.** (a) BPC of an exemplary culture extract. (b) EIC data of **40** (*m/z* $[M+H]^+$ = 425.31). (c) EIC/MS$^2$ data of **5** (m/z $[M+H]^+$ = 411.29).

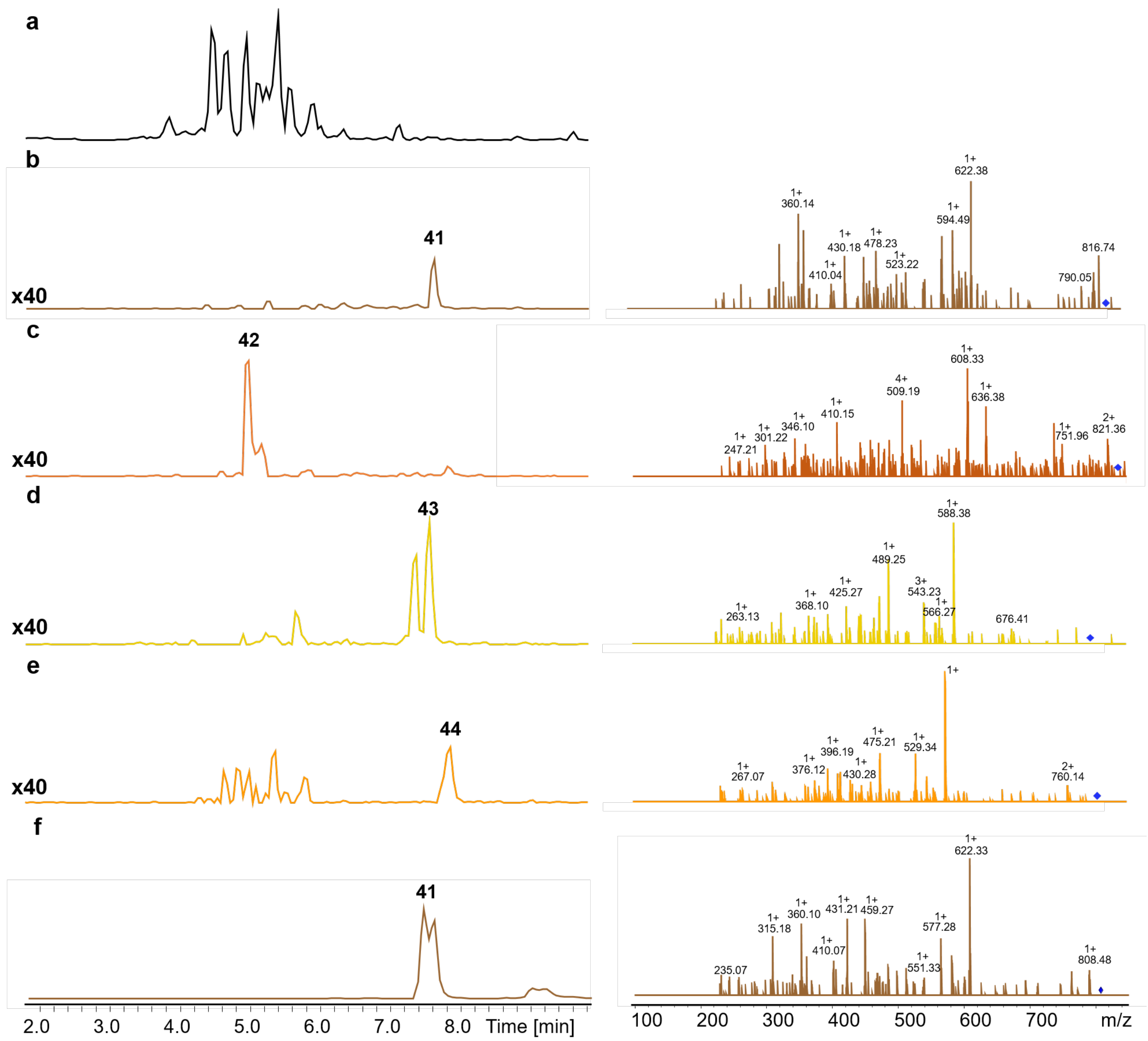

**Figure S29. HPLC/MS data refers to Figure 3 (NRPS-32) of compounds 41, 42, 43 and 44 produced in *E. coli* DH10B::*mtaA*.** (a) BPC of an exemplary culture extract. (b) EIC/MS$^2$ data of **41** (*m/z* $[M+H]^+$ = 826.45). (c) EIC/MS$^2$ data of **42** (*m/z* $[M+H]^+$ = 840.47). (d) EIC/MS$^2$ data of **43** (*m/z* $[M+H]^+$ = 792.47). (e) EIC/MS$^2$ data of **44** (*m/z* $[M+H]^+$ = 806.48). (f) EIC/MS$^2$ data of synthetic **41** (*m/z* $[M+H]^+$ = 826.45).

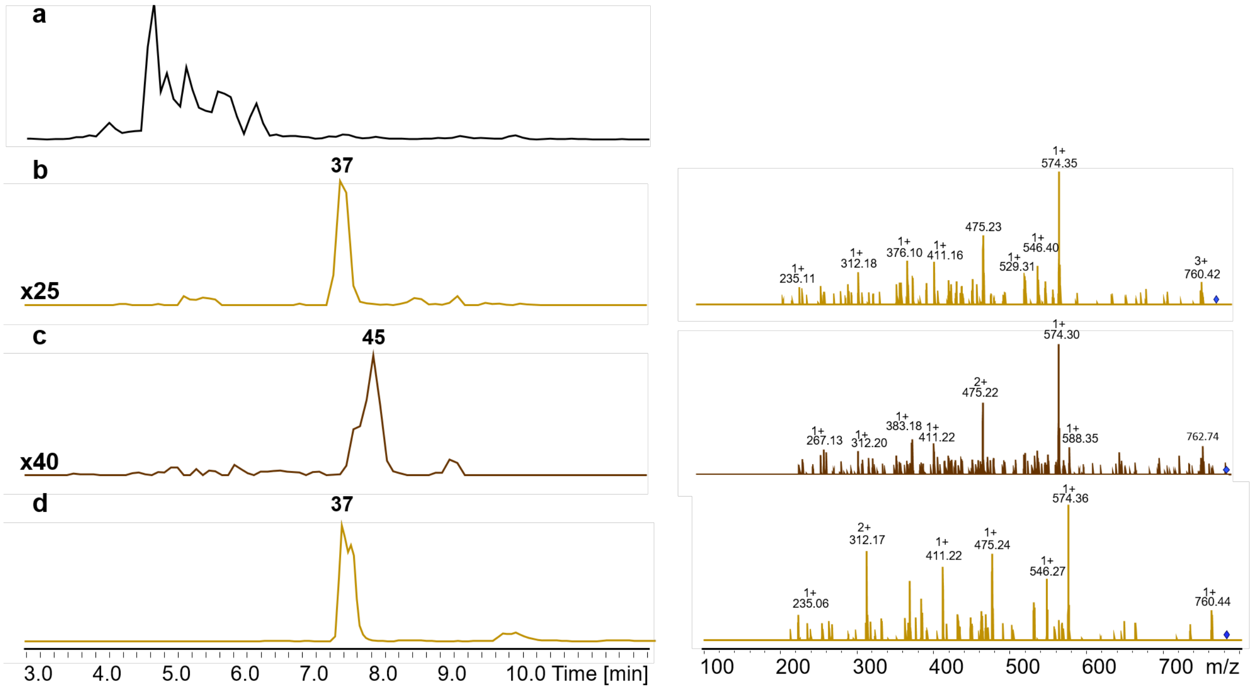

**Figure S30. HPLC/MS data refers to Figure 3 (NRPS-33) of compounds 37 and 45 produced in *E. coli* DH10B::*mtaA*.** (a) BPC of an exemplary culture extract. (b) EIC/MS$^2$ data of **37** (*m/z* $[M+H]^+$ = 778.45). (c) EIC/MS$^2$ data of **45** (*m/z* $[M+H]^+$ = 792.47). (d) EIC/MS$^2$ data of synthetic **37** (*m/z* $[M+H]^+$ = 778.45).

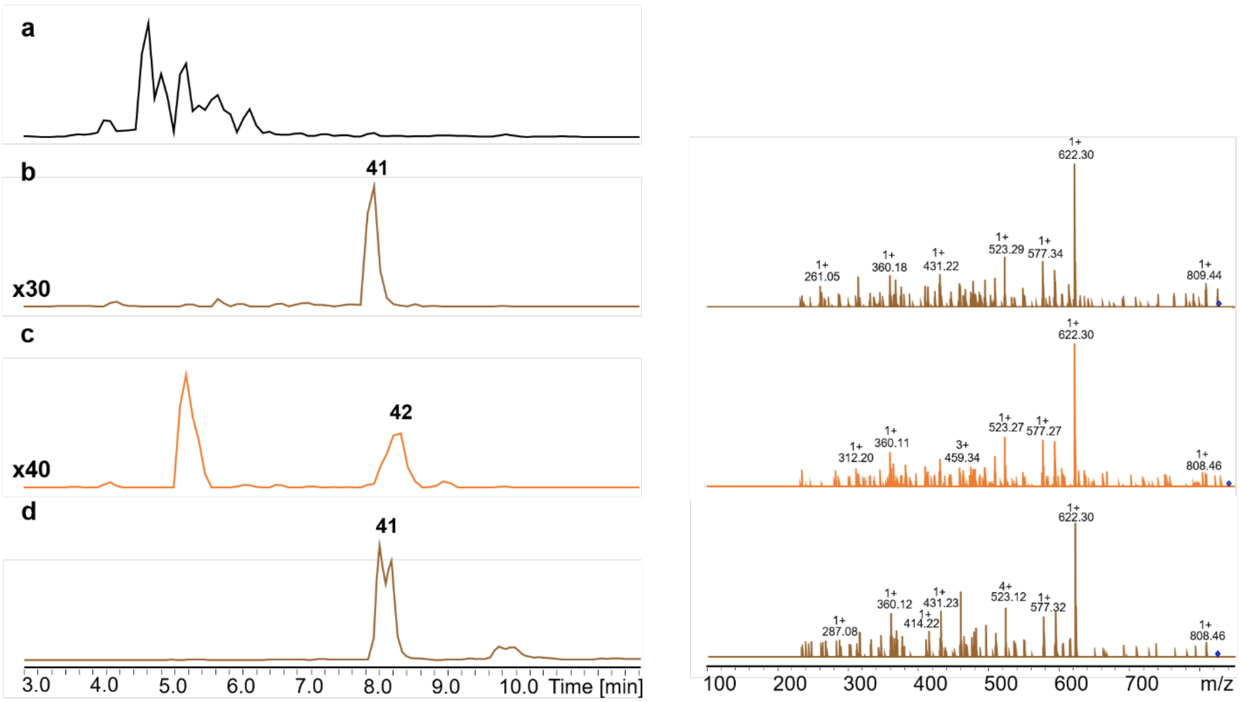

**Figure S31. HPLC/MS data refers to Figure 3 (NRPS-34) of compounds 41 and 42 produced in *E. coli* DH10B::*mtaA*.** (a) BPC of an exemplary culture extract. (b) EIC/MS$^2$ data of **41** (*m/z* [M+H]$^+$ = 826.45). (c) EIC/MS$^2$ data of **42** (*m/z* [M+H]$^+$ = 840.47). (d) EIC/MS$^2$ data of synthetic **41** (*m/z* [M+H]$^+$ = 826.45).

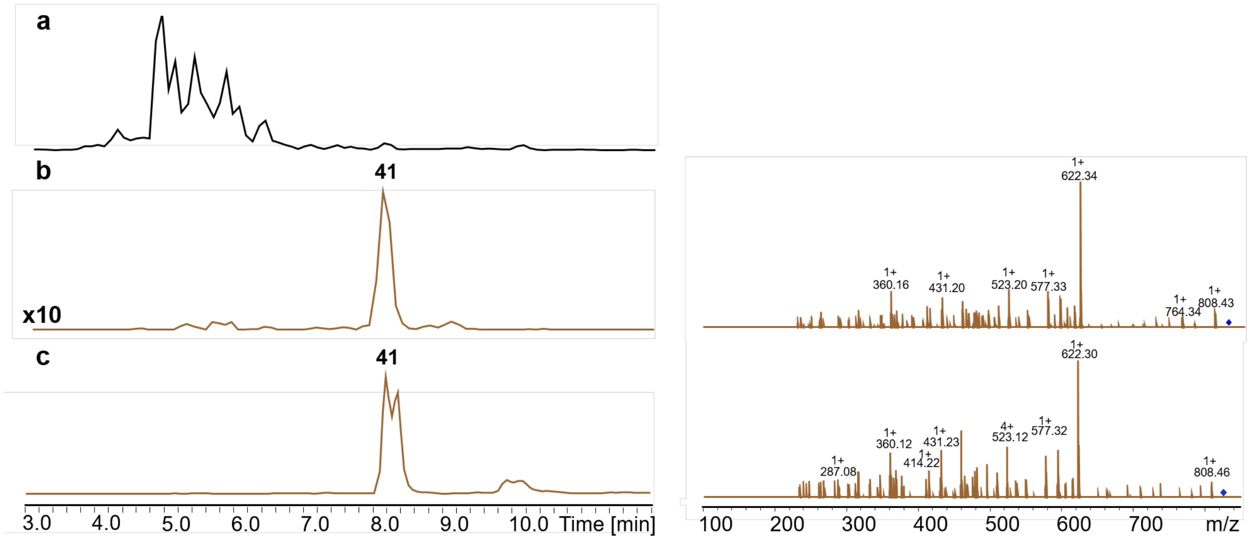

**Figure S32. HPLC/MS data refers to Figure 3 (NRPS-35) of compound 41 produced in *E. coli* DH10B::*mtaA*.** (a) BPC of an exemplary culture extract. (b) EIC/MS$^2$ data of **41** (*m/z* [M+H]$^+$ = 826.45). (c) EIC/MS$^2$ data of synthetic **41** (*m/z* [M+H]$^+$ = 826.45).

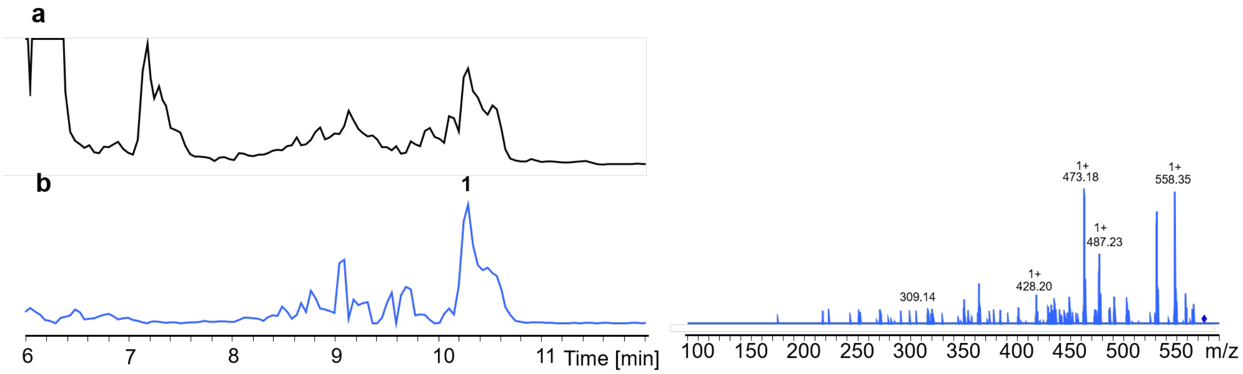

**Figure S33. HPLC/MS data refers to Figure 3 (NRPS-36) of compound 1 produced in *E. coli* DH10B::*mtaA*.** (a) BPC of an exemplary culture extract. (b) EIC/MS$^2$ data of **1** ($m/z$ $[M+H]^+$ = 586.40).

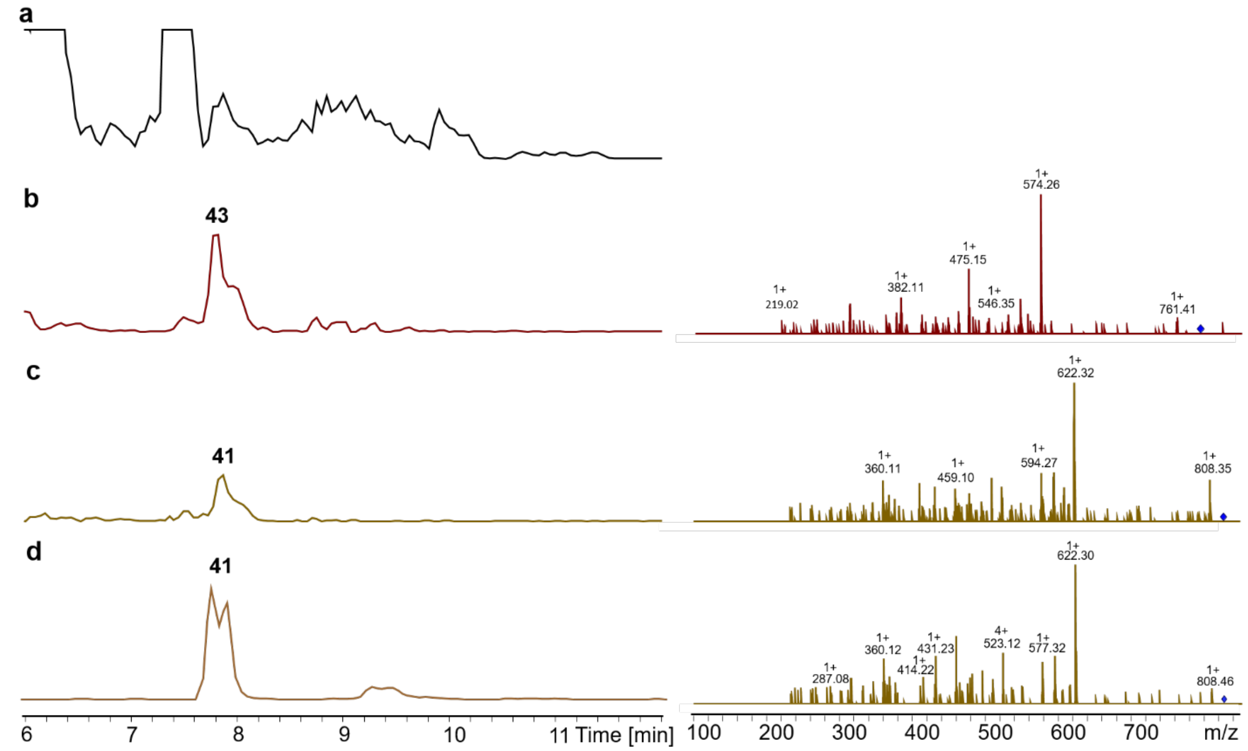

**Figure S34. HPLC/MS data refers to Figure 3 (NRPS-37) of compounds 43 and 41 produced in *E. coli* DH10B::*mtaA*.** (a) BPC of an exemplary culture extract. (b) EIC/MS$^2$ data of **43** ($m/z$ $[M+H]^+$ = 792.47). (c) EIC/MS$^2$ data of **41** ($m/z$ $[M+H]^+$ = 826.45). (d) EIC/MS$^2$ data of synthetic **41** ($m/z$ $[M+H]^+$ = 826.45).

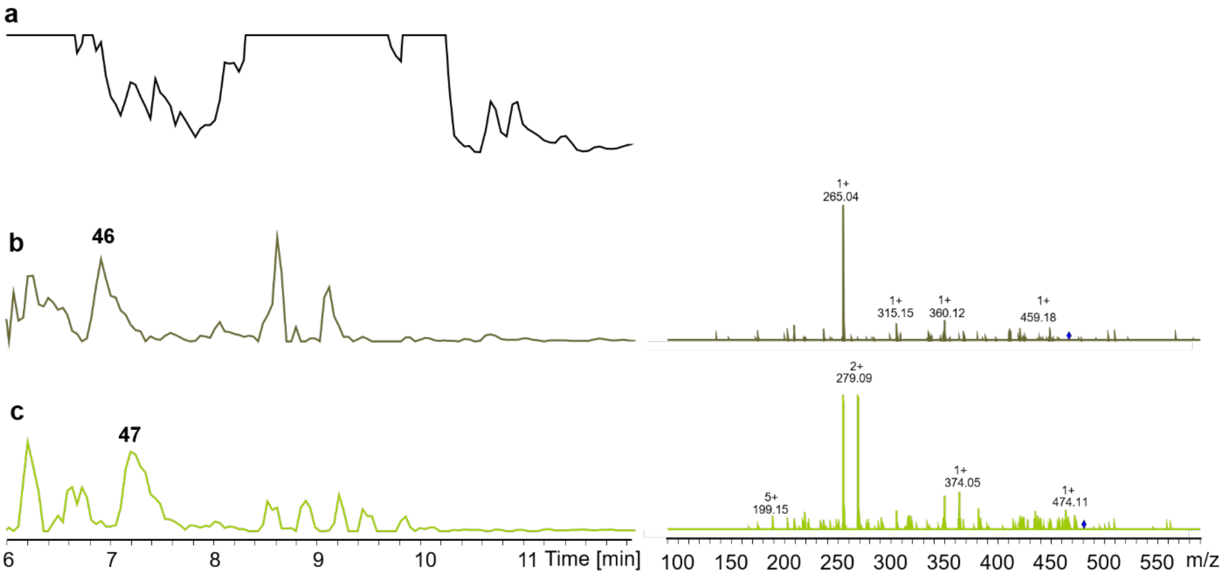

**Figure S35. HPLC/MS data refers to Figure 3 (NRPS-38) of compounds 46 and 47 produced in *E. coli* DH10B::*mtaA*.** (a) BPC of an exemplary culture extract. (b) EIC/$MS^2$ data of **46** (*m/z* $[M+H]^+$ = 476.30). (c) EIC/$MS^2$ data of **47** (*m/z* $[M+H]^+$ = 490.32).

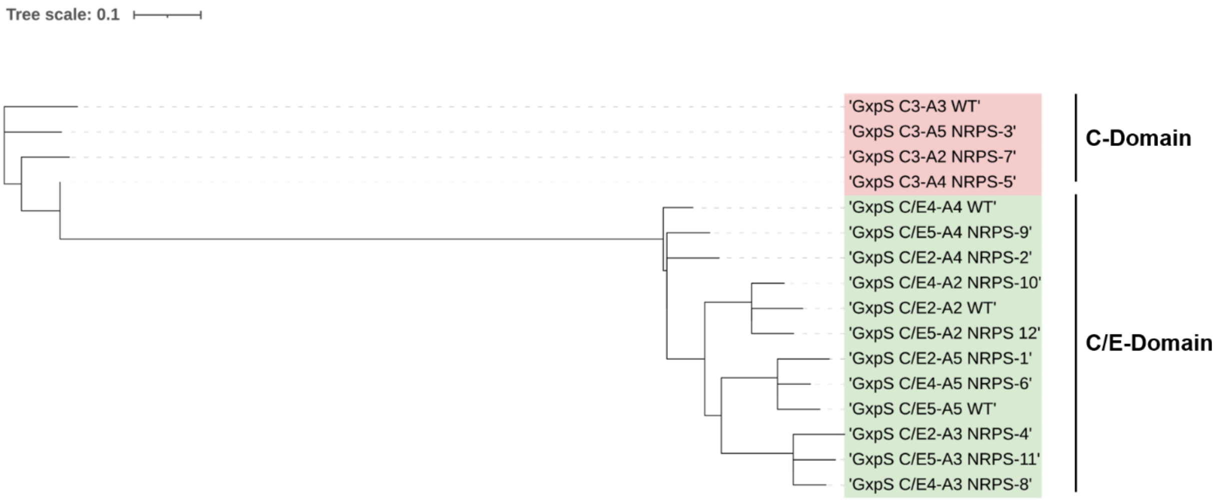

**Figure S36. Phylogenetic three of extracted C-A and C/E-A didomains from GxpS.** WT C-A and C/E-A didomains were sequenced aligned with non-natural C-A and C/E-A didomains from NRPS-1 to -16. Alignment was used to build a three applying RAxML. Results show a clear clustering of the C-domains into C- and C/E indicating the importance of C-domain types. A-domain specificity clustering only becomes clear on the second level. C-Domain type did not change for NRPS-9, -2, -10, -12, -1 and -6. C-domain Changes from C to C/E applies for NRPS-4, -11 and -8 and changes from C/E to C applies for NRPS-3, -7 and -5.

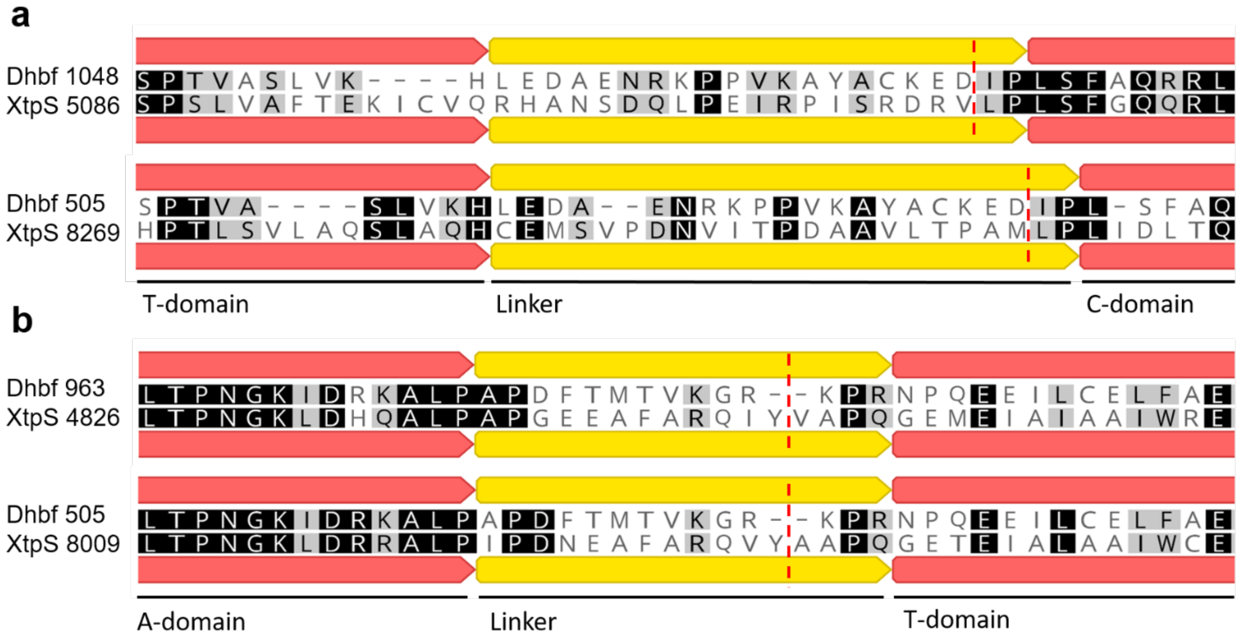

**Figure S37. Sequence alignments of XtpS linker sequences.** (a) XtpS $T_2$-$C_2$ (XtpS 5086) and $T_3$-$C_3$ (XtpS 8269) were aligned to the linker region excised from Dhbf crystal structure (Protein Database ID: 5U89). (b) XtpS $A_2$-$T_2$ (XtpS 4826) and $A_3$-$T_3$ (XtpS 8009) were aligned to the linker region excised from Dhbf crystal structure. Red dashed lines indicate SYNZIP insertion points.

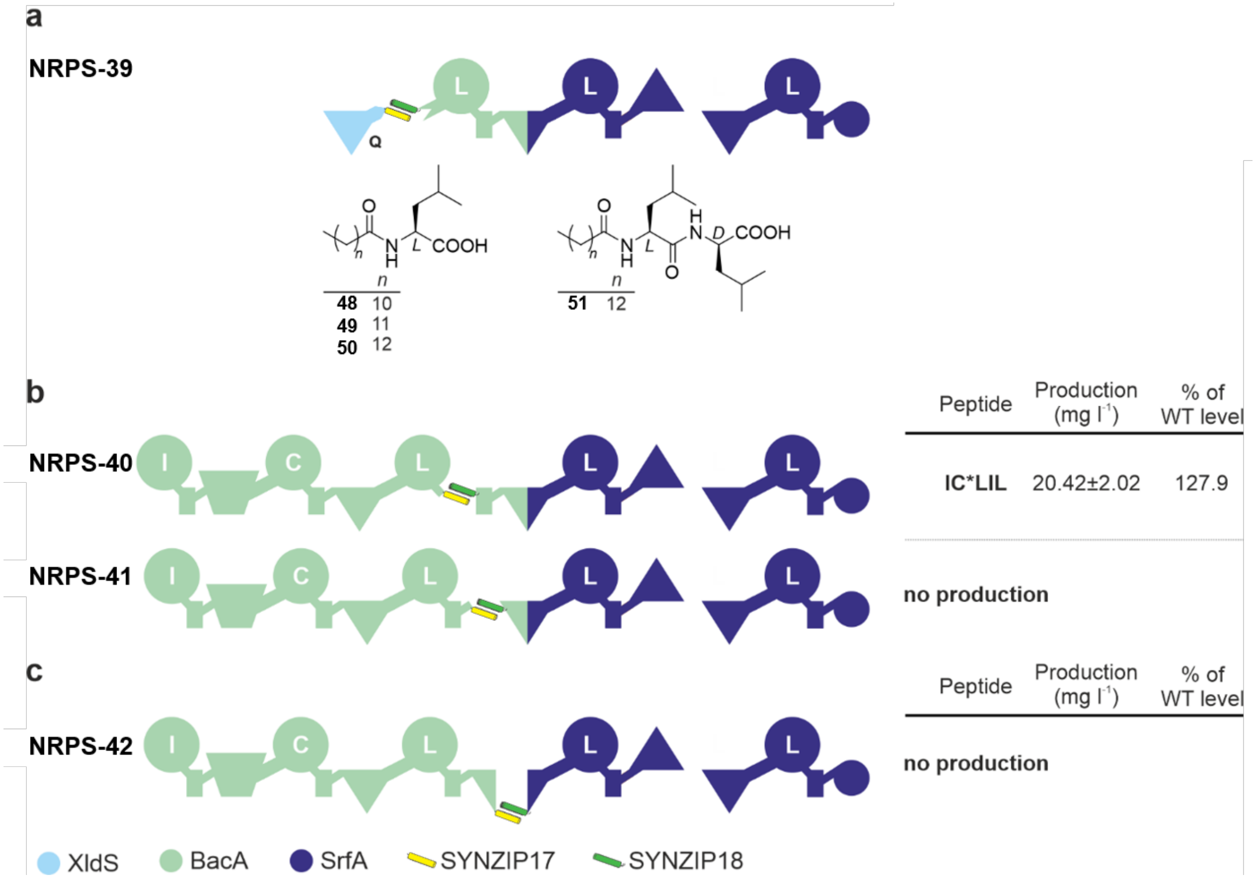

| Peptide | Production (mg l⁻¹) | % of WT level |
|---|---|---|
| IC*LIL | 20.42±2.02 | 127.9 |
| no production | | |

| Peptide | Production (mg l⁻¹) | % of WT level |
|---|---|---|
| no production | | |

**Figure S38.** Additional examples of bipartite type S NRPS split in between and within RtpS modules[1].

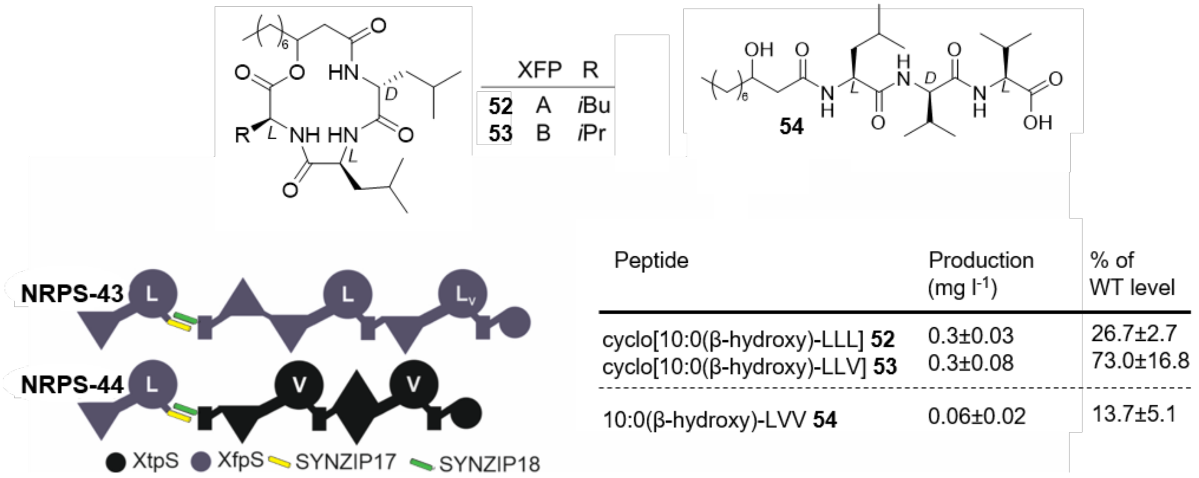

| Peptide | Production (mg l⁻¹) | % of WT level |
|---|---|---|
| cyclo[10:0(β-hydroxy)-LLL] **52** | 0.3±0.03 | 26.7±2.7 |
| cyclo[10:0(β-hydroxy)-LLV] **53** | 0.3±0.08 | 73.0±16.8 |
| 10:0(β-hydroxy)-LVV **54** | 0.06±0.02 | 13.7±5.1 |

**Figure S39.** Additional examples of bipartite type S NRPS split within modules of xefoampeptide- producing NRPS (XfpS)[8] and between XfpS and XtpS.

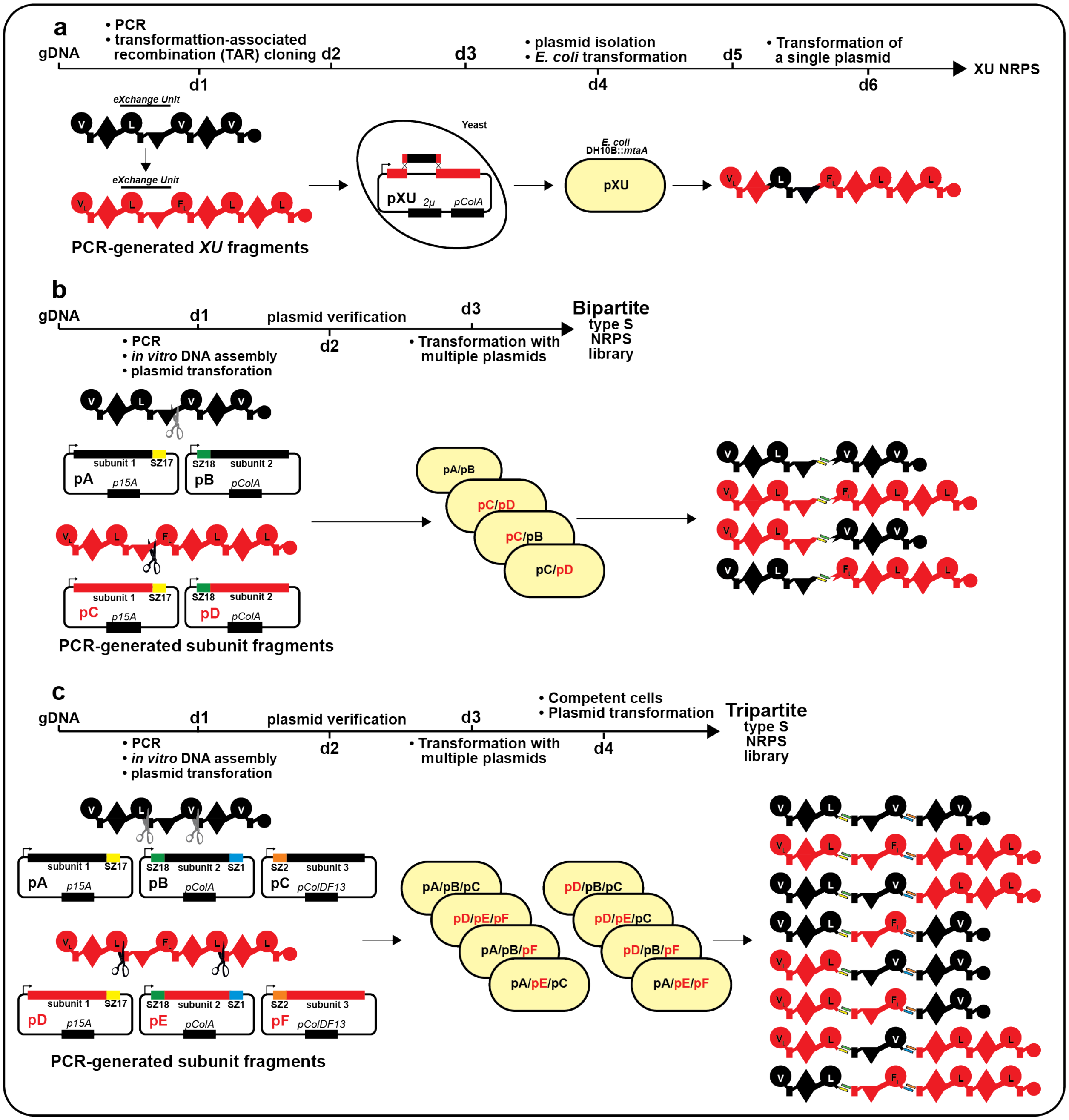

**Figure S40. Extended comparison of workflows to generate chimeric NRPSs.** (a) “Classic” Yeast-based TAR cloning approach. Here, two BGCs were used to generate one chimeric NRPS applying the XU concept. Ideally, cloning of one chimeric NRPS takes 6 days. (b) Generation of bipartite typeS NRPS libraries. Here, two BGCs were split within the XU concept fusion site and generated subunits 1 and 2 were assembled into two distinct plasmid BBs carrying differing ORIs and resistance cassettes. Subunit 1 plasmids and subunit 2 plasmids were co-transformed into *E. coli* DH10B::*mtaA* in different combinations resulting in four typeS NRPSs. Cloning can be achieved in only half the days (3 days) compared to TAR cloning. Furthermore, throughput of recombinant NRPS can be increased exponentially with the number of building blocks. (c) Generation of tripartite typeS NRPS libraries. Here, two BGCs were split twice within the A-T linker region. Subunits 1-3 were assembled into three distinct plasmids carrying differing ORIs and resistance cassettes. Subunit 1, subunit 2 und subunit 3 plasmids were co-transformed into *E. coli*

DH10B::*mtaA* in different combinations. Cloning ca be archived within 4 days. Throughput increases more rapidly compared to bipartite typeS NRPS.

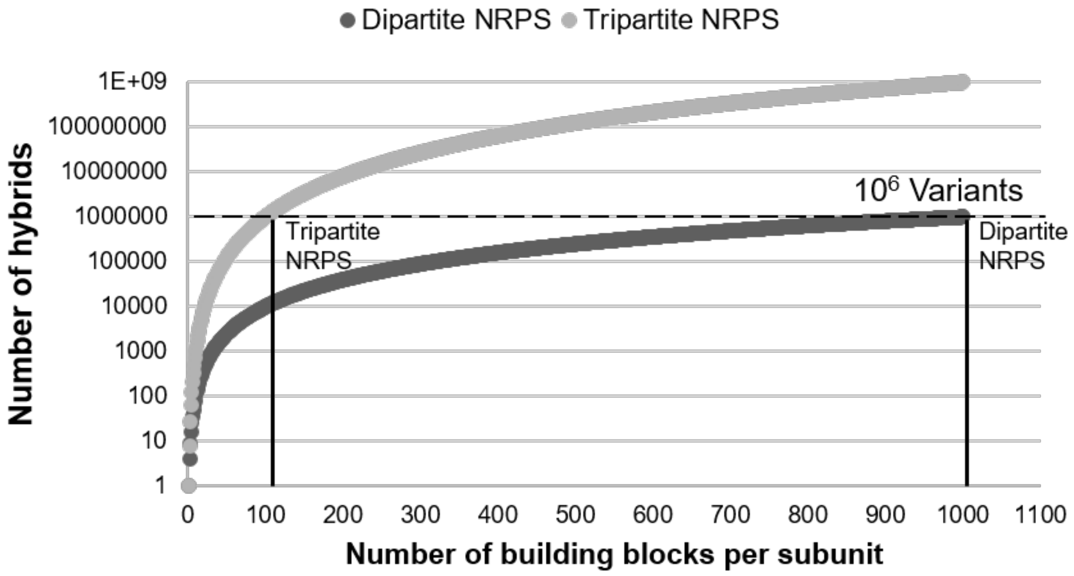

**Figure S41. Comparing the throughput between di- and tripartite NRPSs.** Number of hybrids increases exponentially with the number if building blocks per subunit for both di- and tripartite NRPSs. The number of hybrids increases more rapidly for dipartite NRPS. To build a library with $10^6$ variants, one requires 100 building blocks per subunit (in total 2000) for dipartite NRPS and only 100 building blocks per subunit (in total 300) for tripartite NRPS.

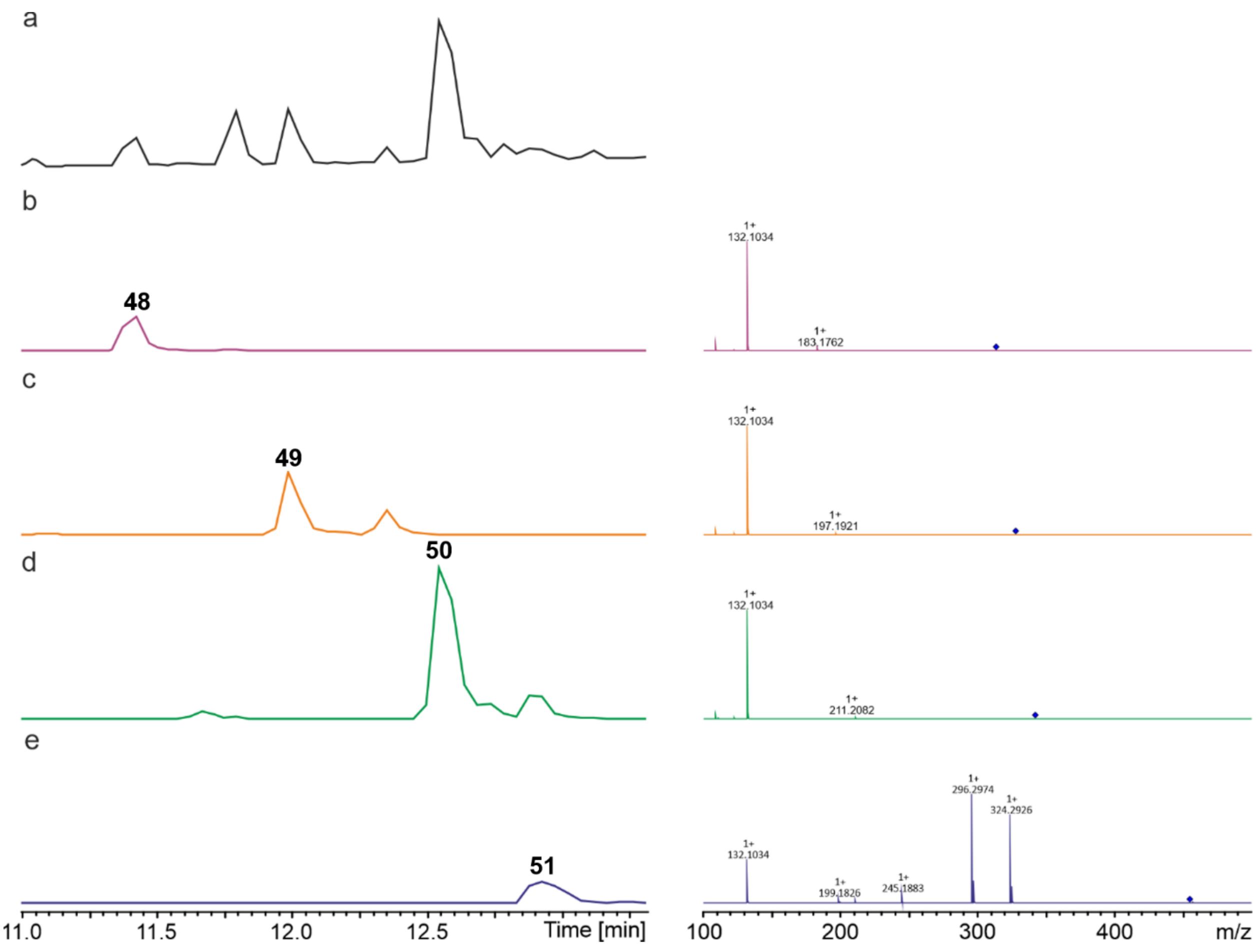

**Figure S42. HPLC/MS data refers to Supplementary Figure 18 (NRPS-39) of compounds 48, 49, 50 and 51 produced in *E. coli* DH10B::*mtaA*.** (a) BPC of an exemplary culture extract. (b) EIC/MS2 data of 48 (*m/z* $[M+H]^+$ = 314.27). (c) EIC/MS2 data of 49 (*m/z* $[M+H]^+$ = 328.29). (d) EIC/MS2 data of 50 (*m/z* $[M+H]^+$ = 342.20). (e) EIC/MS2 data of 51 (*m/z* $[M+H]^+$ = 455.38).

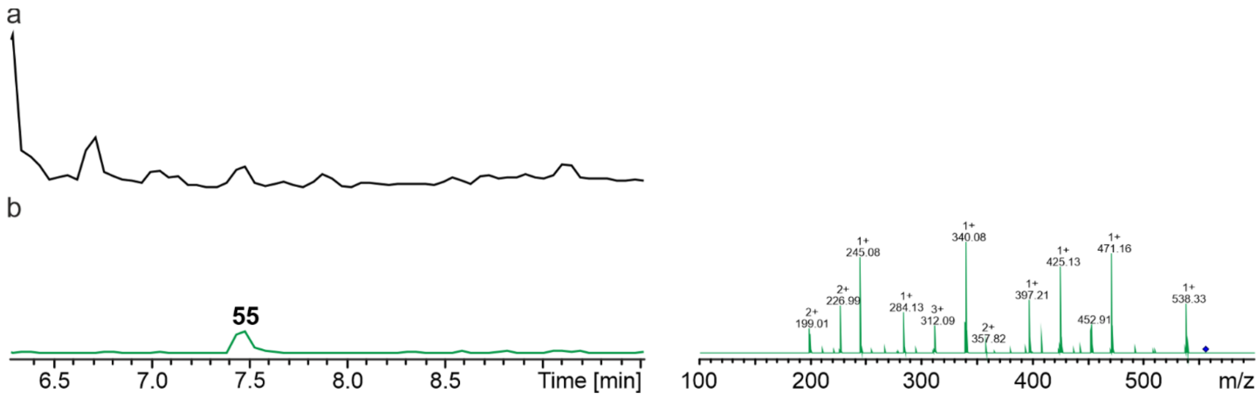

**Figure S43. HPLC/MS data refers to Supplementary Figure 18 (NRPS-40) of compound 55 produced in *E. coli* DH10B::*mtaA*.** (a) BPC of an exemplary culture extract. (b) EIC/MS$^2$ data of **55** (*m/z* $[M+H]^+$ = 556.35).

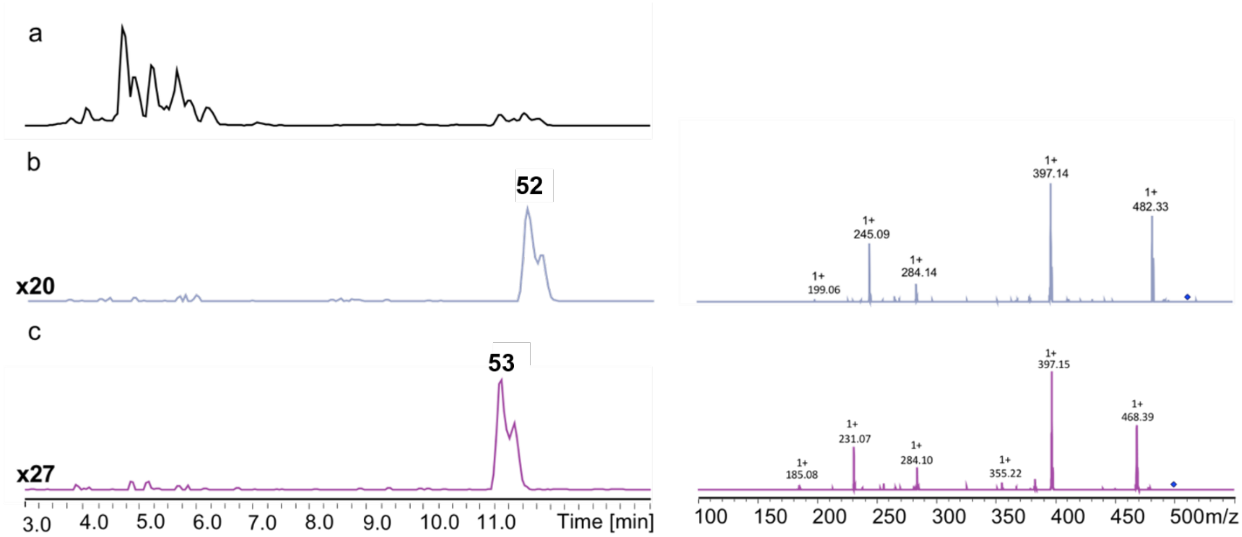

**Figure S44. HPLC/MS data refers to Supplementary Figure 19 (NRPS-43) of compounds 52 and 53 produced in *E. coli* DH10B::*mtaA*.** (a) BPC of an exemplary culture extract. (b) EIC/MS2 data of 52 (*m/z* [M+H]$^+$ = 510.39). (c) EIC/MS2 data of 53 (*m/z* [M+H]$^+$ = 496.37).

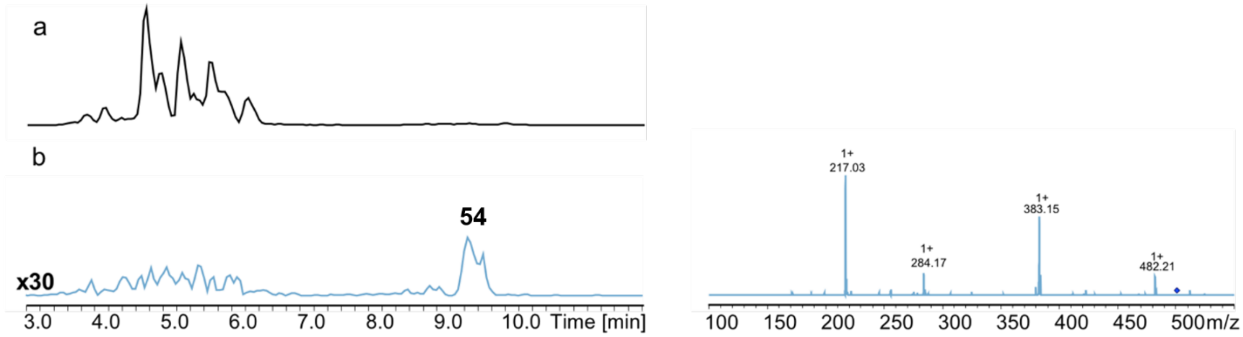

**Figure S45. HPLC/MS data refers to Supplementary Figure 19 (NRPS-44) of compound 54 produced in *E. coli* DH10B::*mtaA*.** (a) BPC of an exemplary culture extract. (b) EIC/MS2 data of 43 (*m/z* [M+H]$^+$ = 500.37).
